# Electrophile scanning by chemical proteomics reveals a potent pan-active DUB probe for investigation of deubiquitinase activity in live cells

**DOI:** 10.1101/2022.09.28.509970

**Authors:** Daniel Conole, Fangyuan Cao, Christopher W. Am Ende, Liang Xue, Sheila Kantesaria, Dahye Kang, Jun Jin, Dafydd Owen, Linda Lohr, Monica Schenone, Jaimeen D. Majmudar, Edward W. Tate

**Author notes:** Auckland Cancer Society Research Centre, Faculty of Medical and Health Sciences, University of Auckland, 85 Park Road, Grafton, Auckland, 1023, New Zealand (Daniel Conole).

## Abstract

Deubiquitinases (DUBs) are proteases that hydrolyze isopeptide bonds linking ubiquitin to protein substrates, which can lead to reduced substrate degradation through the ubiquitin proteasome system. Deregulation of DUB activity has been implicated in many disease states, including cancer, neurodegeneration and inflammation, making them potentially attractive targets for therapeutic intervention. The >100 known DUB enzymes have been classified primarily by their conserved active sites, but we are still building our understanding of their substrate profiles, localization and regulation of DUB activity in diverse contexts. Ubiquitin-derived covalent activity-based probes (ABPs) are the premier tool for DUB activity profiling, but their large recognition element impedes cellular permeability and presents an unmet need for small molecule ABPs which account for local DUB concentration, protein interactions, complexes, and organelle compartmentalization in intact cells or organisms. Here, through comprehensive warhead profiling we identify cyanopyrrolidine (CNPy) probe **IMP-2373** (**12**), a small molecule pan-DUB ABP to monitor DUB activity in physiologically relevant live cell systems. Through chemical proteomics and targeted assays we demonstrate that **IMP-2373** quantitatively engages more than 35 DUBs in live cells across a range of non-toxic concentrations, and in diverse cell lines and disease models, and we demonstrate its application to quantification of changes in intracellular DUB activity during MYC deregulation in a model of B cell lymphoma. **IMP-2373** thus offers a complementary tool to ubiquitin ABPs to monitor dynamic DUB activity in the context of disease-relevant phenotypes.

**SYNOPSIS TOC:** *Graphical Abstract:* 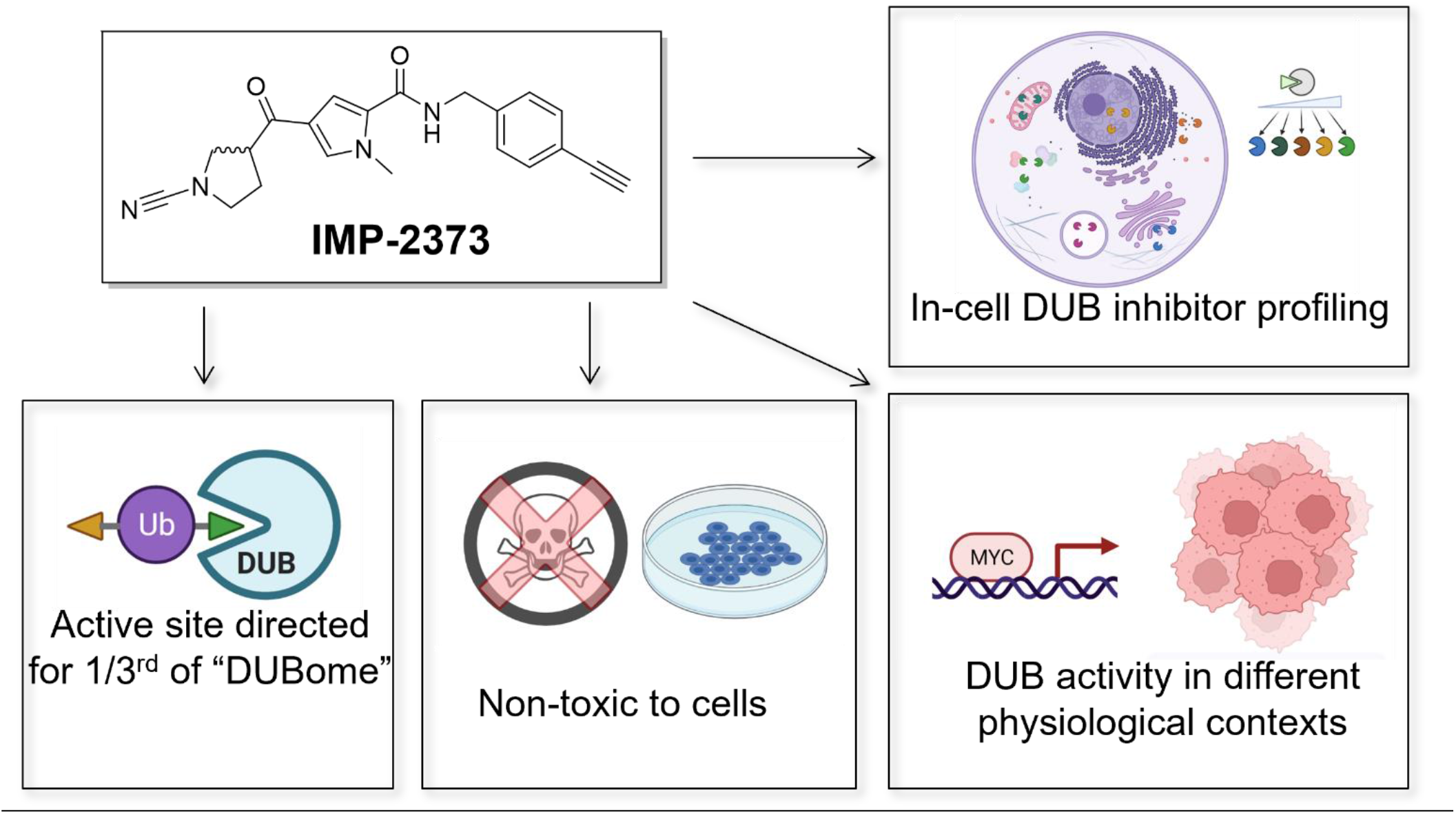

## Introduction

The ubiquitin proteasome system (UPS) regulates a myriad of intracellular processes including protein turnover, transcriptional regulation, DNA damage, protein complex formation, cellular trafficking and localization, inflammation and autophagy.^1^ The E1/E2/E3 ligase cascade appends ubiquitin (Ub) to substrate proteins, a small post-translational modification which often tags proteins for degradation at the proteasome, although E3 ligase-mediated elongation of ubiquitin chains to various branched or linear forms can lead to other functional outcomes.^2–4^ Approximately 110 individual deubiquitinase (DUB) proteases catalyze Ub hydrolysis from protein substrates or Ub chains, thereby counteracting Ub ligase activity and regulating the highly dynamic UPS. Altered DUB activity has been linked to a number of diseases and several DUBs are considered promising drug targets, with DUB inhibitors at various stages of preclinical development;^5–7^ however, target validation for DUB inhibitors has proven challenging. For example, a recent Phase I/II multiple myeloma trial of VLX1570, a putative USP14/UCHL5 covalent inhibitor, was terminated due to dose-limiting toxicity,^8^ with subsequent proteomic analyses revealing modification of a diverse range of proteins and off-target toxicity through protein aggregation.^9^ There is a pressing need for improved chemical tools and technologies to better understand DUB abundance, localization, activity and substrate profiles in health and disease, and to support development of novel, effective and selective DUB-targeted therapeutics.^10–12^ Activity-based probes (ABPs) represent a uniquely powerful tool for exploring changes in cellular DUB activity, and are based on an electrophilic warhead targeted to the DUB active site by a recognition scaffold, with DUB activity read out by a reporter group such as a dye or affinity handle.^13^ The majority of DUBs contain papain-class cysteine peptidase active sites amenable to covalent labeling by an appropriately designed ABP,^5,7^ and the first generation of DUB ABPs based on ubiquitin as the recognition element ^14^ have served to monitor DUB proteolytic activity and substrates in disease states,^15–17^ Ub chain cleavage selectivity,^18,19^ and DUB inhibitor potency and selectivity.^20,21^ Whilst the 8kDa Ub recognition element makes extensive interactions with the DUB and thereby delivers specific DUB enrichment from complex biological media, very poor cellular uptake restricts their effective use to analysis of cell lysates. The consequent loss of native organelle compartmentalization leads to dilution of DUB concentration and dissociation of protein-protein interactions (PPIs) involved in DUB activity.^22,23^ The disconnect between enzyme activity in lysates and live cells is well-recognized,^7^ and limits the capacity of Ub ABPs to profile dynamic intracellular DUB activity or its role in a particular disease state.^24^ Ub ABP uptake can be forced by high concentration and conjugation to cell-penetrating peptides (CPPs),^22^ but these complex approaches further disrupt cell membrane integrity and are not generally applicable to diverse cell lines, primary cells or animal models.^25,26^

Small molecule DUB ABPs with broad in-family DUB reactivity which passively diffuse into cells with minimal perturbation to cell physiology have the potential to complement Ub ABPs by profiling intracellular DUB activity or inhibition across many DUBs simultaneously. Two types of small molecule DUB ABP with intracellular labeling activity have been reported to date: highly targeted cyanopyrrolidine (CNPy) probes for the DUB UCHL1 (e.g. **IMP-1710**),^27–30^ and the pan-reactive chloromethyl ketone (CMK) pyrrole benzylamide probe (**4**).^31^ ABP **4** was shown to engage at least nine Ubiquitin-Specific Proteases (USPs) by proteomics, and could be used to measure USP4 activity in live osteosarcoma cells. Whilst this probe offered a promising proof of concept, it lacks pan-DUB coverage and in-class selectivity, and its limited DUB specificity leads to considerable toxicity close to concentrations useful for profiling. Here, we envisaged expanding the scope of this small molecule scaffold to engage the active DUB proteome (or DUBome), with sufficient potency and selectivity to permit activity profiling without toxicity. We designed and extensively profiled a library of thirteen probes based on this scaffold covering a diverse range of warhead reactivities and electrophile geometries, exploring intracellular protein labeling, cell viability and DUB target engagement and activity profiles. These screens led to the discovery of a next generation, pan-active, cell permeable DUB ABP **IMP-2373** (**12**) bearing a cyanopyrrolidine (CNPy) warhead that exhibits privileged DUB labeling and selectivity among all warhead classes investigated, across a range of cell lines and disease models. **IMP-2373** (**12**) represents a novel and versatile small molecule tools for probing DUB biology in complex physiological systems, which complement existing Ub ABPs.

## Results

### Chemical proteomic profiling of small molecule electrophilic warheads uncovers cyanopyrrolidine as a privileged DUB-targeting moiety

We first designed and synthesized a library of fourteen pyrrole benzylamide probes displaying diverse cysteine reactive electrophiles, including the “first generation” CMK probe **4**, selected to reflect a range of reactivity and geometric diversity, whilst maintaining synthetic tractability.^32^ Osteosarcoma (U2OS), glioblastoma (U87-MG) and breast cancer (T47D) cell lines were selected for initial probe screening experiments, as these widely-used lines together express >80% of all DUBs, as measured by mRNA profiles (Fig S1). Live cells from each cell line were incubated with each probe at 0.3 or 3 µM, and labeling analyzed by sodium dodecyl sulphate–polyacrylamide gel electrophoresis (SDS-PAGE) and in-gel fluorescence (IGF) following cell lysis and probe ligation to TAMRA via copper(I)-catalyzed azide-alkyne cycloaddition (CuAAC) (Fig. S2A). Based on these data, probes were separated into more and less reactive groups (Fig. S2B-C), and proteome-wide target engagement profiles determined by multiplexed quantitative Tandem Mass Tag (TMT) Activity-Based Protein Profiling (ABPP) at two concentrations (low and high) for more reactive warheads or a single higher concentration for less reactive warheads (Fig. 1A and B, S3). As reported previously, CMK probe **4** exhibited broad DUB target engagement with 14 DUBs enriched by log2 fold change >0.4 compared with vehicle (DMSO) (Fig 1A, S4A). Interestingly, the mildly reactive CNPy probe **12** also exhibited effective target engagement (log2 fold change >0.4 vs DMSO) for nine DUBs, with a broadly complementary profile to CMK probe **4** (Fig 1B, S4A). These data are consistent with a number of reports on CNPy covalent DUB inhibitors,^28,29,33–36^ and suggested that CNPy may be a privileged DUB ABP warhead.

**Figure 1.**
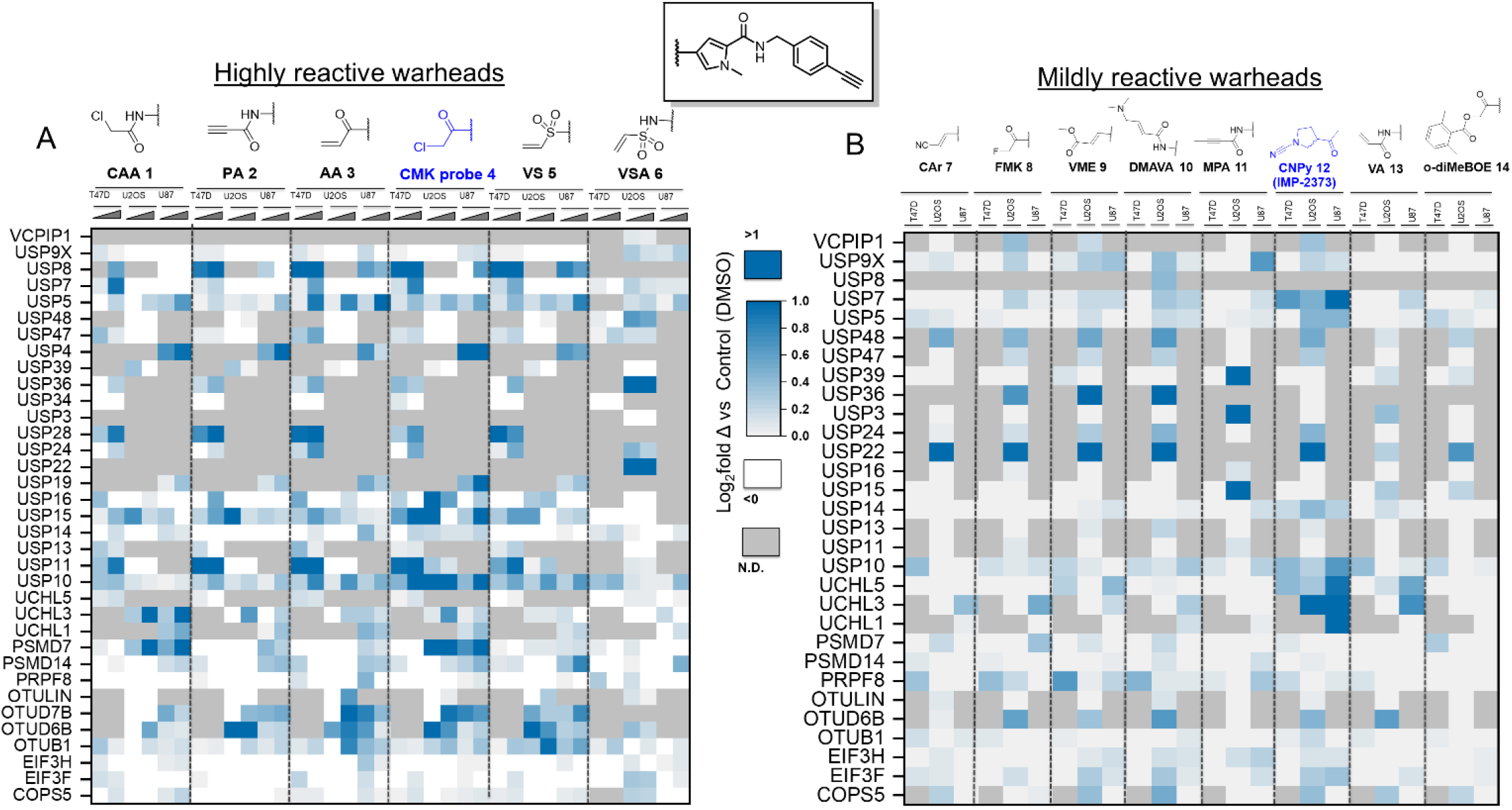
Quantitative proteomic activity-based protein profiling of a series of electrophile-armed methyl pyrroles in three cell lines identifies cyanopyrrolidine (CNPy) probe IMP-2373 (**12**) as a multi-DUB targeting probe. (A) Heat map of DUB target engagement obtained by ABPP for scaffolds harboring highly reactive electrophiles, evaluated at low (0.3 µM, 1.5 h) and high (3 µM, 1.5 h) concentrations in each of three cell lines (T47D, U2OS, U87-MG). (B) Heat map of DUB target engagement obtained by ABPP for scaffolds harboring mildly reactive electrophiles, evaluated at 3 µM (1.5 h) in three cell lines (T47D, U2OS and U87-MG).

### Cyanopyrrolidine probe IMP-2373 (12) is a potent broad-spectrum DUB enzyme inhibitor with minimal off-target cell toxicity

We next compared this indicative assessment of cellular activity and selectivity against capacity to inhibit enzyme activity against a panel of 42 recombinant DUBs (Fig. 2A). Strikingly, CNPy probe **12** displayed a markedly superior profile with respect to potency and promiscuity of DUB inhibition over all other warheads tested, similar to that observed by proteomic profiling, and complementary to that previously reported for CMK probe **4** (Fig. S4B).^31^ Despite possessing a promising DUB engagement profile (Fig 1A), **4** exhibits cytotoxic effects within a few hours at 1 µM which are likely due to the high electrophilicity and promiscuous reactivity of the CMK warhead, confounding its use as a DUB ABP which should ideally show minimal impact on cell physiology at concentrations sufficient for measurable ABP engagement (Fig. 2B, S5A-F).^31^

**Figure 2.**
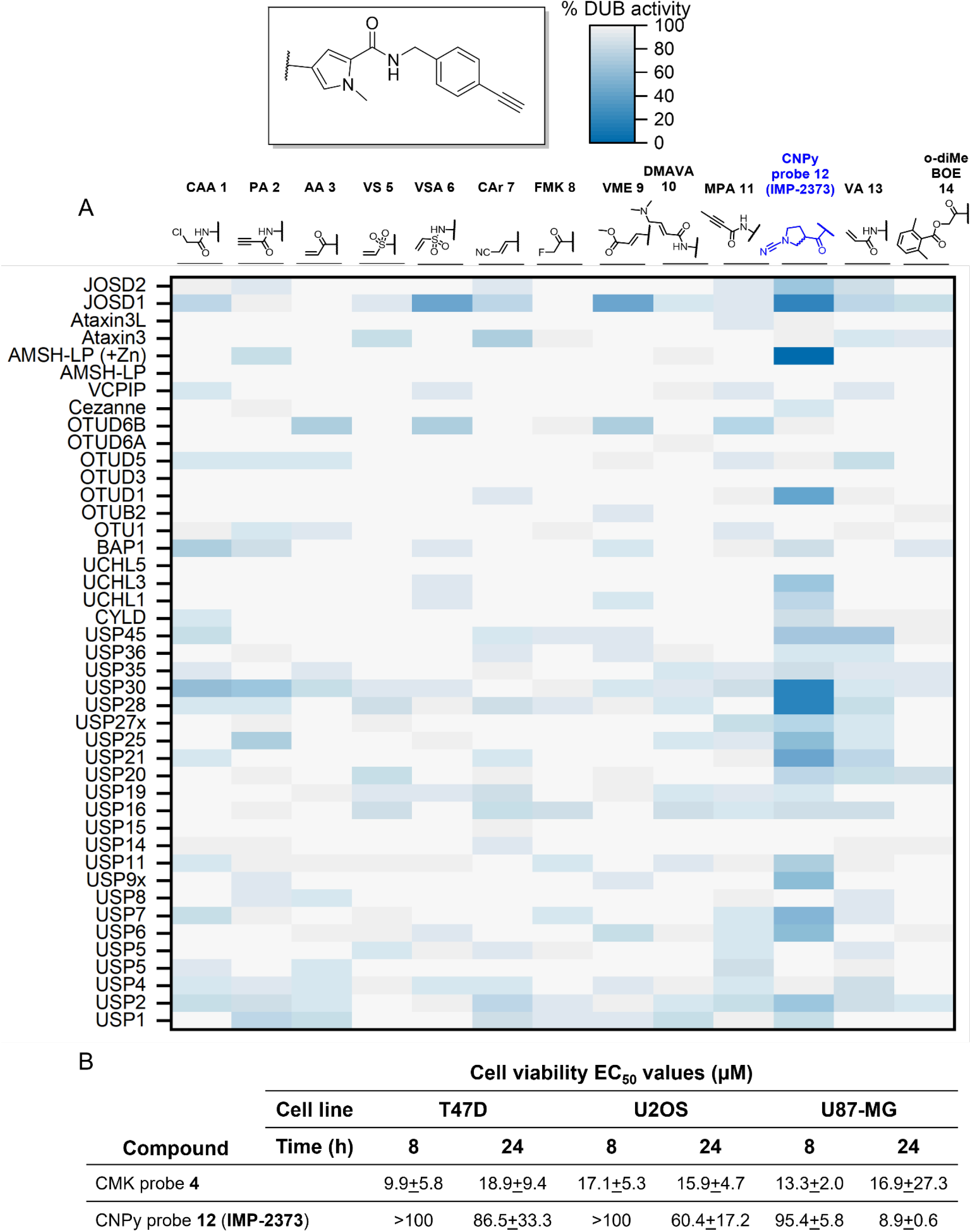
Cyanopyrrolidine ABP (CNPy probe 12) engages a wide range of DUBs in biochemical activity profiling, and shows low concentration- and time-dependent cytotoxicity. B –Cell viability measured by EthD-1 and Calcein AM dual dye cell death assay of CNPy probe 12 (IMP-2373) and CMK probe 4 in T47D, U2OS, and U87-MG cells.

Conversely, CNPy probe **12** did not affect cell viability at 8 h treatment, and remained tolerated up to 50 µM after 24 h, as measured by three distinct cell death assays (ethidium homodimer (EthD-1), Calcein AM dual dye (Fig 2B, S5A-C), and Sytox Green time-resolved cell death imaging (Fig S5D-F)), with the exception of U87-MG cells, which may be due in part to the strong dependence of gliomas on UCHL1 activity for proliferation.^37,38^ Competition experiments with a fluorescently tagged (TAMRA) Ubiquitin ABP (Ub-ABP) suggested inhibition of multiple DUBs in cell lysates, consistent with activity-based binding of CNPy probe **12** (Fig S6A), which was prioritized for further experiments and renamed **IMP-2373** to support future reference beyond the present study.

### CNPy IMP-2373 is a broad-spectrum ABP for a significant proportion of the DUBome

Encouraged by the low cytotoxicity of **IMP-2373**, we explored probe concentration and an extended incubation time (4 h) to optimize DUB engagement (Fig S6B). A total of 34 DUBs, or approximately one third of all known DUBs, were enriched log2 fold > 0.5 vs DMSO control for at least one of the concentrations tested (Fig 3A-B and D). Furthermore, enrichment of the majority of DUBs engaged by an HA (human influenza hemagglutinin peptide)-tagged ubiquitin propargyl amide (HA-Ub-PA) ABP were outcompeted by **IMP-2373** (log2 fold change < −0.5, negative for competition) (Fig. 3D).

**Figure 3.**
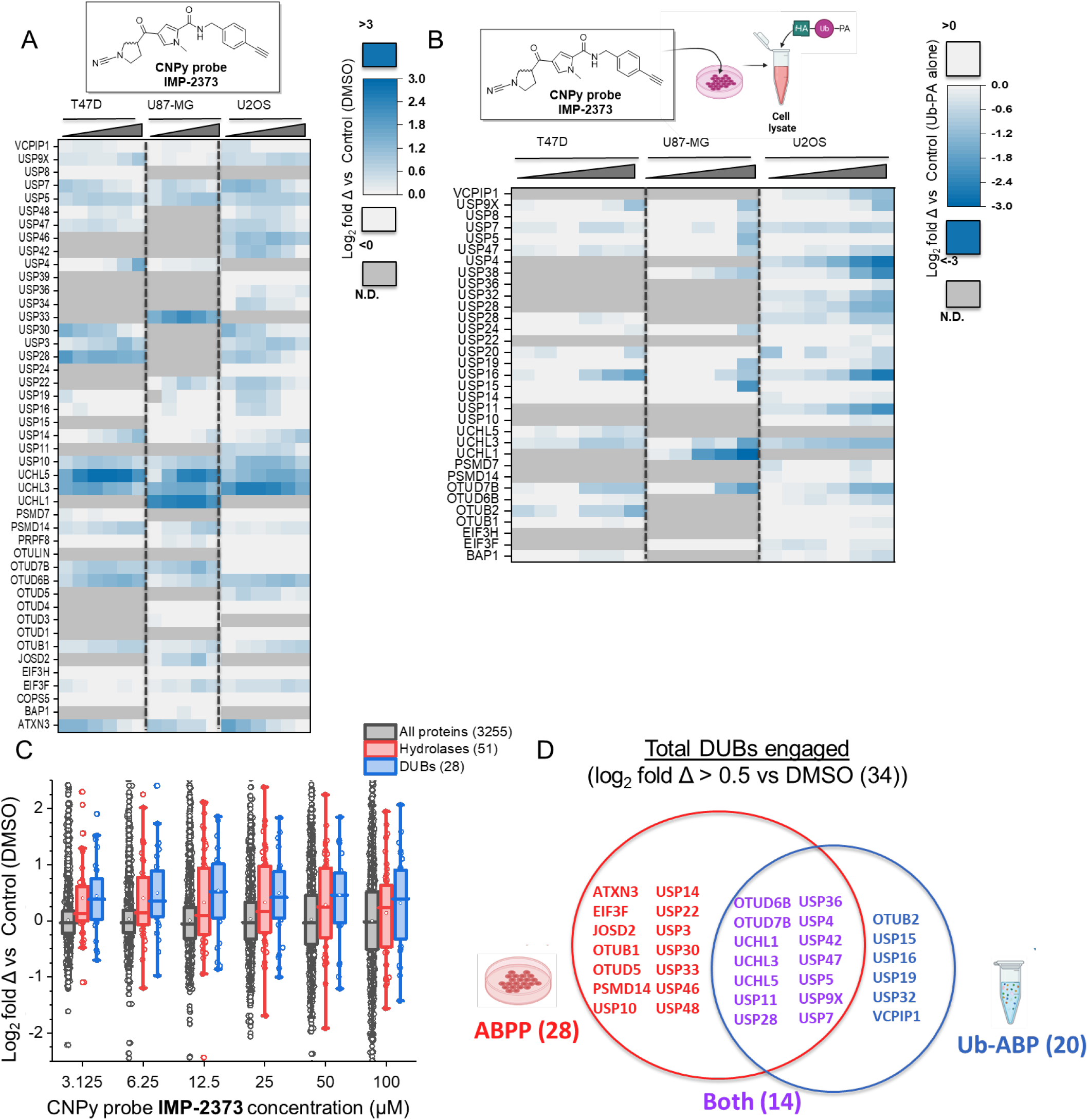
Higher CNPy probe **IMP-2373** concentrations and a longer treatment time (4 h) allows for activity-based profiling of >35% of the DUBome. Heat maps for activity-based protein profiling (ABPP) (A) and competitive ubiquitin activity-based profiling (Ub-ABP) (B) CNPy probe **IMP-2373** (concentrations: 3.125, 6.25, 12.5, 25, 50 and 100 µM, 4 h treatment, 37°C, 5% CO2) in 3 cell lines. N.D. - not detected. (C) In T47D cells, DUBs were preferentially enriched over all other proteins, and over hydrolases, the parent protease class of DUBs. (D) CNPy probe **IMP-2373** inhibits the activity–of 36 DUB (log2 fold change >0.5 vs DMSO or <-0.5 vs Ub-ABP) across all sub-classes (with the exception of ZUP1) by ABPP or Ub-ABP competition.

14 DUBs were identified as hits in both studies, consistent with activity-based engagement by **IMP-2373** at the DUB active site (Fig 3D). Statistical analysis of relative protein abundance after probe enrichment suggested that DUBs were differentially enriched, not only relative to all proteins identified but also relative to hydrolases, the parent enzyme class which encompasses DUBs (Fig. 3C). A total of approximately 70 non-DUB proteins were enriched over control for at least one of the concentrations tested, with only 8 of these non-DUB proteins conserved across the three cell lines (Fig S7A-D); gene ontology analysis determined that these off-targets are primarily peptidases (Fig S7E). Since the CNPy warhead features a stereogenic center, we were interested to know whether its enantiomers showed a preference for different DUB subclasses. We separated the enantiomers by chiral chromatography and profiled each by ABPP and Ub-ABP proteomics in T47D cells (Fig S8A). Interestingly, statistical analysis suggested that one of the enantiomers exhibited slightly more potent DUB enrichment at low concentration (0.5 µM, Fig S7B) but with no clear trend except for the UCH subfamily, consistent with previous reports on stereoselectivity in UCHL1 ABPs (Fig S8C-D).^28^

We next explored the capacity of **IMP-2373** to act as a competitive probe for in-cell target engagement by selective DUB inhibitors, a powerful and useful application of ABPs in drug discovery and development.^9,39^ Cells were pretreated for 1 h with increasing concentrations of a selective CNPy active site UCHL1 inhibitor (B1)^28^ or USP30 (FT385),^40^ followed by 10 µM **IMP-2373** for 1 h. Competitive activity-based profiling (Fig S9A-C) by pull-down and Western blot analysis confirmed potent concentration-dependent in-cell target engagement for each inhibitor (Fig. 4A, S9D-E). No competition was observed at any concentration for **IMP-1711**, the inactive enantiomer of the UHCL1 inhibitor (Fig. 4A, S9D), consistent with robust activity-based profiling.^28^

**Figure 4.**
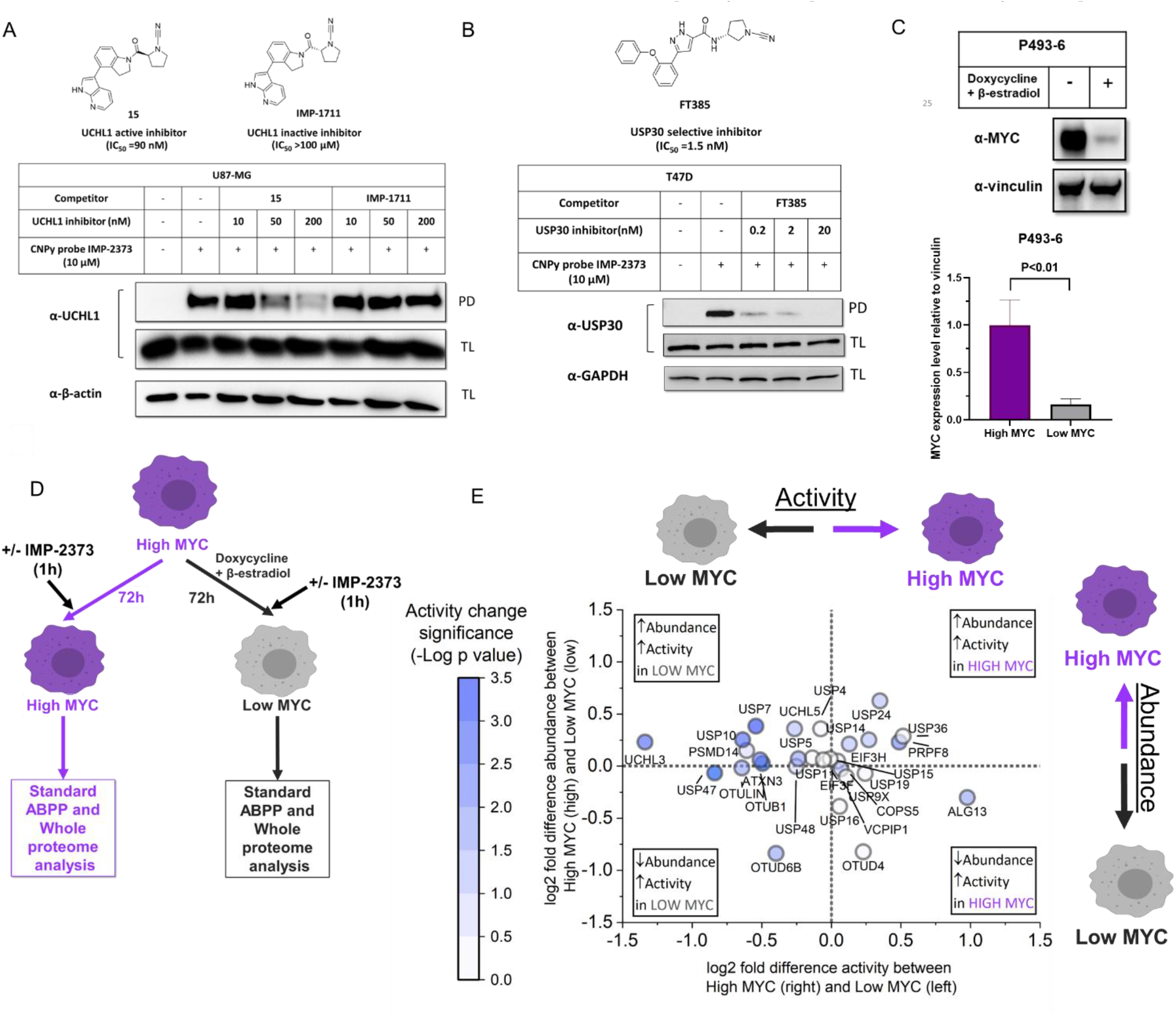
Cyanopyrrolidine ABP **IMP-2373** can be applied to profile inhibitors and to monitor DUB activity in response to differential MYC expression. Target engagement of either compound **15** (UCHL1 active inhibitor), IMP-1711 (UCHL1 inactive inhibitor) (A), or FT385 (USP30 selective inhibitor) (B) was captured by competition ABPP with CNPy probe **IMP-2373**. TL – Total Lysate; PD-Pull down. C – Western blot validation of statistically significant MYC protein level reduction in P493-6 cells in response to treatment with doxycycline and β-estradiol for 72 h. D – Experimental design to detect changes in DUB activity in response to MYC deregulation. E – Statistically significant differences in DUB activity and abundance between low and high levels of MYC, as measured by ABPP and whole proteome profiling with and without CNPy ABP **IMP-2373** (25 µM, 1 h, n = 3).

### CNPy ABP IMP-2373 enables differential DUB activity profiling during c-Myc deregulation in a model of B cell lymphoma

To demonstrate the potential of **IMP-2373** as a chemical tool to monitor changes in DUB activity in disease models, we turned to a widely-used model of MYC deregulation in cancer.^41^ MYC is a multifunctional transcription factor which regulates expression of a large number of genes involved in cellular growth, proliferation and metabolism.^42^ MYC deregulation can lead to dramatically altered protein synthesis by driving massive increases in gene transcription, and enhanced production of ribosomes and translation initiation factors,^43^ promoting cell growth, cell cycle progression, and genome instability, ultimately leading to oncogenesis and malignant tumor growth. Aberrant MYC is an oncogenic driver in > 50% of human cancers, and the mechanisms by which MYC-deregulated cancers cope with radically altered protein turnover may present novel therapeutic targets.^42^ Attempts to understand MYC-deregulated cancers are complicated by the fact that it is an essential gene for many normal cellular processes, and tightly regulated by many context-dependent post-translational modifications, with MYC overexpression typically leading to apoptosis in non-cancerous cells.^41^

Multiple components of the UPS have been proposed to regulate MYC ubiquitination and stability including both E3 Ub ligases (FBXL14,^43^ UBR5,^42,44^ FBW7^45^) and DUBs (USP13,^43^ USP29,^46^ OTUB1^47^). Here, we applied **IMP-2373** to test the hypothesis that MYC deregulation drives dynamic changes in DUB activity as part of the adaptation of cancer cells to increased protein synthesis, employing a human lymphoblastoid B cell line, P493-6, in which conditional MYC expression is under the control of an inducible promoter.^41^ P493-6 cells constitutively express c-MYC (high-MYC), however addition of doxycycline and β-estradiol for 72 h potently downregulates expression, resulting in a low-MYC state (Fig 4C, S10). Cells in each state were exposed to a short treatment with **IMP-2373**, (25 µM, 1 h). Subsequently, whole proteome and activity-based proteomic analyses were undertaken to enable differential quantification of both overall protein expression and probe labeling in high- vs low-MYC cells, for each probe treated or untreated control condition (Fig 4D). **IMP-2373** captured the activity of 38 DUBs across MYC high and low cells (Fig S11 A-D), and this broad coverage of DUBs by both ABPP and quantitative whole proteome analysis allowed us to make direct comparisons between DUB abundance and activity across high and low MYC cell lines.

Whilst the abundance of 33 DUBs showed only small changes between MYC states, global DUB activity was markedly lower in high-MYC cells (Fig S11E-F, S12A), and there was no overall correlation observed between changes in DUB activity and abundance (Fig S12B). Statistically significant upregulation of activity relative to abundance was observed in low-MYC cells for UCHL3, USP7, USP47, USP10 and ATXN3, offering the first insights into MYC-dependent differential regulation of DUB activity in intact cells (Fig 4E, S11F).

## Discussion

ABPs are well-established for certain enzyme classes, such as fluorophosphonate ABPs for serine hydrolases.^48,49^ A similar “in-family pan-reactive” ABP for DUBs would enable detection of changes in enzyme activity in response to specific cellular perturbations, and elucidation of how DUBs drive particular cellular phenotypes.^50^ In this work, we introduce CNPy probe **IMP-2373** as a scalable and versatile in-cell DUB ABP, with applications including inhibitor profiling and as a tool to study the impact of changes in biological context on DUB activity.

Compared with those of Ub-ABP (Fig 3B), experiments with probe alone and ABPP (Fig 3A) resulted in varied and in some cases counter-intuitive “reverse” dose/response data. The cause of this phenomenon remains unclear, but it may be in part due to promotion of DUB ubiquitination and degradation, of triggered by high target occupancy, or probe-activated cellular stress pathways.^51^ Nonetheless, it suggests caution should be exercised when selecting the concentration and incubation times for various in-cell probe applications. This comprehensive study suggests that that the working concentration range of the probe should not exceed 50 µM, and that a one hour treatment with *ca*. 10-25 µM probe should be sufficient to capture most DUB activities. However lower concentrations may still permit the sensitive capture of specific DUB subsets, such as the UCHL family.

To our knowledge, our ABPP experiments using **IMP-2373** in intact B cell lymphoma cells provide the first evidence for dynamic changes in DUB activity, rather than simple abundance, due to MYC deregulation in cancer. Interestingly, our results also suggest that MYC deregulation may provoke downregulation of multiple DUB activities, which we hypothesize may be an adaptation to the increased rate of protein translation in these cells,^43^ tending to retain ubiquitination, protein degradation and protein turnover. In future it may be interesting to examine the potentially cytotoxic consequence of activation of UCHL3, USP7, USP47, USP10 or ATXN3 in MYC-deregulated cancers, and to investigate the substrate profiles of these enzymes in MYC-deregulated cells. Further to this, we suggest that **IMP-2373** could be applied to understand the activity of DUBs in other, diverse contexts. For example, future studies could explore the impact of other pathological and physiological processes causing rapid changes in protein turnover, such as the switch to cellular quiescence^52^ or senescence in cancer,^53,54^ UPS-independent but Ub-dependent degradation pathways such as autophagy or mitophagy,^55^ or host-pathogen interactions, for example during infection by viruses which hijack endogenous DUB activity.^56^

DUB and Ub substrate selectivity is typically driven by extensive macromolecular interactions, presenting a significant challenge for small molecule DUB probe design which must retain small size to avoid cell impermeability, as is the case for 8 kDa Ub ABPs. Covalent capture of the DUB active enzyme site with an electrophilic warhead offers a potentially powerful approach for potent and selective DUB inhibition, however challenges remain in tuning such warheads towards DUBs. Several studies point to cyanopyrrolidines as a privileged warhead for DUBs, with enhanced reactivity toward the DUB active site relative to other classes of protease.^28,29,33–36^ CNPy probe **IMP-2373** exhibits a clear preference for DUBs over other proteins in general, and even within the hydrolase class, resulting in greatly reduced cytotoxicity compared to previous designs. However, whilst labeling can be readily achieved at sub-toxic concentrations in multiple cell lines, **IMP-2373** retains residual labeling activity at some off-target proteins (Fig S6E) which may risk perturbing cellular phenotype. To address this limitation, future efforts could mine the rapidly growing biochemical CNPy DUB inhibitor patent literature^33,57,66–69,58–65^ and combine these reported biochemical activities with systematic docking analysis across all known and predicted human DUB structures (for example, as derived by AlphaFold)^70^ to optimize the scaffold attached to the CNPy warhead for DUBs over off-target proteins, or for a particular DUB subfamily. In conjunction with conventional structure-based design and optimization, this approach may eventually permit discovery of more potent and selective DUB-privileged small molecules and ABPs for future fundamental biology and therapeutic applications.

## Conclusion

By exploring a diverse range of electrophiles attached to a constant 4-methylpyrrole benzylamide scaffold, we have identified cyanopyrrolidine as a privileged warhead for broad spectrum targeting of DUB enzymes. CNPy probe **IMP-2373** represents the most potent and selective pan-DUB small molecule ABP reported to date, and permits profiling of DUB activities and DUB inhibitors in intact cells, providing a useful complement to Ub ABPs in studies to uncover regulation of DUB activity.

## Supporting information

Supplementary information 3 - 1H NMR spectra of chemical probes

## ASSOCIATED CONTENT

**Supporting Information 1**. Supplementary figures for additional experimental results (Figures S1-12) .pdf

**Supplementary Information 2**. Additional experimental details, materials, and methods .pdf

**Supplementary Information 3**. ^1^H NMR spectra of final chemical probes.pdf

This material is available free of charge via the Internet at http://pubs.acs.org.”

The mass spectrometry proteomics data have been deposited to the ProteomeXchange Consortium via the PRIDE^71^ partner repository with the dataset identifier PXD035417. Reviewer account details are as follows:

**Password:** V4mSEeOH

## AUTHOR INFORMATION

### Author Contributions

The manuscript was written through contributions of all authors. All authors have given approval to the final version of the manuscript.

### Funding Sources

This work was supported by Pfizer Inc.

## ACKNOWLEDGMENT

The authors thank L. Haigh (Department of Chemistry Mass Spectrometry Facility, Imperial College London) for assistance in acquiring nanoLC-MS/MS data, and Linxin Wu, Renyuan Hong, Youzhen Wu and Edelweiss Evrard for their synthetic contributions and outsourcing support. This study was supported by Pfizer Inc.

## ABBREVIATIONS

DUB: Deubiquitinase
Ub: ubiquitin
UPS: ubiquitin proteasome system
ABP: activity-based probe
Ub-ABP: Ub-activity-based probe
CNPy: cyanopyrrolidine

## CONFLICTS OF INTEREST

Jaimeen D. Majmudar, Christopher Am Ende, Dafydd Owen, Monica Schenone, Dahye Kang, Liang Xue, Sheila Kantesaria and Linda Lohr are employees of Pfizer. Edward W. Tate is a director and shareholder of Myricx Pharma Ltd.

