## Supplementary information 1 - supplementary figures for "Electrophile scanning by chemical proteomics reveals a potent pan-active DUB probe for investigation of deubiquitinase activity in live cells"

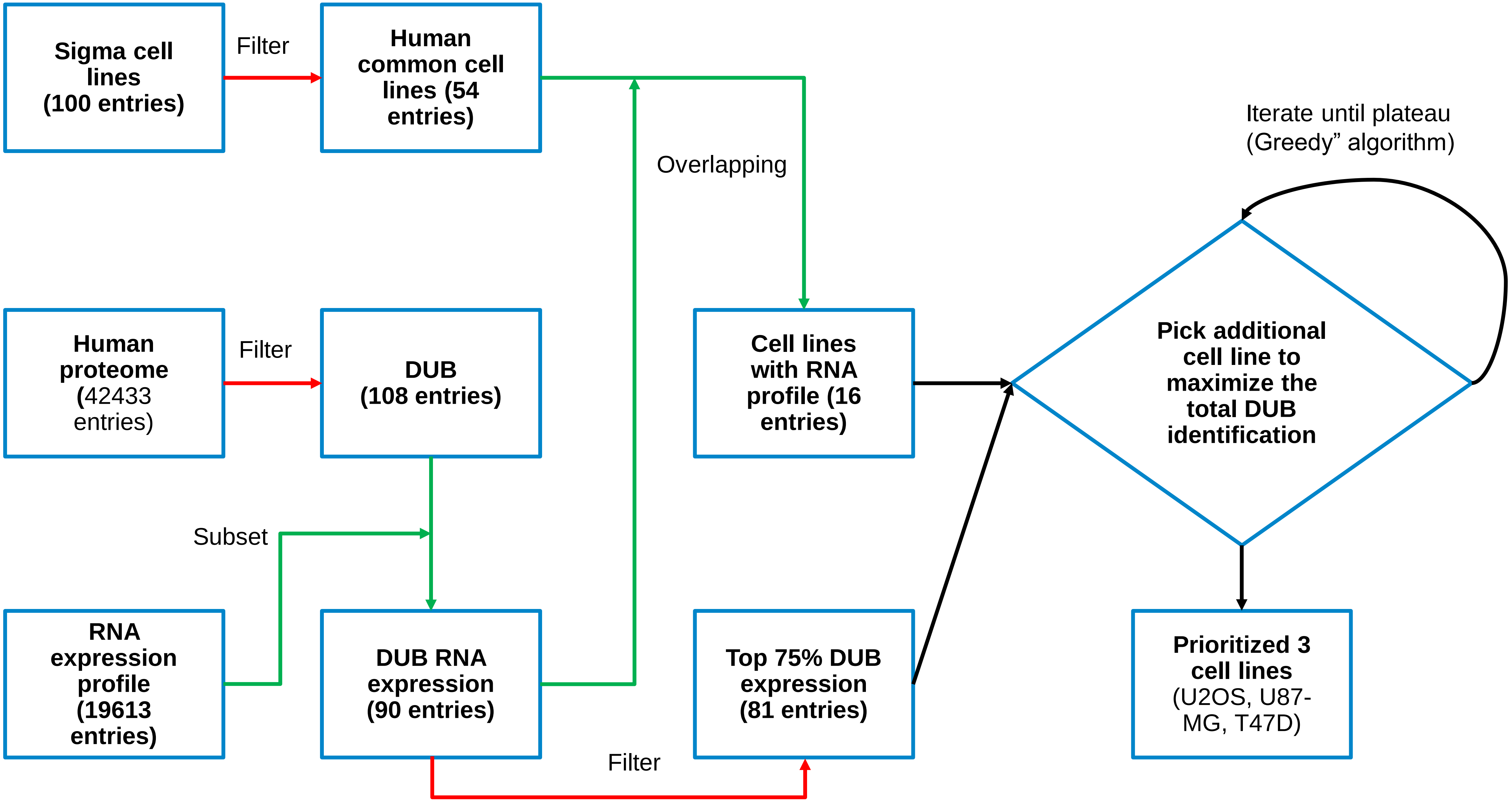


**Figure S1.** Computational workflow to select appropriate cell lines for DUB activity profiling. Cell lines were selected on the basis of commercial availability/accessibility and their DUB RNA expression profiles. The final number of cell lines (n =3) for maximal DUB coverage was determined using the “Greedy” algorithm.

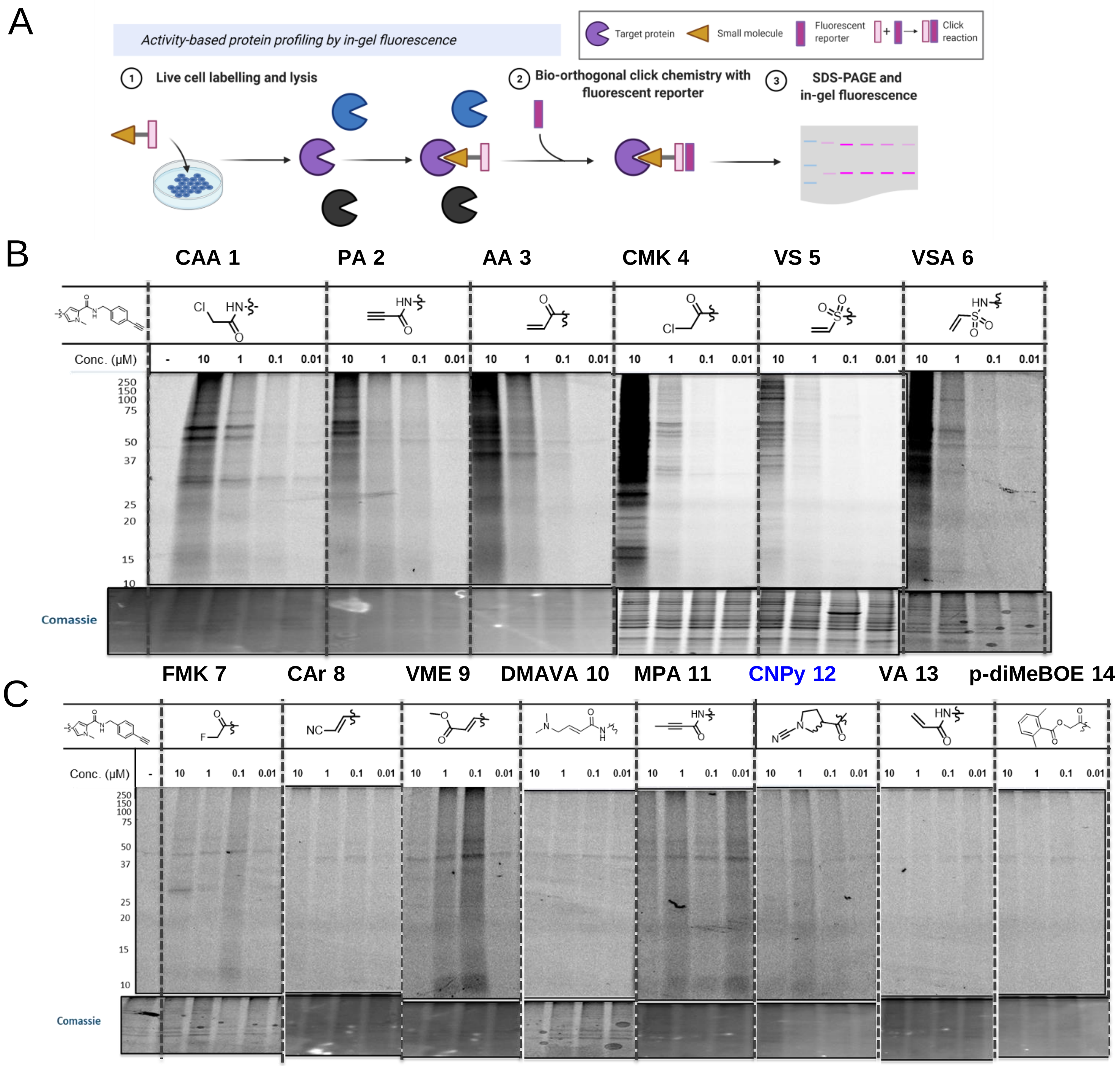


**Figure S2. A –** In-gel fluorescence workflow. Live cell labelling with probes, followed by cell lysis , biorthogonal CuAAC click chemistry, SDS-PAGE and in-gel fluorescence imaging. **B -** Electrophilic warheads (probes **1-6**) that demonstrated concentration dependent labelling by in-gel fluorescence **C -** Electrophilic warheads (probes **7-14**) that did not demonstrate concentration dependent labelling by in-gel fluorescence, up to 10µM. Panels B and C show representative in-gel fluorescence images for U2OS cell line only, however the results were consistent for T47D and U87-MG (data not shown).

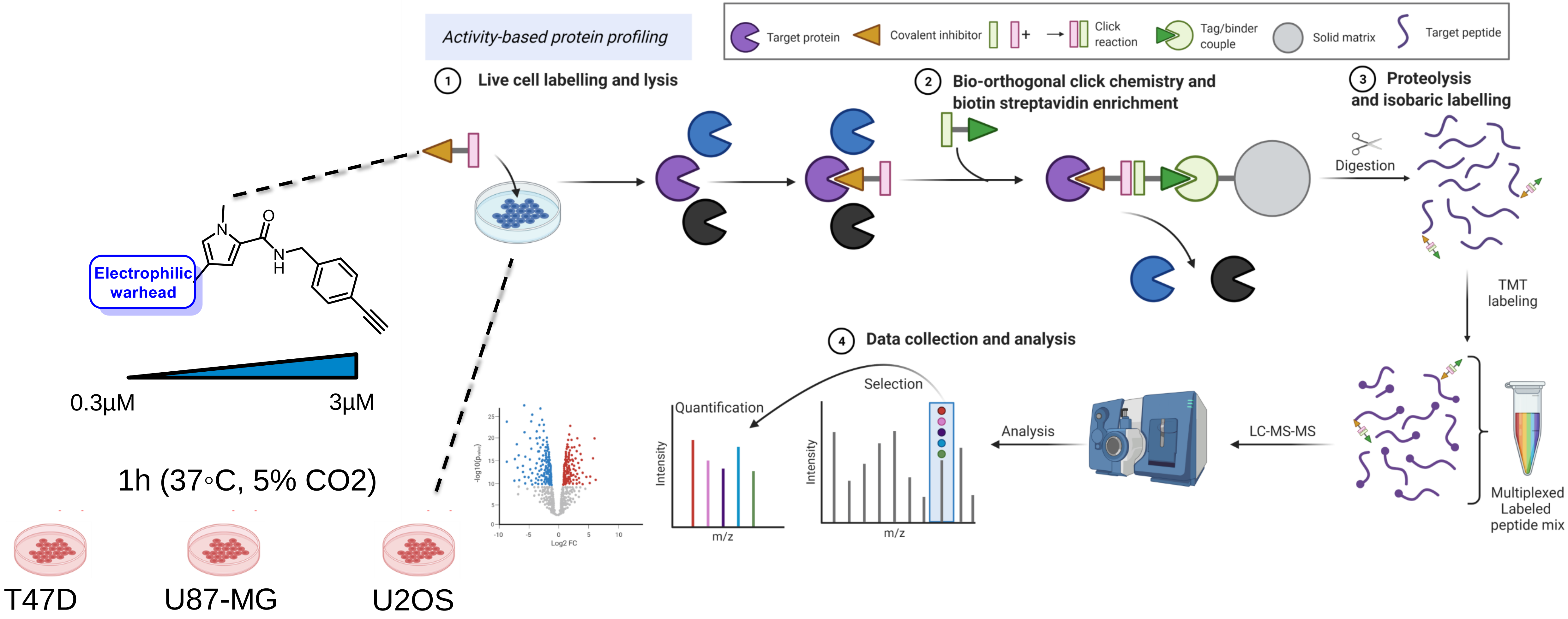


**Figure S3. A typical Activity-based protein profiling (ABPP) workflow.** Live cell labelling with covalent probes **1-14** is followed by cell lysis, click chemistry, avidin enrichment, on-bead trypsin digestion, isobaric Tandem Mass Tag (TMT) labelling and LC-MS/MS analysis.

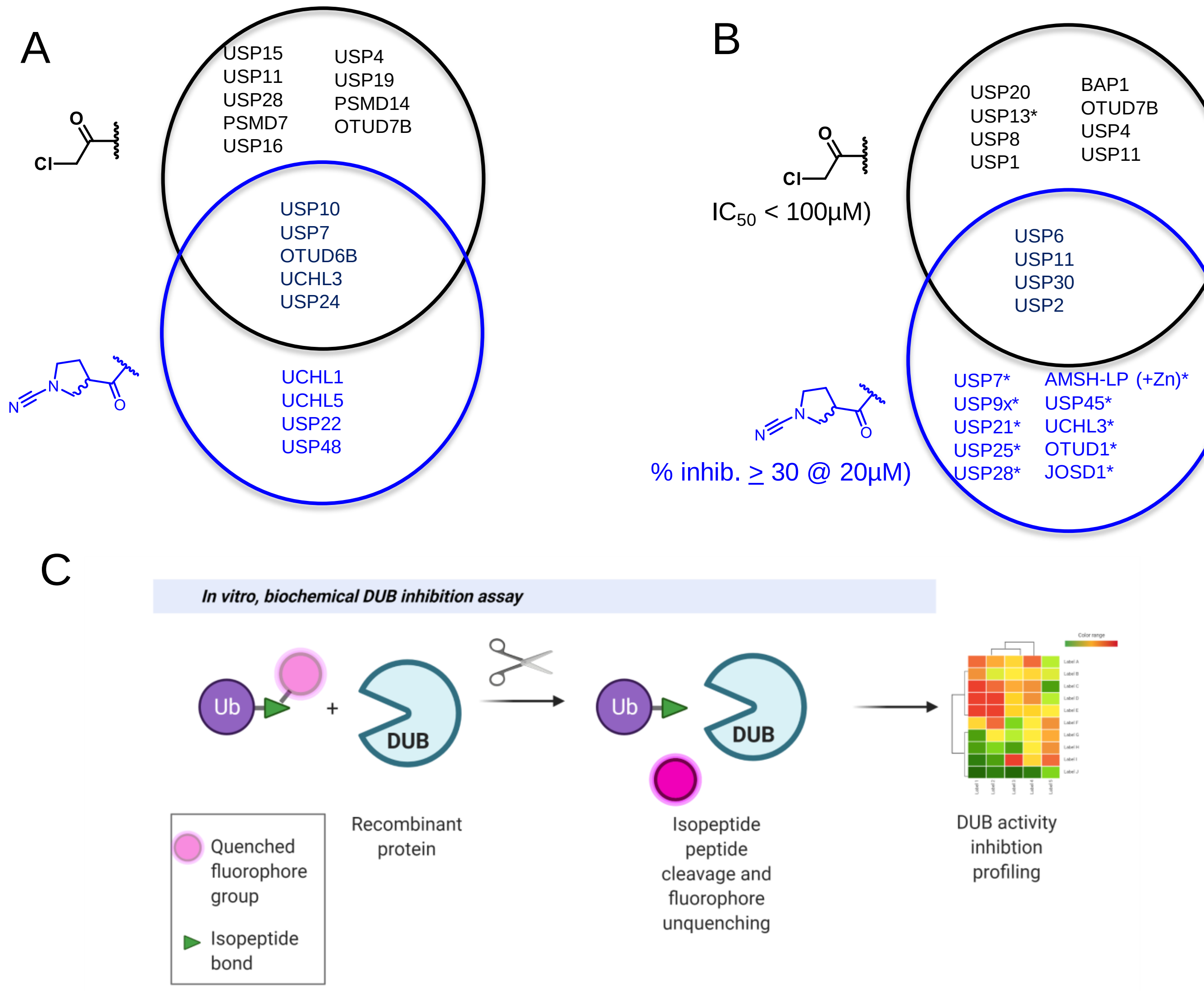


**Figure S4. A-** Coverage of DUBs engaged by $\log_2$-fold $\Delta > 0.4$ by CMK probe **4** and CNPy probe **12** (**IMP-2373**). **B –** Hits obtained from DUB panel biochemical inhibition selectivity screening for 1st generation warhead (CMK probe **4**) vs 2nd generation warhead (CNPy probe **12** (**IMP-2373**)). Screens for both ABPs run employing ubiquitin-rhodamine(110)-glycine substrate based-assay, however these assays were run at different times by different companies. The asterisk denotes the absence of this DUB from the screen against the other ABP. **C –** Schematic of the Rhodamine-110 in vitro biochemical DUB inhibition fluorogenic assay.

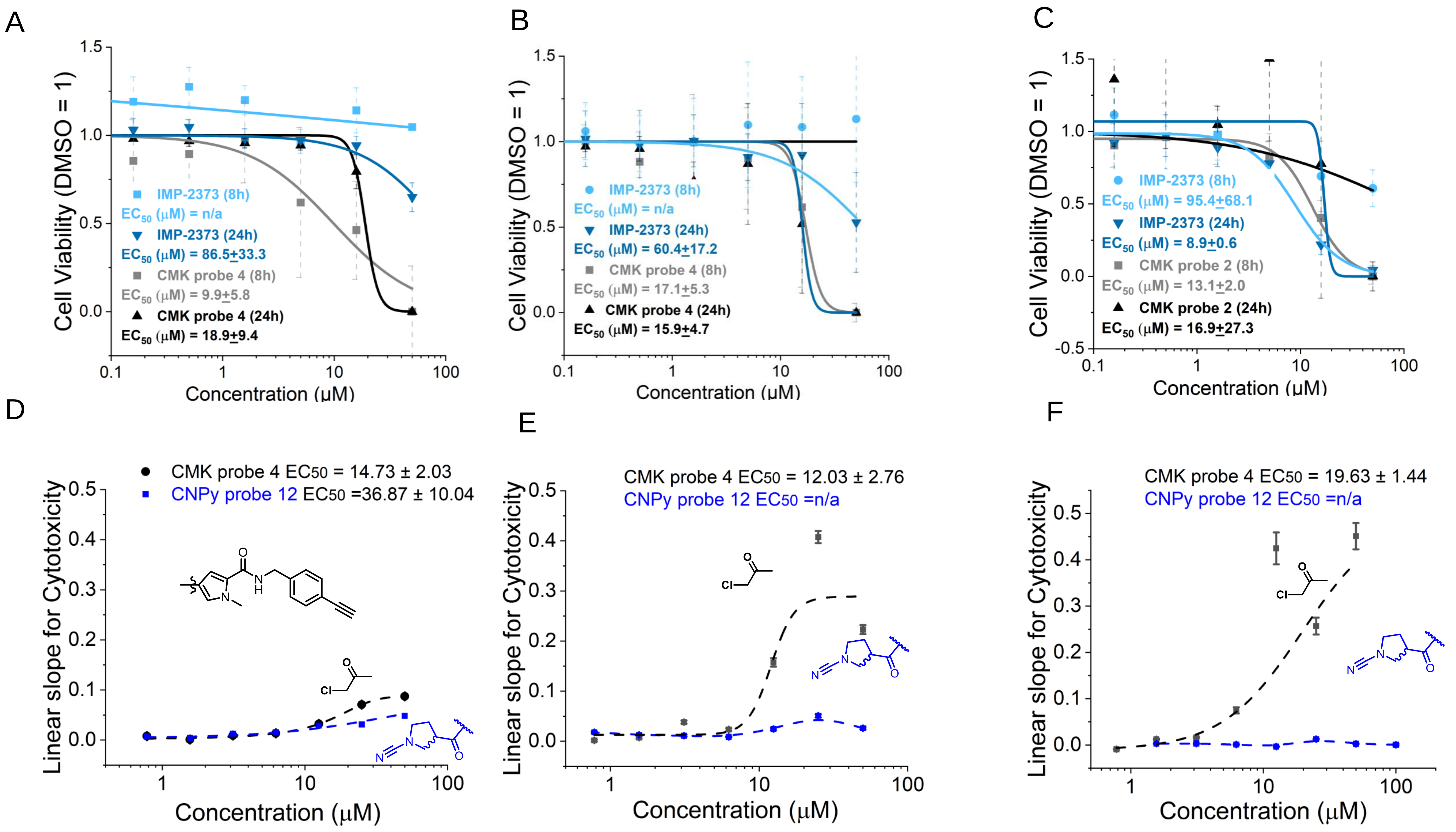


**Figure S5.** Cyanopyrrolidine ABP (CNPy probe **12**) shows low concentration- and time-dependent cytotoxicity. Cell viability measured by EthD-1 and Calcein AM dual dye cell death assay of CNPy probe 12 (IMP-2373) and CMK probe 4 in T47D (**A**), U2OS (**B**), and U87-MG (**C**) cells. Results for CMK probe **4** and CNPy probe **12** (**IMP-2373**) in a time-resolved (over 24h) cell death imaging assay. The greater the linear slope the greater the cytotoxicity of the probe. CNPy probe **12** (**IMP-2373**) is largely non-toxic up to 50uM after 24h in U87-MG (**D**), U2OS (**E**) and T47D (**F**).

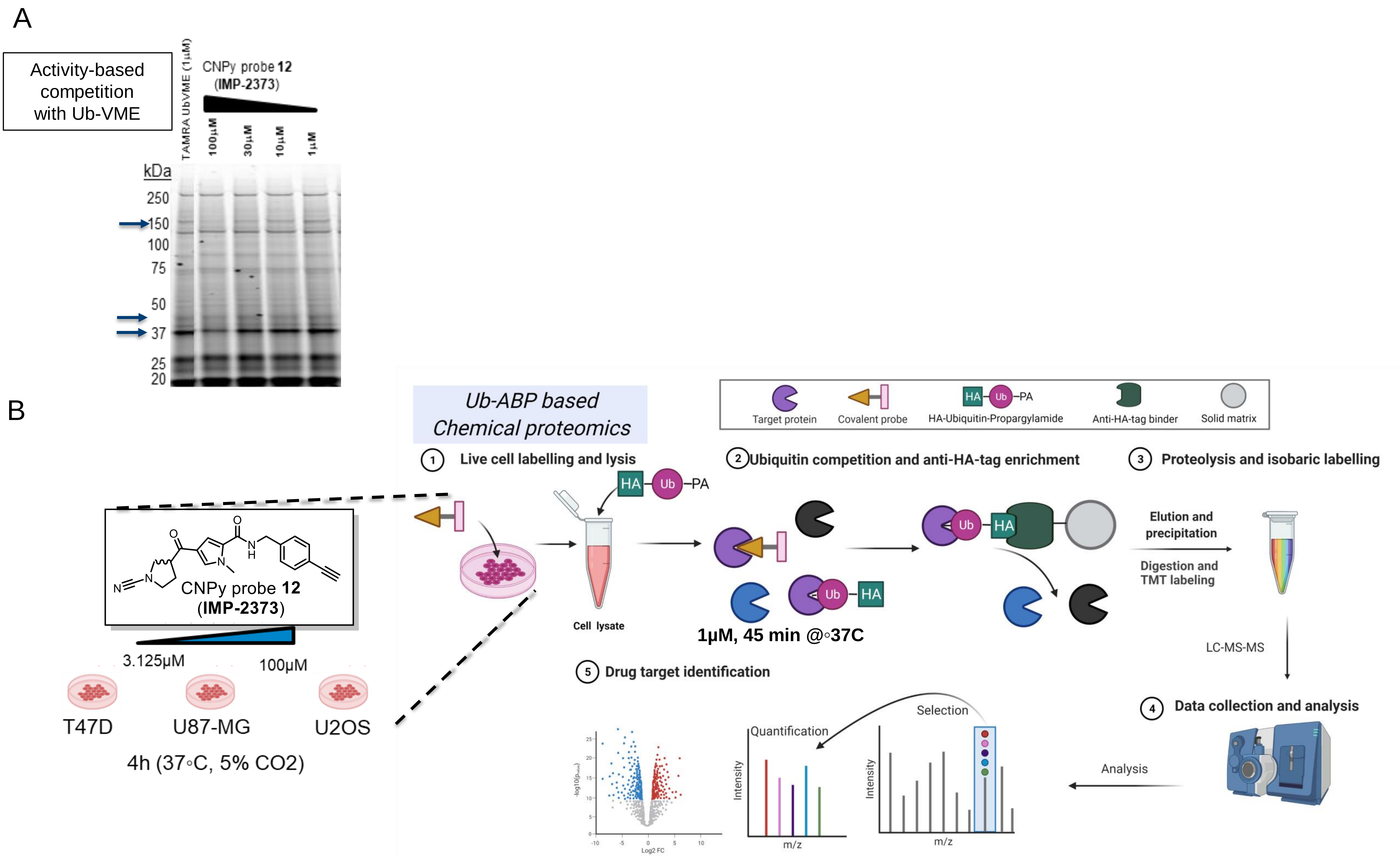


**Figure S6. A –** Ubiquitin ABP competition profiling of CNPy probe **12** in HEK293 lysates; 1 h probe treatment followed by 1 h treatment with TAMRA-Ub-VME (arrows indicate competition of CNPy probe **12** with Ub-ABP for DUB active site engagement). **B -** Typical workflow for Ubiquitin activity-based proteomics, with an extended incubation time (4 h, instead of 1.5 h) and increased concentrations for CNPy probe **12** (**IMP-2373**) . ABPP workflow was as outlined in Figure S2, except with incubation times and increased concentrations for CNPy probe **12** (**IMP-2373**) .

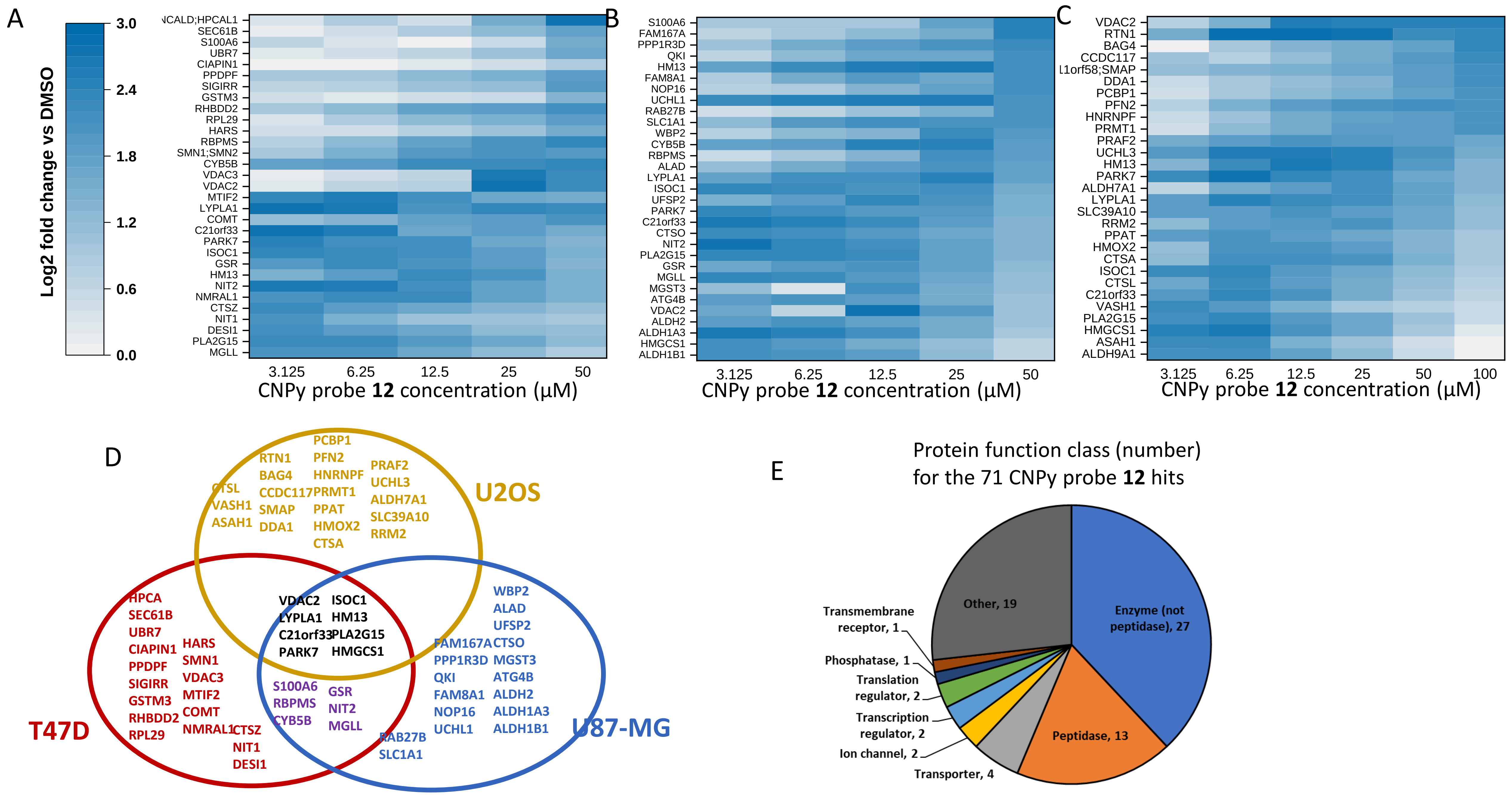


**Figure S7.** Top proteins engaged (log$_2$ fold change >2 @ at least one concentration (n~30 per cell line) by CNPy probe **12** (**IMP-2373**) in T47D (**A**) U87-MG (**B**) and U2OS (**C**). **D -** Summary of conserved CNPy probe **12** (**IMP-2373**) protein targets. **E –** Summary of function classification for top proteins engaged by CNPy probe **12** (**IMP-2373**) (generated by Ingenuity Pathway Analysis).

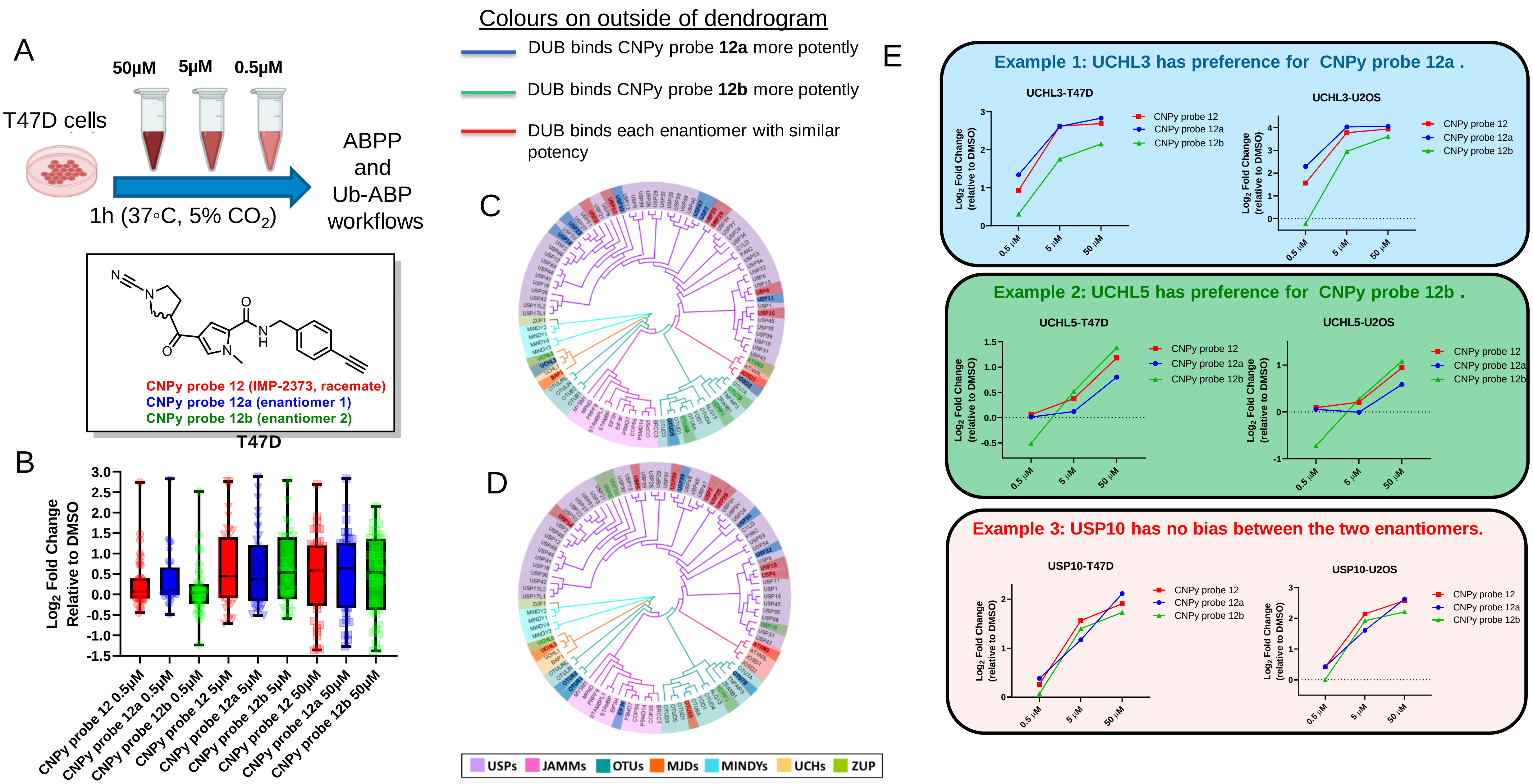


**Figure S8. A –** Chemical structure for CNPy **12** racemate (and compound codes for **enantiomers 12a** and **12b**) and live cell incubation times and probe concentrations employed in activity-based experiments. **B** – Distribution of enriched DUB abundance of CNPy racemate (CNPy probe **12**) and enantiomers (**probe 12a** and **12b**) vs DMSO treated T47D cells. Annotated DUB family tree dendrogram annotated with preferences of DUBs for engagement ($log_2$ fold Δ >0.5 vs DMSO) with the different CNPy probe enantiomers (**probe 12a** and **12b**) from ABPP (**C**) and Ub-ABP (**D**) experiments in live T47D cells. **E –** Selected examples of enantiomer ABP target engagement profiles that exhibit selectivity/bias for individual DUBs.

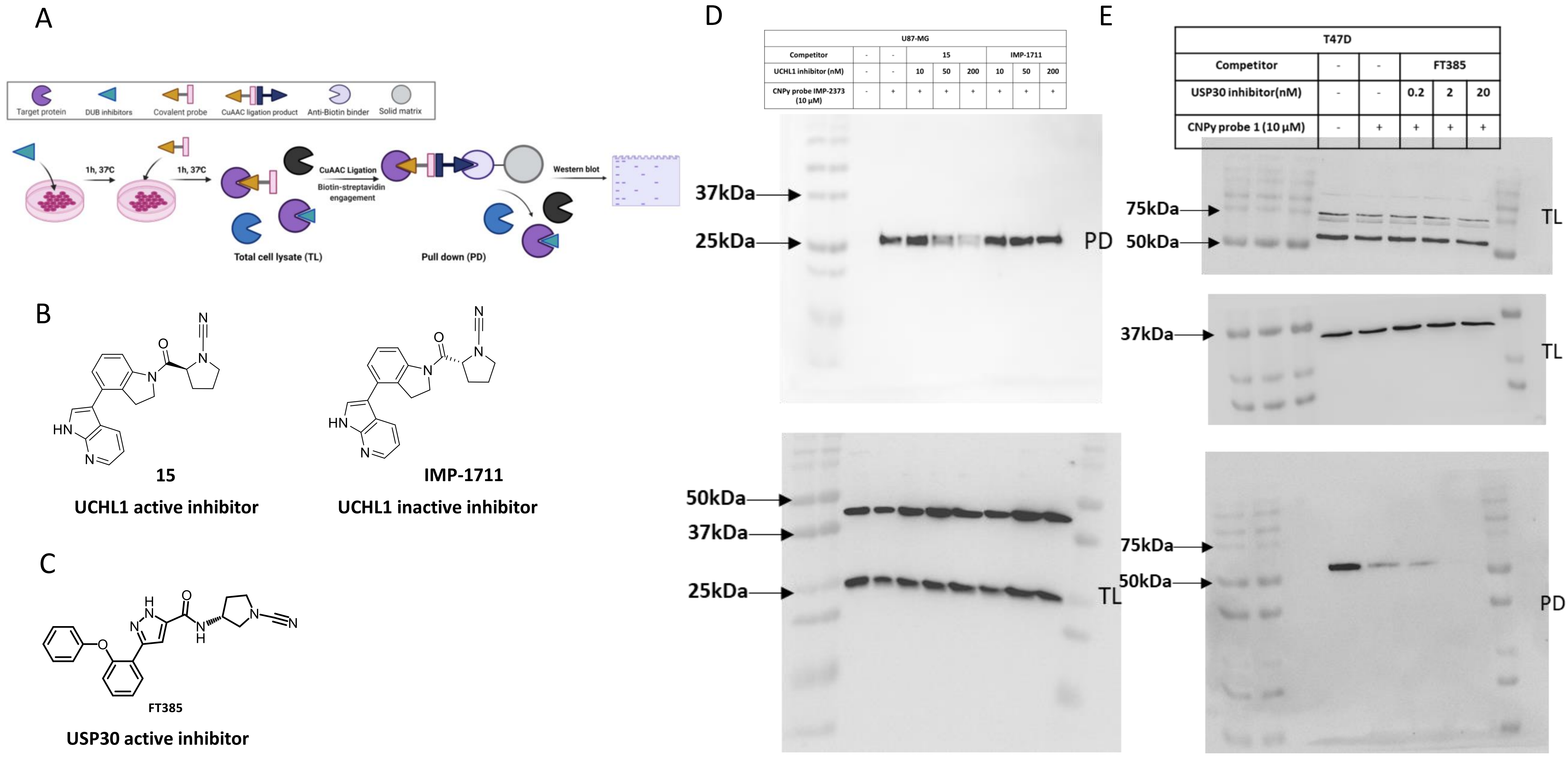


| U87-MG | | | | | | | | |
|---|---|---|---|---|---|---|---|---|
| Competitor | - | - | 15 | | | IMP-1711 | | |
| UCHL1 inhibitor (nM) | - | - | 10 | 50 | 200 | 10 | 50 | 200 |
| CNPy probe IMP-2373 (10 µM) | - | + | + | + | + | + | + | + |

| T47D | | | | | |
|---|---|---|---|---|---|
| Competitor | - | - | FT385 | | |
| USP30 inhibitor(nM) | - | - | 0.2 | 2 | 20 |
| CNPy probe 1 (10 µM) | - | + | + | + | + |

**Figure S9. A –** Competition-based DUB inhibitor profiling workflow with CNPy probe **IMP-2373**. Active site inhibition was quantified by western blot for selected DUBs (UCHL1 and USP30). Chemical structures for the UCHL1 (**B**) and USP30 (**C**) selective DUB inhibitors employed in these experiments. Uncropped blot for UCHL1 (**D**) and USP30 (**E**) inhibitor profiling experiments. TL – Total Lysate; PD – Pull-down.

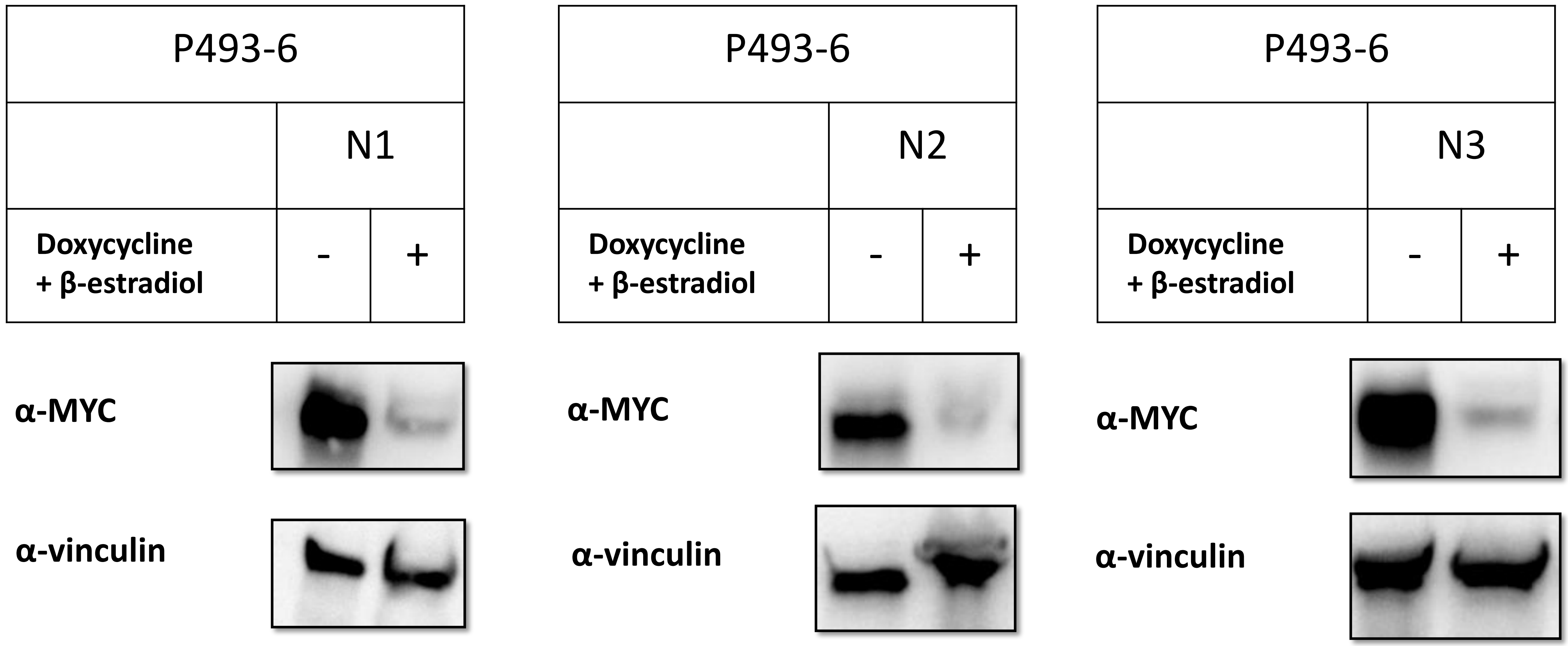


**Figure S10.** Western blot validation of MYC protein level reduction in P493-6 cells in response to 3 independent treatments with doxycycline and β-estradiol for 72 h.

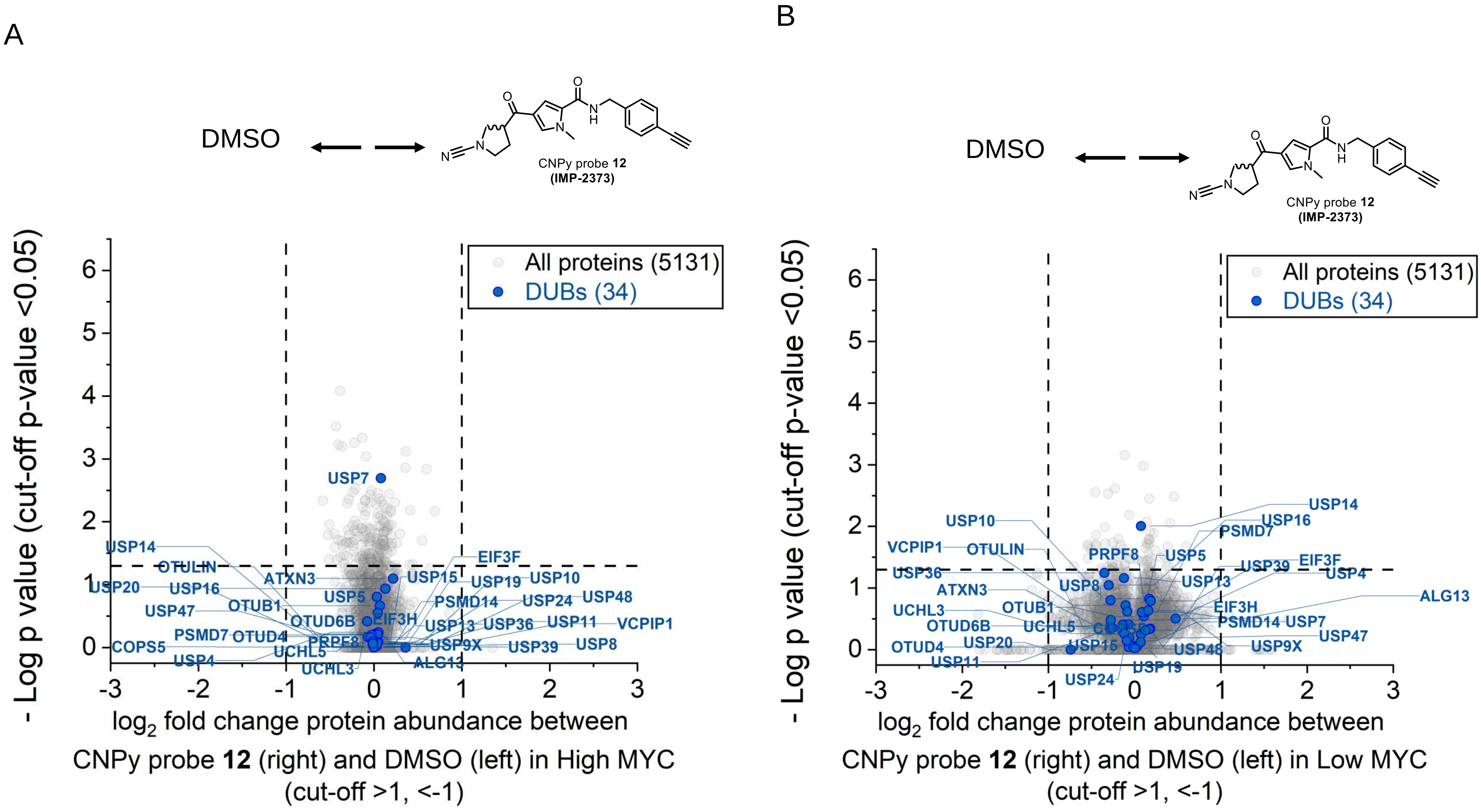


**Figure S11.** Changes DUB abundance in both high (**A**) and low (**B**) MYC cell systems after treatment with CNPy probe **12** (**IMP-2373**) for 1 h at 37°C. Quantitative detection of DUB activity in both high (**C**) and low (**D**) MYC cell systems. **E –** Changes in DUB abundance when switching from a high to low MYC system **F -** Changes in DUB activity when switching from a high to low MYC system.

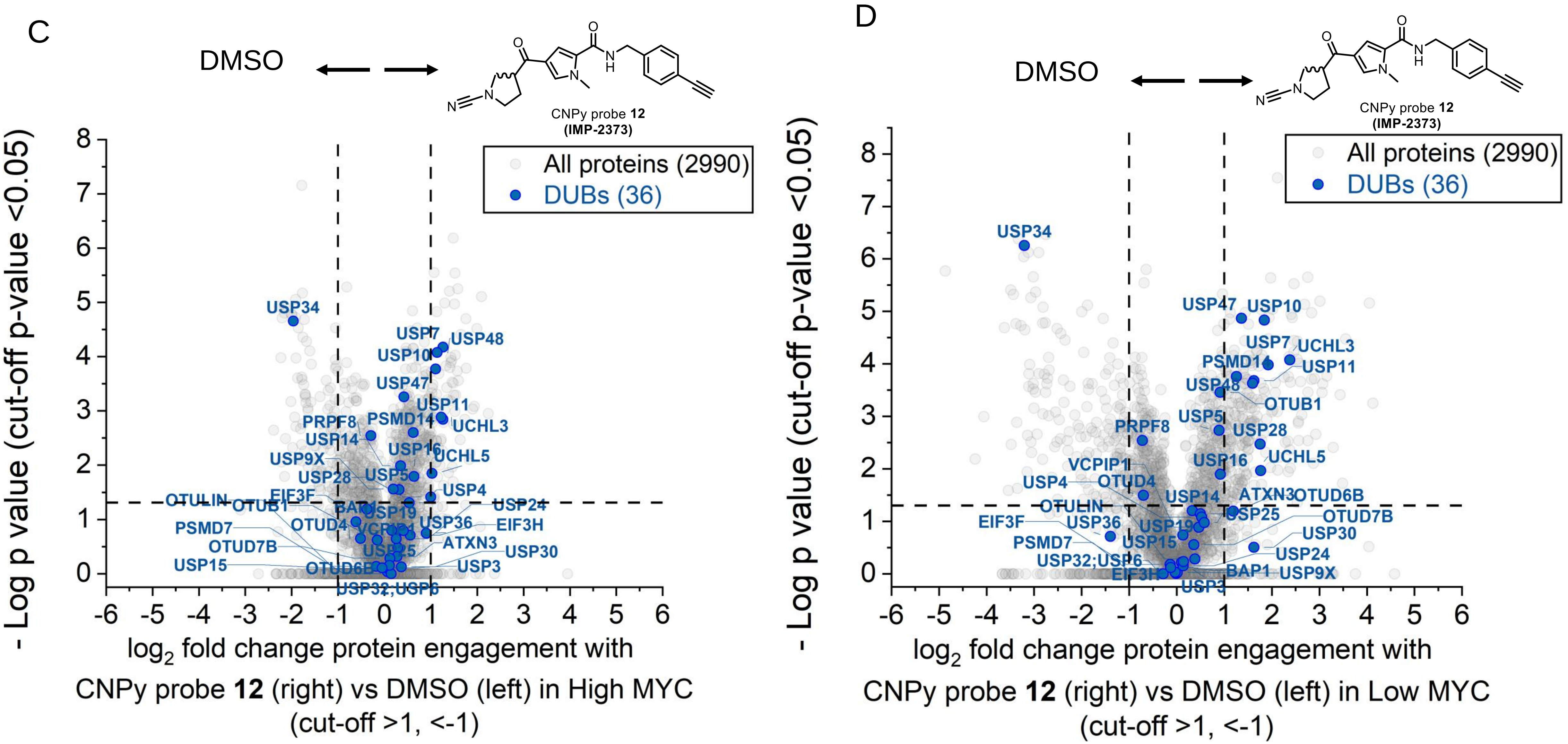


**Figure S11.** Changes DUB abundance in both high (**A**) and low (**B**) MYC cell systems after treatment with CNPy probe **12** (**IMP-2373**) for 1 h at 37°C. Quantitative detection of DUB activity (by target engagement with CNPy probe **12**) in both high (**C**) and low (**D**) MYC cell systems. **E –** Changes in DUB abundance when switching from a high to low MYC system **F -** Changes in DUB activity when switching from a high to low MYC system.

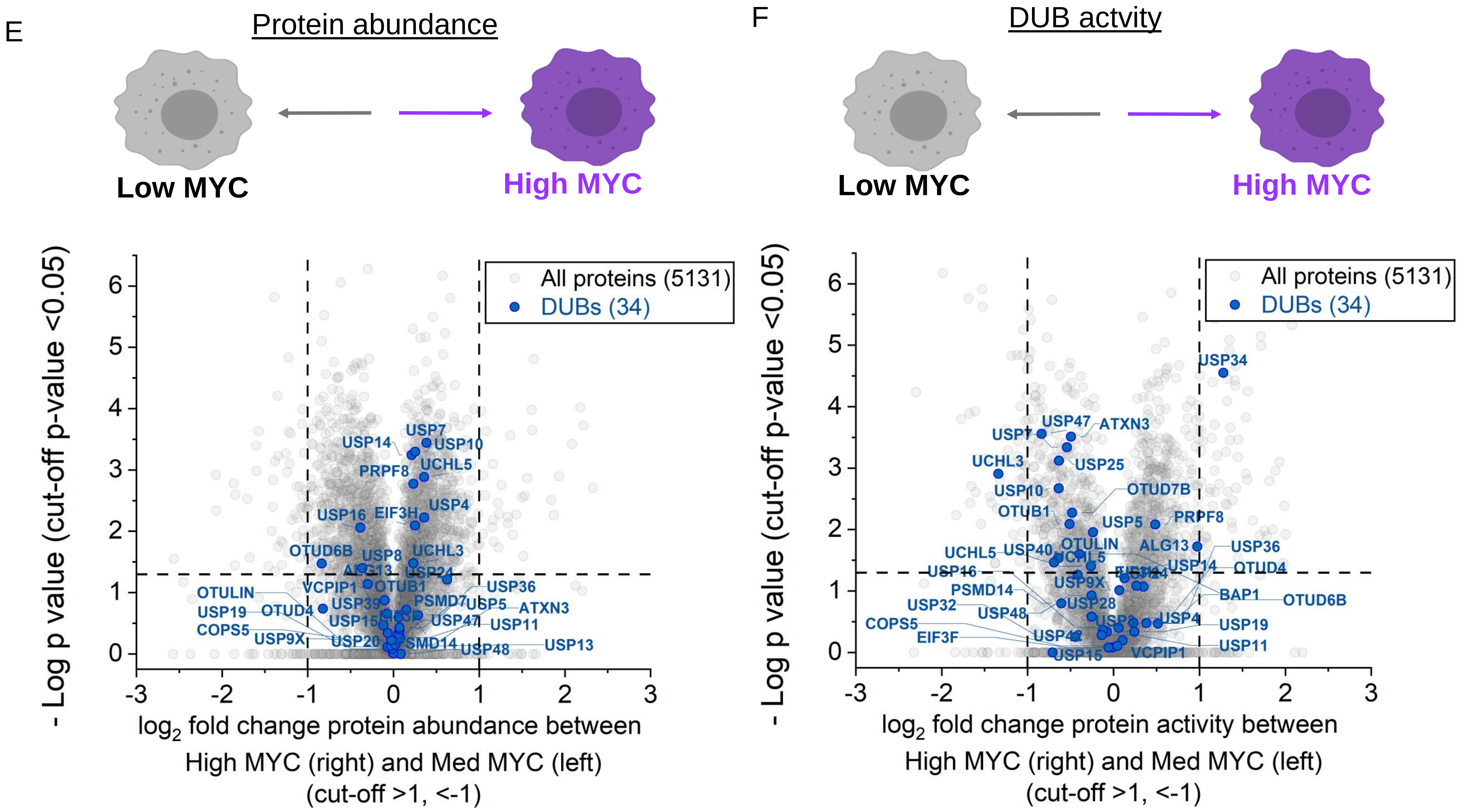


**Figure S11.** Changes DUB abundance in both high (**A**) and low (**B**) MYC cell systems after treatment with CNPy probe **12** (**IMP-2373**) for 1 h at 37°C. Quantitative detection of DUB activity in both high (**C**) and low (**D**) MYC cell systems. **E –** Changes in DUB abundance when switching from a high to low MYC system **F -** Changes in DUB activity when switching from a high to low MYC system.

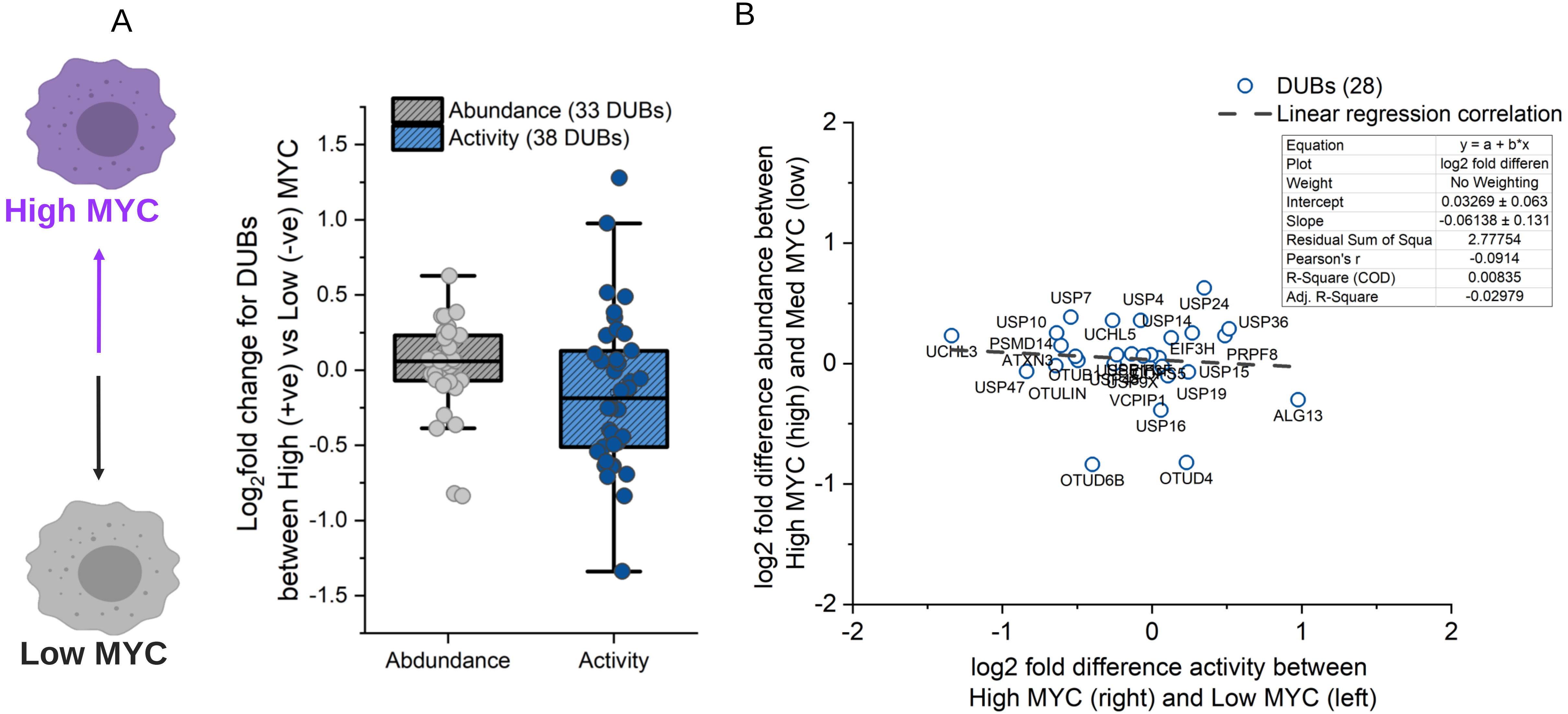


**Figure S12. A –** Overall changes in DUB abundance and activity between high and low MYC cell lines. **B -** Linear regression (Pearson correlation) model for changes in DUB abundance vs DUB activity in high vs low MYC cell models (data from Fig 4E). For 28 DUBs across the 2 datasets, no correlation was observed ($R^2$ value = 0.008).
