## Supplementary information 2 - Materials and methods for "Electrophile scanning by chemical proteomics reveals a potent pan-active DUB probe for investigation of deubiquitinase activity in live cells"

1. **Full chemical structures of probes investigated in this work**

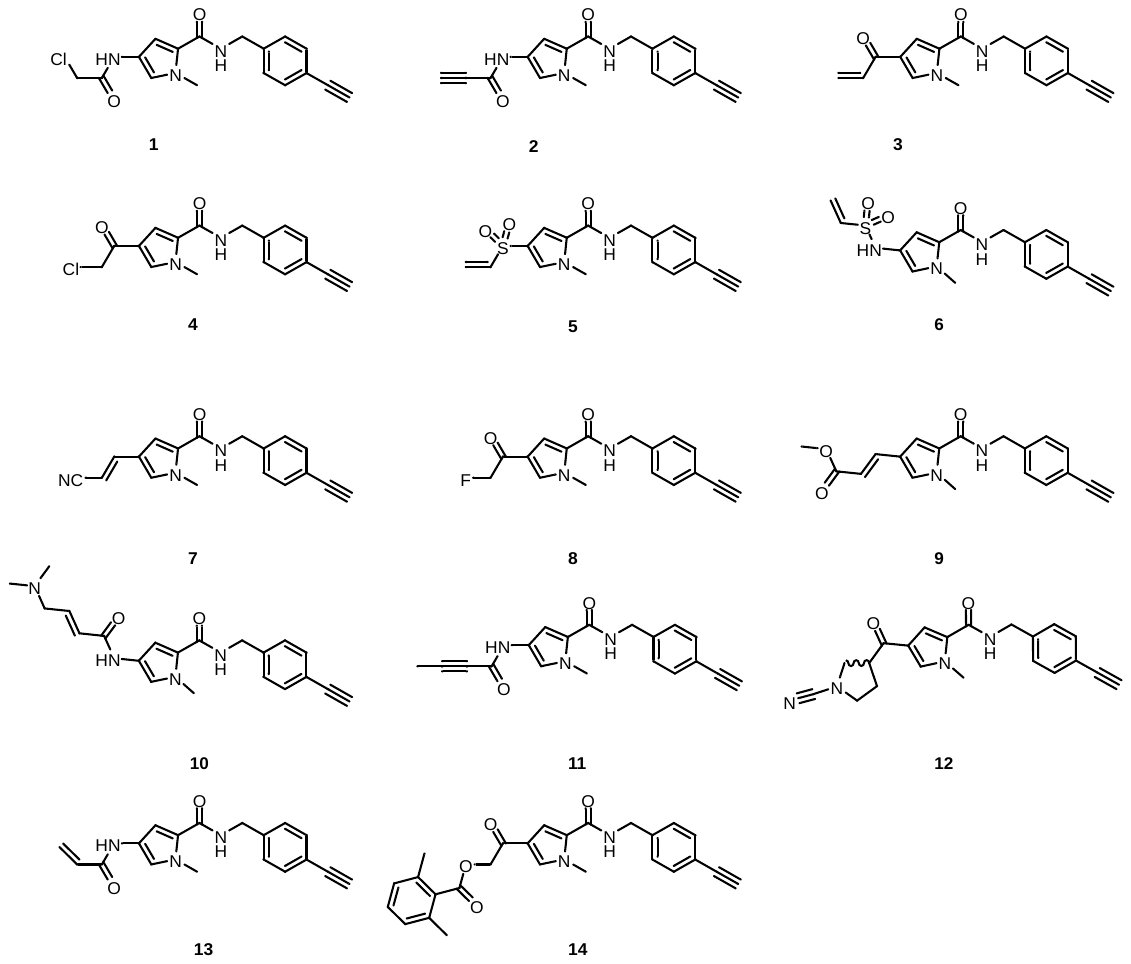

2. **Chemical synthesis of probes**

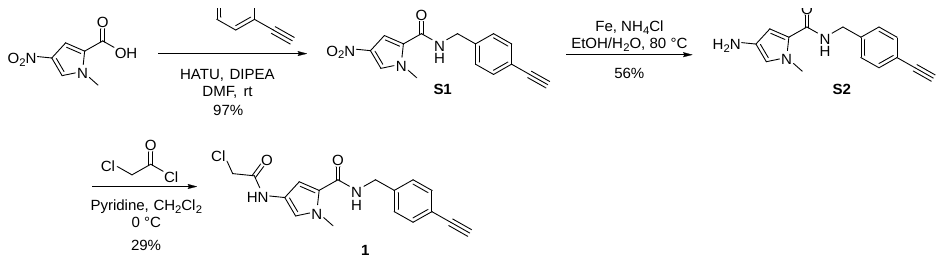

***N*-(4-ethynylbenzyl)-1-methyl-4-nitro-1*H*-pyrrole-2-carboxamide** **(S1):** To a mixture of 1-methyl-4-nitro-1*H*-pyrrole-2-carboxylic acid (310 mg, 1.82 mmol) and HATU (1.04 g, 2.73 mmol) in DMF (15 mL) was added DIPEA (589 mg, 4.56 mmol) and (4-ethynylphenyl)methanamine (239 mg, 1.82 mmol), then the mixture was stirred at 20 °C for 1.5 hours. LCMS showed starting material was consumed and a peak with the desired mass was formed. The reaction was quenched with water (40 mL). The mixture was extracted with EtOAc (50 mL x 2), and the combined organic layers were washed with water (20 mL x 2) and brine (20 mL x 2), dried over anhydrous Na_2_SO_4_, filtered, and concentrated under reduced pressure. The residue was purified silica gel chromatography (0-70% petroleum ether/EtOAc) to afford **S1** (500 mg, 96.9%) as an off-white solid. ^1^H NMR (400 MHz, DMSO-d_6_) δ 8.99 (t, *J* = 5.9 Hz, 1H), 8.15 (d, *J* = 1.8 Hz, 1H), 7.51 (d, *J* = 2.0 Hz, 1H), 7.45 (d, *J* = 8.2 Hz, 2H), 7.31 (d, *J* = 8.1 Hz, 2H), 4.42 (d, *J* = 6.0 Hz, 2H), 4.15 (s, 1H), 3.91 (s, 3H). MS: [M+H]^+^ 284.1.

**4-amino-*N*-(4-ethynylbenzyl)-1-methyl-1*H*-pyrrole-2-carboxamide** **(S2):** To a mixture of *N*-(4-ethynylbenzyl)-1-methyl-4-nitro-1*H*-pyrrole-2-carboxamide (500 mg, 1.77 mmol) in EtOH (15 mL) and H_2_O (15 mL) was added Fe (986 mg, 17.6 mmol) and NH_4_Cl (944 mg, 17.6 mmol), then the mixture was stirred at 80 °C for 2 hours. LCMS showed starting material was consumed and a peak with the desired mass was formed. The mixture was filtered through celite and washed with EtOAc (30 mL). Saturated aqueous Na_2_CO_3_ was added (~pH 9), and then the mixture was extracted with EtOAc (50 mL x 2). The combined organic layers were washed with water (30 mL) and brine (30 mL), dried over anhydrous Na_2_SO_4_, filtered, and concentrated under reduced pressure. The residue was purified by silica gel chromatography (0-7 % CH_2_Cl_2_/MeOH) to afford **S2** (250 mg, 55.9%) as brown oil. ^1^H NMR (400 MHz, DMSO-d_6_) δ 8.32 (t, *J* = 6.2 Hz, 1H), 7.42 (d, *J* = 8.2 Hz, 2H), 7.27 (d, *J* = 8.3 Hz, 2H), 6.26 (d, *J* = 2.1 Hz, 1H), 6.23 (d, *J* = 2.0 Hz, 1H), 4.35 (d, *J* = 6.1 Hz, 2H), 4.13 (s, 1H), 3.86 (bs, 2H), 3.68 (s, 3H). MS: [M+H]^+^ 254.1

**4-(2-chloroacetamido)-*N*-(4-ethynylbenzyl)-1-methyl-1*H*-pyrrole-2-carboxamide (1):** To a solution of 4-amino-*N*-(4-ethynylbenzyl)-1-methyl-1*H*-pyrrole-2-carboxamide (50 mg, 0.20 mmol) in CH_2_Cl_2_ (10 mL) was added 2-chloroacetyl chloride (33.4 mg, 0.296 mmol) and pyridine (31.2 mg, 0.395 mmol) at 0 °C and the mixture was stirred at 0 °C for 1 hour. LCMS indicated starting material was consumed and a peak with the desired mass was formed. The mixture was washed with water (20 mL x 2) and brine (20 mL), dried over anhydrous Na_2_SO_4_, filtered, and concentrated under reduced pressure. The crude product was purified by preparative TLC (1:2, petroleum ether/EtOAc) to afford **1** (19mg, 29% yield) as an off-white solid. ^1^H NMR (400 MHz, CDCl_3_) δ = 8.06 (br s, 1H), 7.47 (d, *J* = 7.8 Hz, 2H), 7.29 - 7.28 (m, 2H), 7.13 (s, 1H), 6.58 (s, 1H), 6.12 (br s, 1H), 4.56 (s, 2H), 4.16 (s, 2H), 3.94 (s, 3H), 3.07 (s, 1H). MS: [M+H]^+^ 330.1

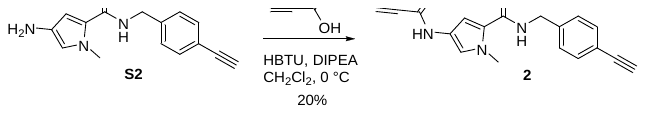

***N*-(4-ethynylbenzyl)-1-methyl-4-propiolamido-1*H*-pyrrole-2-carboxamide (2):** To a mixture of but-2-ynoic acid (24.9 mg, 0.296 mmol), 4-amino-*N*-(4-ethynylbenzyl)-1-methyl-1*H*-pyrrole-2-carboxamide (50 mg, 0.20 mmol) and HBTU (112 mg, 0.296 mmol) in CH_2_Cl_2_ (10 mL) was added DIPEA (63.8 mg, 0.493 mmol) at 0 °C and the mixture was stirred at 0 °C for 1 hour. LCMS indicated starting material was consumed and a peak with the desire mass was formed. The mixture was washed with water (20 mL x 2) and brine (20 mL), dried over anhydrous Na_2_SO_4_, filtered, and concentrated under reduced pressure. The crude product was purified by preparative TLC (1:1, petroleum ether/EtOAc) followed by HPLC (Column: Xbridge-C18 150 x 19 mm, 5 µm; Mobile Phase: 30-40% MeCN/H_2_O (0.1% formic acid); Flow rate: 20 mL/min; Wavelength: 214 nm) to afford **2** (12 mg, 20% yield) as an off-white solid. ^1^H NMR (400 MHz, DMSO-d_6_) δ = 10.78 (s, 1H), 8.66 (t, *J* = 5.7 Hz, 1H), 7.43 (d, *J* = 7.8 Hz, 2H), 7.28 (d, *J* = 7.8 Hz, 2H), 7.14 (s, 1H), 6.81 (d, *J* = 1.2 Hz, 1H), 4.37 (d, *J* = 5.9 Hz, 2H), 4.29 (s, 1H), 4.13 (s, 1H), 3.79 (s, 3H). MS: [M+H]^+^ 306.1

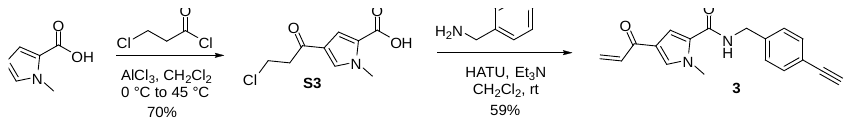

**4-(3-chloropropanoyl)-1-methyl-1*H*-pyrrole-2-carboxylic acid (S3):** To a stirred solution of 1-methyl-1*H*-pyrrole-2-carboxylic acid (1 g, 7.992 mmol) in CH_2_Cl_2_ (15 mL) was added AlCl_3_ (2.13 g, 16 mmol) at 0 °C. The reaction mixture was stirred at 0 °C for 30 min, then 3-chloropropionyl chloride (0.916 mL, 9.59 mmol) was added. The reaction mixture was stirred at 45 °C for 16 h. LCMS indicated starting material was consumed and a peak with the desired mass was formed. The reaction mixture was basified by the addition of aqueous NaHCO_3_ (~pH 9) and filtered through a celite pad. The filtrate was acidified by the addition of HCl (1 N) to ~pH 4 and the solid was collected by filtration to afford **S3** (1.2 g, yield 69.6%) as a white solid. ^1^H NMR (400 MHz, DMSO-d_6_) δ 12.69 (s, 1H), 7.90 (d, *J* = 1.8 Hz, 1H), 7.19 (d, *J* = 2.0 Hz, 1H), 3.90 – 3.86 (m, 2H), 3.89 (s, 3H), 3.25 (t, *J* = 6.3 Hz, 2H). MS: [M+H]^+^ 216.03

**4-acryloyl-*N*-(4-ethynylbenzyl)-1-methyl-1*H*-pyrrole-2-carboxamide (3):** To a mixture of 4-(3-chloropropanoyl)-1-methyl-1*H*-pyrrole-2-carboxylic acid (150 mg, 0.696 mmol), HATU (397 mg, 1.04 mmol), Et_3_N (0.482 mL,3.48 mmol) in CH_2_Cl_2_ (20 mL) at 15 °C was added a solution of (4-ethynylphenyl) methenamine (109 mg, 0.835 mmol) dropwise, and the reaction mixture was stirred at 15 °C for 14 hours. LCMS indicated starting material was consumed and a peak with the desired mass was formed. The mixture was washed with saturated NaHCO_3_, dried over anhydrous Na_2_SO_4_, filtered and concentrated under reduced pressure. The crude product was purified by silica gel chromatography (0-50%, petroleum ether/EtOAc) to afford **3** (110 mg, 59% yield) as a white solid. ^1^H NMR (400 MHz, DMSO-d_6_) δ = 8.88 (t, *J* = 6.1 Hz, 1H), 7.93 (d, *J* = 1.7 Hz, 1H), 7.44 (d, *J* = 7.7 Hz, 2H), 7.38 (d, *J* = 2.0 Hz, 1H), 7.31 (d, *J* = 8.6 Hz, 2H), 7.06 (dd, *J* = 10.3, 17.1 Hz, 1H), 6.26 (dd, *J* = 2.1, 17.0 Hz, 1H), 5.81 (dd, *J* = 2.0, 10.3 Hz, 1H), 4.41 (d, *J* = 6.1 Hz, 2H), 4.15 (s, 1H), 3.90 (s, 3H). MS: [M+H]^+^ 293.2

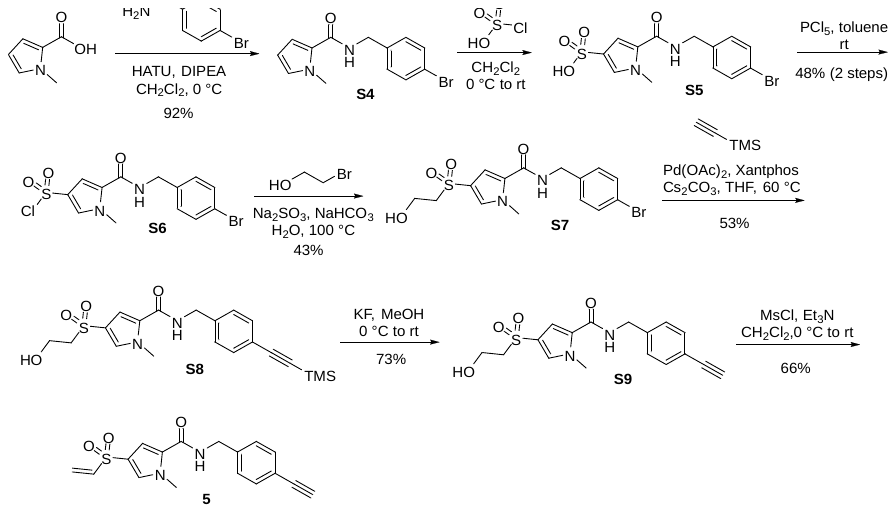

***N*-(4-bromobenzyl)-1-methyl-1*H*-pyrrole-2-carboxamide (S4):** To a stirred solution of 1-methyl-1*H*-pyrrole-2-carboxylic acid (1.25 g, 9.99 mmol), HATU (4.56 g, 12 mmol), DIPEA (1.93 g, 15 mmol) in CH_2_Cl_2_ (50 mL) at 0 °C was added (4-bromophenyl)methanamine (2.04 g, 11 mmol). The reaction mixture was stirred at 0 °C for 1 hour. LCMS showed starting material was consumed and a peak with the desired mass was formed. The reaction was quenched with water (20 mL), and the organic phase was separated and washed with a saturated NaHCO_3_ solution (50 mL) and brine (20 mL), dried over anhydrous Na_2_SO_4_, filtered, and then concentrated under reduced pressure. The residue was purified silica gel chromatography (0-20 % Petroleum ether/EtOAc) to afford **S4** (2.7 g, 92% yield) as a white solid. ^1^H NMR (400 MHz, CD_3_OD) δ 7.46 (d, *J* = 8.4 Hz, 2H), 7.24 (d, *J* = 8.3 Hz, 2H), 6.81 (m, 1H), 6.73 (dd, *J* = 3.9, 2.6 Hz, 1H), 6.05 (dd, *J* = 3.9, 2.6 Hz, 1H), 4.44 (s, 2H), 3.88 (s, 3H). MS: [M+H]^+^ 293.02

**5-((4-bromobenzyl) carbamoyl)-1-methyl-1*H*-pyrrole-3-sulfonic acid (S5):** To a stirred solution of *N*-(4-bromobenzyl)-1-methyl-1*H*-pyrrole-2-carboxamide (2.7 g, 9.2 mmol) in CH_2_Cl_2_ (50 mL) was added chlorosulfonic acid (1.4 g, 12 mmol) at 0 °C. The reaction mixture was stirred at 0 °C for 30 minutes and then at 20 °C for 3 hours. LCMS showed starting material was consumed and a peak with the desired mass was formed. The reaction mixture was concentrated to afford **S5** (4.0 g, crude) as a yellow solid, which was used directly without additional purification. MS: [M+H]^+^ 372.9

**5-((4-bromobenzyl) carbamoyl)-1-methyl-1*H*-pyrrole-3-sulfonyl chloride (S6):** To a stirred solution of 5-((4-bromobenzyl)carbamoyl)-1-methyl-1*H*-pyrrole-3-sulfonic acid (4.0 g, 10.7 mmol) in toluene (100 mL) was added PCl_5_ (4.46 g, 21.4 mmol) at 0 °C. The reaction mixture was stirred at 15 °C for 14 hours. LCMS showed starting material was consumed and a peak with the desired mass was formed. The reaction mixture was poured into ice water (100 mL) and then the aqueous layer was extracted with EtOAc (50 mL x 2), the separated organic layer was dried over anhydrous Na_2_SO_4_ and concentrated under reduced pressure. The residue was purified by silica gel chromatography (0-50% petroleum ether/EtOAc) to afford **S6** (2.0 g, 47.6% yield) as a white solid. ^1^H NMR (400 MHz, DMSO-d_6_) δ 8.68 (t, *J* = 5.6 Hz, 1H), 7.50 (d, *J* = 8.3 Hz, 2H), 7.24 (d, *J* = 8.3 Hz, 2H), 7.02 (d, *J* = 1.6 Hz, 1H), 6.95 (d, *J* = 1.7 Hz, 1H), 4.33 (d, *J* = 4.5 Hz, 2H), 3.79 (s, 3H). MS: [M+H]^+^ 390.9

***N*-(4-bromobenzyl)-4-((2-hydroxyethyl)sulfonyl)-1-methyl-1*H*-pyrrole-2-carboxamide** **(S7):** To a mixture of 5-((4-bromobenzyl)carbamoyl)-1-methyl-1*H*-pyrrole-3-sulfonyl chloride (450 mg, 1.15 mmol), Na_2_SO_3_ (159 mg, 1.26 mmol) and NaHCO_3_ (290 mg, 3.45 mmol) in H_2_O (5 mL), then the reaction mixture was heated to 100 °C for 1 hour. After that the reaction was cooled to room temperature, 2-bromoethan-1-ol (215 mg, 1.72 mmol) was added, then the mixture was heated to 100 °C for 4 hours. LCMS showed starting material was consumed and a peak with the desired mass was formed. The mixture reaction was cooled to room temperature and poured into water (20 mL), the mixture was extracted with EtOAc (10 mL x 2), the combined organic layers were washed by brine (20 mL) and dried over anhydrous Na_2_SO_4_ and concentrated under reduced pressure. The crude product was purified silica gel chromatography (0-20% CH_2_Cl_2_/MeOH) to afford **S7** (200 mg, yield 43.4%) as a white solid. ^1^H NMR (400 MHz, CD_3_OD) δ 7.49 (d, *J* = 1.8 Hz, 1H), 7.47 (d, *J* = 8.5 Hz, 2H), 7.25 (d, *J* = 8.5 Hz, 2H), 7.11 (d, *J* = 1.9 Hz, 1H), 4.45 (s, 2H), 3.95 (s, 3H), 3.87 (t, *J* = 6.4 Hz, 2H), 3.35 (t, *J* = 6.4 Hz, 2H). MS: [M+H]^+^ 401.1

**4-((2-hydroxyethyl)sulfonyl)-1-methyl-N-(4-((trimethylsilyl) ethynyl) benzyl)-1*H*-pyrrole-2-carboxamide (S8):** To a mixture of *N*-(4-bromobenzyl)-4-((2-hydroxyethyl)sulfonyl)-1-methyl-1*H*-pyrrole-2-carboxamide (900 mg, 2.24 mmol), ethynyltrimethylsilane (1 mL, 7 mmol) and Cs_2_CO_3_ (1.46 g, 4.5 mmol) in THF (5 mL) was added Pd(OAc)_2_ (50 mg, 0.22 mmol) and Xantphos (259 mg, 0.449 mmol) under N_2_ protection. The reaction was stirred at 60 °C for 8 hours under N_2_. LCMS showed starting material was consumed a peak with the desired mass was formed. The reaction mixture was cooled to room temperature and quenched with water (20 mL). The mixture was extracted with EtOAc (10 mL x 2). The combined organic layers were washed by brine and dried over anhydrous Na_2_SO_4_, filtered and concentrated under reduced pressure. The residue was purified silica gel chromatography (0-10% CH_2_Cl_2_/MeOH) to afford **S8** (500 mg, yield 53.3%) as a yellow gum. ^1^H NMR (400 MHz, CD_3_OD) δ 7.50 (d, *J* = 1.8 Hz, 1H), 7.39 (d, *J* = 8.3 Hz, 2H), 7.30 (d, *J* = 8.3 Hz, 2H), 7.12 (d, *J* = 1.9 Hz, 1H), 4.49 (s, 2H), 3.96 (s, 3H), 3.88 (t, *J* = 6.4 Hz, 2H), 3.36 (t, *J* = 6.4 Hz, 2H), 0.22 (m, 9H). MS: [M+H]^+^ 419.1

***N*-(4-ethynylbenzyl)-4-((2-hydroxyethyl)sulfonyl)-1-methyl-1*H*-pyrrole-2-carboxamide (S9):** To a solution of 4-(ethylsulfonyl)-1-methyl-*N*-(4-((trimethylsilyl)ethynyl)benzyl)-1*H*-pyrrole-2-carboxamide (500 mg, 1.19 mmol) in MeOH (15 mL) was added KF (346 mg, 5.97 mmol) at 0 °C. The reaction was stirred at 20 °C for 3 hours. LCMS showed starting material was consumed and a peak with the desired mass was formed. The reaction mixture was quenched with water (20 mL) and was extracted with ethyl acetate (10 mL x 2). The combined organic layers were washed with brine (10 mL), dried over anhydrous Na_2_SO_4_, filtered, and concentrated under reduced pressure. The residue was purified silica gel chromatography (0-10% CH­_2_Cl_2_/MeOH) to afford **S9** (300 mg, yield: 72.5%) as a yellow gum. ^1^H NMR (400 MHz, CD_3_OD) δ 7.50 (d, *J* = 1.7 Hz, 1H), 7.43 (d, *J* = 8.4 Hz, 2H), 7.31 (d, *J* = 8.4 Hz, 2H), 7.11 (d, *J* = 1.9 Hz, 1H), 4.50 (s, 2H), 3.95 (s, 3H), 3.87 (t, *J* = 6.4 Hz, 2H), 3.45 (s, 1H), 3.35 (t, *J* = 6.4 Hz, 2H). MS: [M+H]^+^ 347.1

***N*-(4-ethynylbenzyl)-1-methyl-4-(vinylsulfonyl)-1*H*-pyrrole-2-carboxamide (5):** To a mixture of *N*-(4-ethynylbenzyl)-4-((2-hydroxyethyl)sulfonyl)-1-methyl-1*H*-pyrrole-2-carboxamide (80 mg, 0.23 mmol) in CH_2_Cl_2_ (5 mL) at 0 ^o^C was added Et_3_N (0.1 mL, 0.7 mmol), followed by MsCl (40 mg,0.35 mmol) in CH_2_Cl_2_ (1 mL) and the reaction was stirred at 20 °C for 2 h. LCMS indicated the desired product mass was present and the reaction was poured over water (20 mL). The mixture was extracted with EtOAc (2x10 mL) and the combined organic fractions were dried over anhydrous Na_2_SO_4_, filtered and concentrated under reduced pressure. The crude product was purified by silica gel chromatography (0-50% petroleum ether/EtOAc) to afford **5** (50 mg, 66% yield) as a white solid. ^1^H NMR (400 MHz, DMSO-d_6_) δ = 8.91 (t, *J* = 5.9 Hz, 1H), 7.66 (d, *J* = 1.3 Hz, 1H), 7.44 (d, *J* = 7.9 Hz, 2H), 7.30 (d, *J* = 7.9 Hz, 2H), 7.19 (d, *J* = 1.8 Hz, 1H), 7.00 (dd, *J* = 9.7, 16.2 Hz, 1H), 6.15 (d, *J* = 16.7 Hz, 1H), 6.04 (d, *J* = 10.1 Hz, 1H), 4.39 (d, *J* = 5.7 Hz, 2H), 4.15 (s, 1H), 3.89 (s, 3H). MS: [M+H]^+^ 329.1

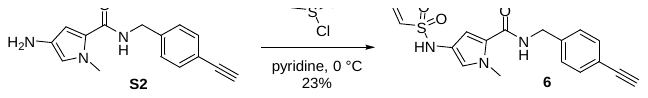

***N*-(4-ethynylbenzyl)-1-methyl-4-(vinylsulfonamido)-1*H*-pyrrole-2-carboxamide (6):** To a solution of 4-amino-*N*-(4-ethynylbenzyl)-1-methyl-1*H*-pyrrole-2-carboxamide (70 mg, 0.28 mmol) was added ethenesulfonyl chloride (35 mg, 0.276mmol) in pyridine (26.2 mg, 0.332mmol) at 0 °C and the mixture was stirred at 0 °C for 1 hour. LCMS indicated starting material was consumed and a peak with the desired mass was formed. The mixture was washed with water (20 mL x 2) and brine (20 mL), dried over anhydrous Na_2_SO_4_, filtered, and concentrated under reduced pressure. The crude product was purified by HPLC (Column: Kromasil-C18 100 x 21.2 mm, 5 µm; Mobile Phase: 35-45% MeCN/H_2_O (0.1% formic acid); Flowrate: 20 mL/min; Wavelength: 214 nm) to afford **6** (22 mg, 23% yield) as an off-white solid. ^1^H NMR (400 MHz, DMSO-d_6_) δ = 9.33 (s, 1H), 8.61 (t, *J* = 6.0 Hz, 1H), 7.43 (d, *J* = 8.1 Hz, 2H), 7.28 (d, *J* = 8.1 Hz, 2H), 6.75 - 6.65 (m, 3H), 6.04 - 5.95 (m, 2H), 4.36 (d, *J* = 6.1 Hz, 2H), 4.13 (s, 1H), 3.77 (s, 3H). MS: [M+H]^+^ 344.1

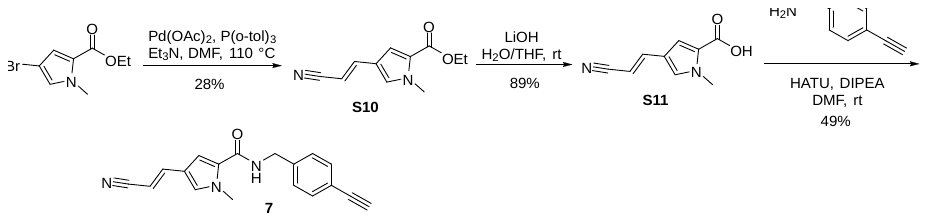

**Ethyl (*E*)-4-(2-cyanovinyl)-1-methyl-1*H*-pyrrole-2-carboxylate** **(S10):** A mixture of ethyl 4-bromo-1-methyl-1*H*-pyrrole-2-carboxylate (300 mg, 1.29 mmol), acrylonitrile (549 mg, 10.3 mmol), Pd(OAc)_2_ (58 mg, 0.259 mmol), (*o*-MeC_6_H_4_)_3_P (177 mg, 0.582 mmol) and Et_3_N (392 mg, 3.88 mmol) in DMF (10 mL) was stirred at 110 °C for 24 hours in a 25 mL sealed tube under N_2_. LCMS showed trace starting material remained and the desired mass peak was formed. The reaction was quenched with water (20 mL). The mixture was extracted with EtOAc (50 mL x 2). The combined organic layer was washed with water (20 mL x 2) and brine (20 mL x 2), dried over anhydrous Na_2_SO_4_, filtered, and concentrated under reduced pressure. The residue was purified by silica gel chromatography (0-30% petroleum ether/EtOAc) to afford **S10** (75 mg, 28%) as a yellow solid. ^1^H NMR (400 MHz, CDCl_3_) δ 7.20 (d, *J* = 16.4 Hz, 1H), 7.08 (d, *J* = 1.9 Hz, 1H), 6.96 (d, *J* = 1.7 Hz, 1H), 5.52 (d, *J* = 16.4 Hz, 1H), 4.30 (d, *J* = 7.1 Hz, 2H), 3.92 (s, 3H), 1.36 (t, *J* = 7.1 Hz, 3H). MS: [M+H]^+^ 205.1

***(E)*-4-(2-cyanovinyl)-1-methyl-1*H*-pyrrole-2-carboxylic acid** **(S11):** To a mixture of ethyl (*E*)-4-(2-cyanovinyl)-1-methyl-1*H*-pyrrole-2-carboxylate (65 mg, 0.32 mmol) in THF (3 mL) and H_2_O (3 mL) was added LiOH (22.9 mg, 0.955 mmol), then the mixture was stirred at 20 °C for 16 hours. LCMS showed starting material was consumed and a peak with the desired mass was formed. The mixture was acidified by the addition of 2 N HCl to ~pH 3 and then the mixture was extracted with EtOAc (50 mL x 2). The combined organic fraction was washed with water (30 mL) and brine (30 mL), dried over anhydrous Na_2_SO_4_, filtered, and concentrated under reduced pressure to afford **S11** (50 mg, 89%) as a yellow solid. ^1^H NMR (400 MHz, DMSO-d_6_) δ 12.59 (s, 1H), 7.42 (d, *J* = 1.9 Hz, 1H), 7.40 (d, *J* = 16.4 Hz, 1H), 7.19 (d, *J* = 1.9 Hz, 1H), 5.99 (d, *J* = 16.4 Hz, 1H), 3.84 (s, 3H). MS: (M-H)^-^ 175.1

***(E)*-4-(2-cyanovinyl)-*N*-(4-ethynylbenzyl)-1-methyl-1*H*-pyrrole-2-carboxamide (7):** To a mixture of (*E*)-4-(2-cyanovinyl)-1-methyl-1*H*-pyrrole-2-carboxylic acid (45 mg, 0.26 mmol), (4-ethynylphenyl) methanamine (33.5 mg, 0.255 mmol) and HATU (146 mg, 0.383 mmol) in DMF (5 mL) was added DIPEA (82.5 mg, 0.639 mmol) and the mixture was stirred at 20 °C for 2 hours. LCMS indicated starting material was consumed and a peak with the desire mass was formed. The reaction was quenched with water (10 mL). The mixture was extracted with EtOAc (30 mL x 2). The combined organic fraction was washed with water (20 mL x 2) and brine (20 mL), dried over anhydrous Na_2_SO_4_, filtered, and concentrated under reduced pressure. The crude product was purified by preparative TLC (1:1 petroleum ether/EtOAc) to afford **7** (36 mg, 49% yield) as an off-white solid. ^1^H NMR (400 MHz, DMSO-d_6_) δ = 8.78 (t, *J* = 6.0 Hz, 1H), 7.44 (d, *J* = 8.1 Hz, 2H), 7.41 (s, 1H), 7.36 (s, 1H), 7.30 (d, *J* = 8.1 Hz, 2H), 7.11 (s, 1H), 5.76 (d, *J* = 16.4 Hz, 1H), 4.40 (d, *J* = 5.9 Hz, 2H), 4.14 (s, 1H), 3.83 (s, 3H). MS: [M+H]^+^ 290.1

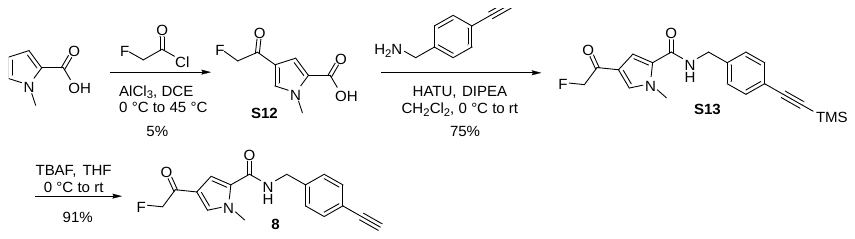

**4-(2-fluoroacetyl)-1-methyl-1*H*-pyrrole-2-carboxylic acid (S12):** To a stirred solution of 1-methyl-1*H*-pyrrole-2-carboxylic acid (1.25 g, 9.99 mmol) in DCE (50 mL) was added AlCl_3_ (3.33 g, 25 mmol) at 0 °C. The reaction mixture was stirred at 0 °C for 30 minutes, then 2-fluoroacetyl chloride (1.93 g, 20.0 mmol) was added. The reaction mixture was stirred at 45 °C for 15 hours. LCMS indicated trace starting material remained and a peak with the desired mass was formed. The reaction mixture was basified by the addition of LiOH.H_2_O (1M) and filtered through a celite pad. The filtrate was acidified by HCl (1 M) to ~pH 4 and the solid was collected by filtration. The residue was purified by reverse phase chromatography (Combi-Flash, 80 g C18, using a 0-30% gradient of MeCN in H_2_O (0.5% formic acid)) to afford the **S12** (100 mg, 5.4%) as a white solid. ^1^H NMR (400 MHz, DMSO-d_6_) δ 12.63 (s, 1H), 7.86 (d, *J* = 1.9 Hz, 1H), 7.20 (d, *J* = 2.0 Hz, 1H), 5.43 (d, *J* = 46.8 Hz, 2H), 3.89 (s, 3H).^19^F NMR (400 MHz, DMSO-d_6_) δ -230.76. MS: [M+H]^+^ 186.05

**4-(2-fluoroacetyl)-1-methyl-*N*-(4-((trimethylsilyl)ethynyl)benzyl)-1*H*-pyrrole-2-carboxamide (S13):** To a mixture of 4-(2-fluoroacetyl)-1-methyl-1*H*-pyrrole-2-carboxylic acid (100 mg, 0.54 mmol) in CH_2_Cl­_2_ (10 mL) was added HATU (267 mg, 0.702 mmol), DIPEA (139 mg, 1.08 mmol) and (4-((trimethylsilyl)ethynyl) phenyl) methenamine (132 mg, 0.648 mmol) at 0 °C, then the mixture was stirred at 25 °C for 3 hours. LCMS indicated starting material was consumed and a peak with the desired mass was formed. Then the reaction mixture was poured into water (30 mL). The organic layer was separated and the aqueous layer was extracted with CH_2_Cl_2_ (10 mL*2). The combined organic fraction was dried over anhydrous Na_2_SO_4_, filtered and concentrated under reduced pressure. The residue was purified by silica gel chromatography (0-50% petroleum ether/EtOAc) to afford **S13** (150 mg, 75%) as a white solid. MS: [M+H]^+^ 371.1

***N-*(4-ethynylbenzyl)-4-(2-fluoroacetyl)-1-methyl-1*H*-pyrrole-2-carboxamide (8):** To a mixture of 4-(2-fluoroacetyl)-1-methyl-*N*-(4-((trimethylsilyl)ethynyl) benzyl)-1*H*-pyrrole-2-carboxamide (150 mg, 0.405 mmol) in THF (10 mL) was added TBAF (0.4 mL, 0.4 mmol) at 0 °C. the mixture was allowed to warm to 25°C and stirred at this temperature for 30 mins. LCMS indicated starting material was consumed and a peak with the desired mass was formed. The mixture was concentrated and purified by silica gel chromatography (0-50% petroleum ether/EtOAc) to afford **8** (110 mg, 91% yield) as a white solid. ^1^H NMR (400 MHz, DMSO-d_6_) δ = 7.51 (t, *J* = 6.0 Hz, 1H), 7.47 (d, *J* = 8.4 Hz, 2H), 7.28 (d, *J* = 8.4 Hz, 2H), 7.14 (d, *J* = 1.6 Hz, 2H), 6.48 (s, 1H), 5.10 (d, *J* = 46.7 Hz, 2H), 4.56 (d, *J* = 5.9 Hz, 2H), 4.00 (s, 1H), 3.08 (s, 3H).^19^F NMR (400 MHz, DMSO-d_6_) δ -225.77. MS: [M+H]^+^ 299.1

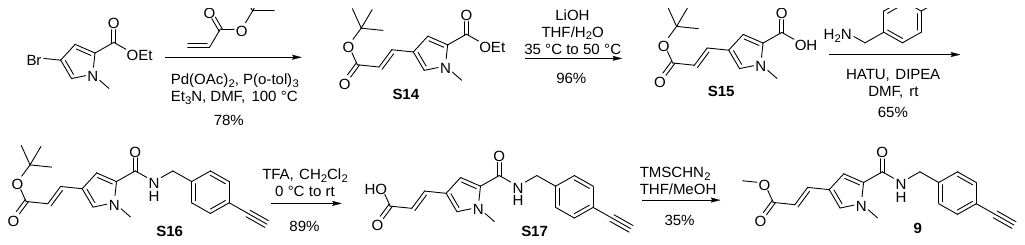

**Ethyl (E)-4-(3-(tert-butoxy)-3-oxoprop-1-en-1-yl)-1-methyl-*1H*-pyrrole-2-carboxylate** **(S14):** A mixture of ethyl 4-bromo-1-methyl-1*H*-pyrrole-2-carboxylate (300 mg, 1.29 mmol), *tert*-butyl acrylate (828 mg, 6.46mmol), Pd(OAc)_2_ (58 mg, 0.259 mmol), (*o*-MeC_6_H_4_)_3_P (177 mg, 0.582 mmol) and Et_3_N (392 mg, 3.88 mmol) in DMF (10 mL) was stirred at 100 °C for 60 hours in a 25 mL sealed tube under N_2_. LCMS indicated starting material was consumed and a peak with the desired mass was formed. The reaction was quenched with water (20 mL). The mixture was extracted with EtOAc (50 mL x 2). The combined organic fraction was washed with water (20 mL x 2) and brine (20 mL x 2), dried over anhydrous Na_2_SO_4_, filtered, and concentrated under reduced pressure. The residue was purified by silica gel chromatography (0-20 % petroleum ether/EtOAc) to afford **S14** (280 mg, 77.5%) as a yellow solid. ^1^H NMR (400 MHz, DMSO-d_6_) δ 7.54 (d, *J* = 1.7 Hz, 1H), 7.37 (d, *J* = 15.9 Hz, 1H), 7.19 (d, *J* = 1.9 Hz, 1H), 6.15 (d, *J* = 15.8 Hz, 1H), 4.22 (q, J = 7.1 Hz, 2H), 3.84 (s, 3H), 1.45 (s, 9H), 1.28 (t, *J* = 7.1 Hz, 3H). MS: [M+H]^+^ 280.3

***(E)*-4-(3-(tert-butoxy)-3-oxoprop-1-en-1-yl)-1-methyl-1*H*-pyrrole-2-carboxylic acid** **(S15):** To a mixture of ethyl (*E*)-4-(3-(tert-butoxy)-3-oxoprop-1-en-1-yl)-1-methyl-1*H*-pyrrole-2-carboxylate (220 mg, 0.788 mmol) in THF (6 mL) and H_2_O (6 mL) was added LiOH (94.3 mg, 3.94 mmol), then the mixture was stirred at 35 °C for 16 hours and then the mixture was stirred at 50 °C for 3 hours. LCMS indicated starting material was consumed and a peak with the desired mass was formed. The mixture was acidified by the addition of 2 N HCl to ~pH 3, and then the mixture was extracted with EtOAc (50 mL x 2). The organic fraction was washed with water (20 mL) and brine (30 mL), dried over anhydrous Na_2_SO_4_, filtered, and concentrated under reduced pressure to afford the **S15** (190 mg, 96%) as an off-white solid. ^1^H NMR (400 MHz, DMSO-d_6_) δ 12.42 (s, 1H), 7.49 (d, *J* = 1.5 Hz, 1H), 7.36 (d, *J* = 15.9 Hz, 1H), 7.13 (d, *J* = 1.6 Hz, 1H), 6.11 (d, *J* = 15.8 Hz, 1H), 3.83 (s, 3H), 1.45 (s, 9H). MS: [M-H]^-^ 250.1

***tert*-butyl (*E*)-3-(5-((4-ethynylbenzyl)carbamoyl)-1-methyl-1*H*-pyrrol-3-yl)acrylate** **(S16):** To a mixture of (*E*)-4-(3-(tert-butoxy)-3-oxoprop-1-en-1-yl)-1-methyl-1*H*-pyrrole-2-carboxylic acid (190 mg, 0.756 mmol), (4-ethynylphenyl) methanamine (109 mg, 0.832 mmol) and HATU (431 mg, 1.13mmol) in DMF (6 mL) was added DIPEA (244 mg, 1.89 mmol), then the mixture was stirred at 20 °C for 16 hours. LCMS indicated starting material was consumed and a peak with the desired mass was formed. The reaction was quenched with water (20 mL). The mixture was extracted with EtOAc (30 mL x 2). The combined organic fraction was washed with water (20 mL x 2) and brine (20 mL x 2), dried over anhydrous Na_2_SO_4_, filtered, and concentrated under reduced pressure. The residue was purified by silica gel chromatography (0-80 % petroleum ether/EtOAc) to afford **S16** (180 mg, 65.3%) as a yellow solid. ^1^H NMR (400 MHz, DMSO-d_6_) δ 8.65 (t, *J* = 6.0 Hz, 1H), 7.44 (d, *J* = 8.1 Hz, 2H), 7.37 (d, *J* = 15.6 Hz, 2H), 7.31 (d, *J* = 8.1 Hz, 2H), 7.17 (d, *J* = 1.5 Hz, 1H), 5.98 (d, *J* = 15.8 Hz, 1H), 4.41 (d, *J* = 5.9 Hz, 2H), 4.14 (s, 1H), 3.82 (s, 3H), 1.45 (s, 9H). MS: [M+H]^+^ 365.1

***(E)*-3-(5-((4-ethynylbenzyl) carbamoyl)-1-methyl-1*H*-pyrrol-3-yl) acrylic acid** **(S17):** To a solution of *tert*-butyl (*E*)-3-(5-((4-ethynylbenzyl) carbamoyl)-1-methyl-1*H*-pyrrol-3-yl) acrylate (160 mg, 0.439 mmol) in CH_2_Cl_2_ (15 mL) was added TFA (1.5 mL) at 0 °C, then the mixture was stirred at 20 °C for 2 hours. LCMS showed trace starting material remained and the desired mass peak was formed. The reaction was quenched with water (20 mL), basified by the addition of saturated NaHCO_3_ to ~pH 3 and then extracted with CH_2_Cl_2_ (30 mL x 3). The organic phase was washed with brine (30 mL), dried over anhydrous Na_2_SO_4_, filtered, and concentrated under reduced pressure to afford the **S17** (120 mg, 88.6%) as a yellow solid. MS: [M+H]^+^ 309.1

**Methyl (*E*)-3-(5-((4-ethynylbenzyl)carbamoyl)-1-methyl-1*H*-pyrrol-3-yl)acrylate (9):** To a mixture of (*E*)-3-(5-((4-ethynylbenzyl)carbamoyl)-1-methyl-1*H*-pyrrol-3-yl)acrylic acid (110 mg, 0.357 mmol) in THF (10 mL) and MeOH (3 mL) was added TMSCHN_2_ (204 mg, 1.78 mmol) and the mixture was stirred at 20 °C for 2 hours. LCMS indicated starting material was consumed and a peak with the desired mass was formed. The reaction was quenched with water (20 mL) and extracted with EtOAc (30 mL x 2). The combined organic fraction was washed with water (20 mL) and brine (20 mL), dried over anhydrous Na_2_SO_4_, filtered, and concentrated under reduced pressure. The crude product was purified by preparative TLC (2:1, petroleum ether/EtOAc) to afford **9** (40 mg, 35% yield) as an off-white solid. ^1^H NMR (400 MHz, DMSO-d_6_) δ = 8.71 (t, *J* = 6.0 Hz, 1H), 7.50 (d, *J* = 15.9 Hz, 1H), 7.44 (d, *J* = 8.3 Hz, 2H), 7.42 (d, *J* = 1.5 Hz, 1H), 7.31 (d, *J* = 8.3 Hz, 2H), 7.17 (d, *J* = 1.7 Hz, 1H), 6.09 (d, *J* = 15.7 Hz, 1H), 4.41 (d, *J* = 5.9 Hz, 2H), 4.14 (s, 1H), 3.83 (s, 3H), 3.67 (s, 3H). MS: [M+H]^+^ 323.1

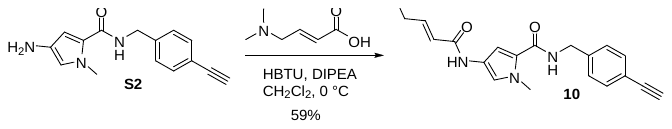

***(E)*-4-(4-(dimethylamino)but-2-enamido)-*N*-(4-ethynylbenzyl)-1-methyl-1*H*-pyrrole-2-carboxamide (10):** To a mixture of (*E*)-4-(dimethylamino)but-2-enoic acid (69 mg, 0.53 mmol), 4-amino-*N*-(4-ethynylbenzyl)-1-methyl-1*H*-pyrrole-2-carboxamide (70 mg, 0.28 mmol) and HBTU (199 mg, 0.53 mmol) in CH_2_Cl_2_ (10 mL) was added DIPEA (90 mg, 0.7 mmol) at 0 °C and the mixture was stirred at 0 °C for 1 hour. LCMS showed starting material was consumed and a peak with the desired mass was formed. The reaction mixture was washed with water (20 mL x 2) and brine (20 mL x 2). The organic fraction was dried over anhydrous Na_2_SO_4_, filtered and concentrated under reduced pressure. The residue was purified by preparative TLC (1:1 petroleum ether/EtOAc) followed by HPLC (Column: Kromasil-C18 100 x 21.2 mm, 5 µm; Mobile Phase: MeCN/H_2_O (0.1% formic acid); Flowrate: 25 mL/min; Wavelength: 214 nm) to afford **10** (52 mg, 59% yield) as a brown solid. ^1^H NMR (400 MHz, DMSO-d_6_) δ = 10.17 (s, 1H), 8.63 (t, *J* = 6.1 Hz, 1H), 7.44 (d, *J* = 8.1 Hz, 2H), 7.29 (d, *J* = 8.1 Hz, 2H), 7.25 (d, *J* = 1.2 Hz, 1H), 6.82 (d, *J* = 1.2 Hz, 1H), 6.64 (td, *J* = 6.6, 15.4 Hz, 1H), 6.31 (d, *J* = 15.4 Hz, 1H), 4.39 (d, *J* = 6.1 Hz, 2H), 4.13 (s, 1H), 3.81 (s, 3H), 3.62 (d, *J* = 5.4 Hz, 2H), 2.64 - 2.53 (m, 6H). MS: [M+H]^+^ 365.2

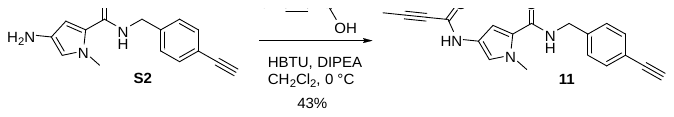

***4*-(but-2-ynamido)-*N*-(4-ethynylbenzyl)-1-methyl-1*H*-pyrrole-2-carboxamide (11):** To a mixture of but-2-ynoic acid (24.9 mg, 0.296 mmol), 4-amino-*N*-(4-ethynylbenzyl)-1-methyl-1*H*-pyrrole-2-carboxamide (50 mg, 0.20 mmol) and HBTU (112 mg, 0.296 mmol) in CH_2_Cl_2_ (10 mL) was added DIPEA (63.8 mg, 0.493 mmol) at 0 °C and the mixture was stirred at 0 °C for 1 hour. LCMS indicated starting material was consumed and a peak with the desired mass was formed. The mixture was washed with water (20 mL x 2) and brine (20 mL), dried over anhydrous Na_2_SO_4_, filtered and concentrated under reduced pressure. The crude product was purified by preparative TLC (1:1, petroleum ether/EtOAc) to give product that was further purified by HPLC (Column: Xbridge-C18 150 x 19 mm, 5 µm; Mobile Phase: 32-42% MeCN/H_2_O (0.1% formic acid); Flowrate: 20 mL/min; Wavelength: 214 nm) to afford **11** (31 mg, 43% yield) as an off-white solid. ^1^H NMR (400 MHz, DMSO-d_6_) δ = 10.55 (s, 1H), 8.64 (t, *J* = 6.0 Hz, 1H), 7.43 (d, *J* = 8.1 Hz, 2H), 7.28 (d, *J* = 8.1 Hz, 2H), 7.12 (s, 1H), 6.79 (s, 1H), 4.36 (d, *J* = 6.1 Hz, 2H), 4.13 (s, 1H), 3.78 (s, 3H), 2.01 (s, 3H). MS: [M+H]^+^ 320.1

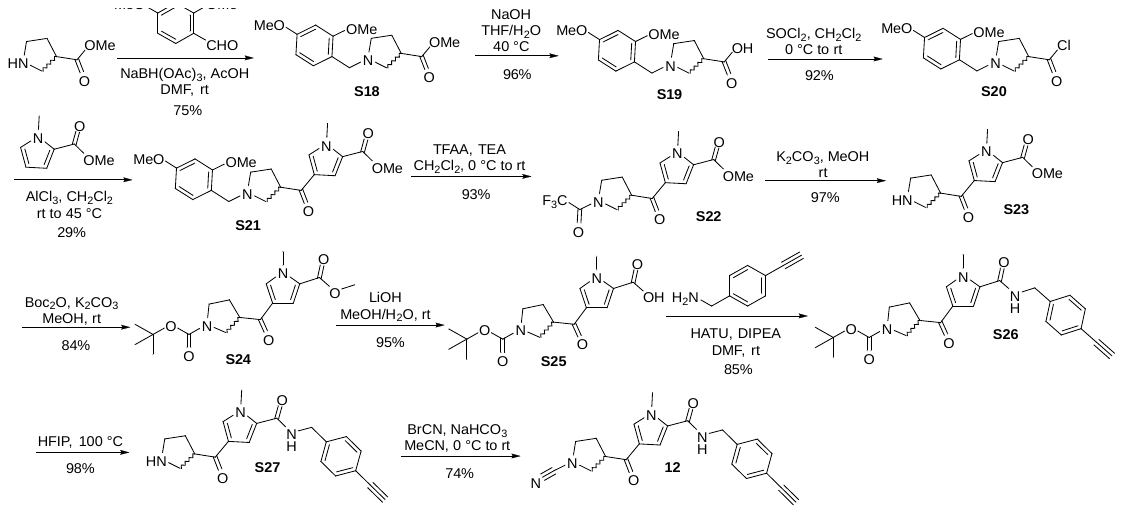

**Methyl 1-(2,4-dimethoxybenzyl)pyrrolidine-3-carboxylate (S18)**: To a mixture of methyl pyrrolidine-3-carboxylate hydrochloride (7.00 g, 42.3 mmol) and 2,4-dimethoxybenzaldehyde (7.02 g, 42.3 mmol) in DMF (150 mL) was added AcOH (2.54 g, 42.3 mmol) and NaBH(OAc)_3_ (26.9 g, 127 mmol), then the mixture was stirred at 25 °C for 16 hours. LCMS showed starting material was consumed and a peak with the desired mass was formed. The mixture was quenched with water (200 mL) and adjusted to ~pH 8 by the addition of saturated Na_2_CO_3_, then the mixture was extracted by EtOAc (100 mL x 2). The organic phase was washed with brine, dried over anhydrous Na_2_SO_4_, filtered, and concentrated under reduced pressure. The residue was purified by silica gel chromatography (0-20% MeOH/CH_2_Cl_2_) to afford **S18** (8.8 g, 75%) as a yellow oil. ^1^H NMR (400 MHz, CDCl_3_) δ 7.21 (d, *J* = 8.3 Hz, 1H), 6.48 – 6.43 (m, 2H), 3.80 (s, 3H), 3.80 (s, 3H), 3.68 (s, 3H), 3.63 (s, 2H), 3.06 – 2.95 (m, 2H), 2.80 – 2.74 (m, 1H), 2.65 (dd, *J* = 9.0, 7.3 Hz, 1H), 2.59 – 2.50 (m, 1H), 2.13 – 2.05 (m, 2H). MS: [M+H]^+^ 280.2

**1-(2,4-dimethoxybenzyl)pyrrolidine-3-carboxylic acid (S19)**: To a mixture of methyl 1-(2,4 dimethoxybenzyl)pyrrolidine-3-carboxylate (8.80 g, 31.5 mmol) in THF (80 mL) and H_2_O (80 mL) was added NaOH (3.8 g, 95 mmol), then the mixture was stirred at 40 °C for 2 hours. LCMS showed starting material was consumed and a peak with the desired mass was formed. The solvent of THF was removed, and the mixture was acidized by the addition of 6 N HCl to ~pH 3, then concentrated under reduced pressure. The residue was dissolved with CH_2_Cl_2_/MeOH (100 mL/10 mL) and filtered through Celite. The filter cake was washed with CH_2_Cl_2_/MeOH (50 mL/5 mL), then the filtrate was concentrated to afford **S19** (8 g, 96%) as an off-white solid, which was used directly without any further purification. ^1^H NMR (400 MHz, D_2_O) δ 7.47 (d, *J* = 8.4 Hz, 1H), 6.65 (s, 1H), 6.62 (d, *J* = 8.4 Hz, 1H), 4.38 – 4.35 (m, 2H), 3.84 (s, 3H), 3.81 (s, 3H), 3.72 3.63 (m, 1H), 3.58 – 3.53 (m, 1H), 3.46 – 3.39 (m, 1H), 3.34 – 3.21 (m, 2H), 2.41 – 2.19 (m, 2H). MS: [M+H]^+^ 266.2

**1-(2,4-dimethoxybenzyl)pyrrolidine-3-carbonyl chloride (S20)**: To a mixture of 1-(2,4-dimethoxybenzyl) pyrrolidine-3-carboxylic acid (7.5 g, 28.3 mmol) in CH_2_Cl_2_ (200 mL) was added SOCl_2_ (6.16 mL, 84.9 mmol) at 0 °C dropwise over 10 minutes, then the mixture was stirred at 25 °C for 16 hours. LCMS showed starting material was consumed and a peak with the desired mass was formed. The mixture was concentrated to afford **S20** (7.34 g, 91.5%) as a dark gum. The gum was used directly without any further purification. ^1^H NMR (400 MHz, CDCl_3_) δ 7.47 (d, *J* = 8.4 Hz, 1H), 6.56 (dd, *J* = 8.4, 2.3 Hz, 1H), 6.49 (d, *J* = 2.3 Hz, 1H), 4.26 (d, *J* = 5.0 Hz, 2H), 4.06 – 3.92 (m, 2H), 3.88 (s, 3H), 3.83 (s, 3H), 3.72 – 3.63 (m, 1H), 3.34 – 3.28 (m, 1H), 3.05 – 2.93 (m, 1H), 2.84 – 2.71 (m, 1H), 2.48 – 2.36 (m, 1H). MS: [M+H]^+^ 280.2 (quenched by MeOH)

**Methyl 4-(1-(2,4-dimethoxybenzyl)pyrrolidine-3-carbonyl)-1-methyl-1*H*-pyrrole-2-carboxylate (S21)**: To a mixture of AlCl_3_ (5.75 g, 43.1 mmol) in CH_2_Cl_2_ (500 mL) was added 1-(2,4-dimethoxybenzyl) pyrrolidine-3-carbonyl chloride (7.3 g, 25.8 mmol) at 0 °C under N_2_. The mixture was stirred at 0 °C for 30 minutes and warmed to room temperature. After that, methyl 1-methyl-1*H*-pyrrole-2-carboxylate (3.0 g, 21.6 mmol) and additional AlCl_3_ (2.87 g, 21.52 mmol) was added. Then the mixture was stirred at 45 °C for 30 minutes and cooled to room temperature, and additional AlCl_3_ (2.87 g, 21.52 mmol) was added at 25 °C. Then the mixture was stirred at 45 °C for 3 hours. LCMS showed starting material was consumed and a peak with the desired mass was formed. The reaction mixture was cooled to room temperature and poured into ice-water (~600 mL). The mixture was basified by the addition of 2 N NaOH to ~pH 8, then the mixture was extracted with CH_2_Cl_2_ (400 mL x 2). The organic phase was washed with brine (100 mL), dried over anhydrous Na_2_SO_4_, filtered, and concentrated in vacuo. The residue was by silica gel chromatography (0-8% MeOH/CH_2_Cl_2_) to afford **S21** (2.4 g, 28.8%) as a yellow solid. ^1^H NMR (400 MHz, CD_3_CN) δ 7.63 (s, 1H), 7.49 (dd, *J* = 8.0, 1.6 Hz, 1H), 7.26 (d, *J* = 2.0 Hz, 1H), 6.60-6.52 (m, 2H), 4.21 (s, 2H), 3.91 (s, 3H), 3.85 (s, 3H), 3.81 (s, 3H), 3.79 (s, 3H), 3.38 – 3.30 (m, 2H), 3.22 – 3.16 (m, 1H), 3.03 – 2.96 (m, 1H), 2.12 – 2.04 (m, 1H) (2H missing are under water peak). MS: [M+H]^+^ 387.2

**Methyl 1-methyl-4-(1-(2,2,2-trifluoroacetyl)pyrrolidine-3-carbonyl)-1*H*-pyrrole-2-carboxylate (S22)**: To a mixture of methyl 4-(1-(2,4-dimethoxybenzyl)pyrrolidine-3-carbonyl)-1-methyl-1*H*-pyrrole-2-carboxylate (500 mg, 1.29 mmol) in CH_2_Cl_2_ (15 mL) was added and Et_3_N (288 mg, 2.85 mmol) and TFAA (543 mg, 2.59 mmol) at 0 °C, then the mixture was stirred at 25 °C for 2 hours. LCMS showed starting material was consumed and a peak with the desired mass was formed. The reaction was quenched with water (10 mL). The mixture was extracted with CH_2_Cl_2_ (15 mL x 2), and the combined organic layers were washed with brine (10 mL x 2), dried over anhydrous Na_2_SO_4_, filtered, and concentrated in vacuo. The residue was purified silica gel chromatography (0-40% petroleum ether/EtOAc) to afford **S22** (400 mg, 93%) as a light-yellow gum. ^1^H NMR (400 MHz, CDCl_3_) δ 7.47 – 7.40 (m, 1H), 7.32 (t, *J* = 2.2 Hz, 1H), 3.99 and 3.98 (2 s, 3H, rotamers), 3.86 and 3.85 (2 s, 3H, rotamers), 3.80 – 3.63 (m, 5H), 2.41 – 2.13 (m, 2H). ^19^F NMR (400 MHz, CDCl_3_) δ -72.42 and -72.45 (2 s, 1F, rotamers). MS: [M+Na]^+^ 355.1

**Methyl 1-methyl-4-(pyrrolidine-3-carbonyl)-1*H*-pyrrole-2-carboxylate (S23)**: To a mixture of methyl 1-methyl-4-(1-(2,2,2-trifluoroacetyl) pyrrolidine-3-carbonyl)-1*H*-pyrrole-2-carboxylate (400 mg, 1.20 mmol) in MeOH (10 mL) was added K_2_CO_3_ (250 mg, 1.81 mmol) at 25 °C, then the mixture was stirred at 25 °C for 1 hour. LCMS showed starting material was consumed and a peak with the desired mass was formed. The mixture was filtered and directly concentrated to afford **S23** (275 mg, 96.7%) as a white gum, which was used directly without any further purification. MS: [M+H]^+^ 237.2

**Methyl 4-(1-(*tert*-butoxycarbonyl)pyrrolidine-3-carbonyl)-1-methyl-1*H*-pyrrole-2-carboxylate (S24)**: To a mixture of methyl 1-methyl-4-(pyrrolidine-3-carbonyl)-1*H*-pyrrole-2-carboxylate (275 mg, 1.16 mmol) in MeOH (10 mL) was added K_2_CO_3_ (483 mg, 3.49 mmol) and (Boc)_2_O (508 mg, 2.33 mmol) at 25 °C, then the mixture was stirred at 25 °C for 2 hours. LCMS showed starting material was consumed and a peak with the desired mass was formed. The mixture was filtered and concentrated under reduced pressure. The residue was purified by silica gel chromatography (0-80% petroleum ether/EtOAc) to afford **S24** (330 mg, 84%) as a white gum. ^1^H NMR (400 MHz, CDCl_3_) δ 7.42 (s, 1H), 7.33 (s, 1H), 3.97 (s, 3H), 3.85 (s, 3H), 3.67 – 3.49 (m, 4H), 3.43 – 3.36 (m, 1H), 2.22 – 2.12 (m, 2H), 1.46 (s, 9H). MS: [M+Na]^+^ 359.2

**4-(1-(*tert*-butoxycarbonyl)pyrrolidine-3-carbonyl)-1-methyl-1*H*-pyrrole-2-carboxylic acid (S25)**: To a mixture of methyl 4-(1-(*tert*-butoxycarbonyl)pyrrolidine-3-carbonyl)-1-methyl-1*H*-pyrrole-2-carboxylate (330 mg, 0.982 mmol) in MeOH (10 mL) and H_2_O (10 mL) was added LiOH.H_2_O (123 mg, 2.94 mmol) at 25 °C. Then the reaction mixture was stirred at 25 °C for 3 hours. LCMS showed starting material was consumed and a peak with the desired mass was formed. The mixture was concentrated to remove the solvent of MeOH and acidified by the addition of 2 N HCl to ~pH 3, then the mixture was extracted by EtOAc (20 mL x 2). The combined organic layers were washed with brine (15 mL), dried over anhydrous Na_2_SO_4_, filtered, and concentrated to afford **S25** (300 mg, 95%) as a light-yellow solid. ^1^H NMR (400 MHz, CDCl_3_) δ 7.45 (s, 1H), 7.43 (s, 1H), 3.98 (s, 3H), 3.66 – 3.58 (m, 4H), 3.44 – 3.38 (m, 1H), 2.12 – 2.01 (m, 2H), 1.46 (s, 9H). MS: [M+Na]^+^ 345.1

***t*ert-butyl-3-(5-((4-ethynylbenzyl)carbamoyl)-1-methyl-1*H*-pyrrole-3-carbonyl)pyrrolidine-1-carboxylate (S26)**: To a mixture of 4-(1-(*tert*-butoxycarbonyl)pyrrolidine-3-carbonyl)-1-methyl-1*H*-pyrrole-2-carboxylic acid (410 mg, 1.27 mmol), (4-ethynylphenyl)methanamine (184 mg, 1.40 mmol) and HATU (725 mg, 1.91 mmol) in DMF (10 mL) was added DIPEA (493 mg, 3.82 mmol), then the mixture was stirred at 25 °C for 2 hours. LCMS showed starting material was consumed and a peak with the desired mass was formed. The reaction was quenched with water (25 mL) and extracted with EtOAc (20 mL x 2). The combined organic layers were washed with water (20 mL x 2) and brine (15 mL x 2), dried over anhydrous Na_2_SO_4_, filtered, and concentrated under reduced pressure. The crude product was purified silica gel chromatography (0-75% petroleum ether/EtOAc) to afford the **S26** (470 mg, 85%) as a yellow solid. ^1^H NMR (400 MHz, CDCl_3_) δ 7.47 (d, *J* = 8.1 Hz, 2H), 7.36 (s, 1H), 7.29 (d, *J* = 8.0 Hz, 2H), 7.02 (s, 1H), 6.43 (t, *J* = 5.5 Hz, 1H), 4.57 (d, *J* = 5.9 Hz, 2H), 4.00 (s, 3H), 3.70 – 3.49 (m, 4H), 3.41 – 3.35 (m, 1H), 3.08 (s, 1H), 2.25 – 2.16 (m, 1H), 2.13 – 2.06 (m, 1H), 1.45 (s, 9H). MS: [M+Na]^+^ 458.2

***N*-(4-ethynylbenzyl)-1-methyl-4-(pyrrolidine-3-carbonyl)-1*H*-pyrrole-2-carboxamide (S27)**: A mixture of *tert*-butyl 3-(5-((4-ethynylbenzyl) carbamoyl)-1-methyl-1*H*-pyrrole-3-carbonyl) pyrrolidine-1-carboxylate (460 mg, 1.06 mmol) in HFIP (25 mL) was stirred at 100 °C for 32 hours in a 50 mL sealed tube. LCMS showed starting material was consumed and a peak with the desired mass was formed. The mixture was concentrated to afford **S27** (345 mg, 97.4%) as a yellow solid, which was used directly without any further purification. MS: [M+H]^+^ 336.2

***4*-(1-cyanopyrrolidine-3-carbonyl)-*N*-(4-ethynylbenzyl)-1-methyl-1*H*-pyrrole-2-carboxamide (12)**: To a mixture of *N*-(4-ethynylbenzyl)-1-methyl-4-(pyrrolidine-3-carbonyl)-1*H*-pyrrole-2-carboxamide (340 mg, 1.01 mmol) and NaHCO_3_ (255 mg, 3.04 mmol) in MeCN (10 mL) was added BrCN (268 mg, 2.53 mmol) at 0 °C. The mixture was warmed to room temperature and stirred for 16 hours. LCMS showed starting material was consumed and a peak with the desired mass was formed. The reaction was quenched with water (20 mL) and the mixture was extracted with EtOAc (30 mL x 2). The organic fractions were combined, washed with brine (20 mL), dried over anhydrous Na_2_SO_4_, filtered, and then concentrated under reduced pressure. The crude product was purified by silica gel chromatography (0-80% petroleum ether/EtOAc) to afford the **12** (270 mg, 74% yield) as an off-white solid. The mixture of enantiomers (158 mg, 0.438 mmol) was purified by SFC (SFC 150; Column: Daicel CHIRALCEL IC, 250 mm x 30 mm I.D., 10 μm; Mobile Phase: CO_2_/MeOH [0.2% NH_3_ (7 M Solution in MeOH)] = 55/45; Flowrate: 1.5 g/min; Wavelength: UV 214 nm; Temperature: 35 °C) to afford **enantiomer 1** (69.5 mg, 44% yield, 100%) as a white solid and **enantiomer 2** (70.1 mg, 44% yield, 100%) as a white solid. Analytical conditions for purity analysis by SFC (Acquity UPC; Column: Daicel CHIRALPAK IC-3 3 mm*150 mm, 3 μm; Mobile phase: CO_2_/MeOH (0.1%DEA) = 55/45; Flowrate: 1.5 mL/min; UV 215 nm and 254 nm; Temperature: 37 °C. ^1^H NMR (400 MHz, DMSO-d_6_) δ = 8.85 (t, *J* = 6.2 Hz, 1H), 7.86 (s, 1H), 7.44 (d, *J* = 8.1 Hz, 2H), 7.39 – 7.24 (m, 3H), 4.40 (d, *J* = 5.9 Hz, 2H), 4.14 (s, 1H), 3.88 (s, 3H), 3.79 (td, *J* = 7.3, 14.3 Hz, 1H), 3.61 (t, *J* = 8.6 Hz, 1H), 3.51 – 3.47 (m, 1H), 3.42 – 3.39 (m, 2H), 2.23 – 2.15 (m, 1H), 2.03 – 1.94 (m, 1H). 13C NMR (100 MHz, DMSO-d6) δ = 160.7, 140.7, 132.9, 131.7, 127.4, 127.2, 121.6, 120.1, 117.2, 112.3, 83.4, 80.5, 51.9, 50.01, 45.9, 41.7, 37.0, 29.4. HRMS C_21_H_21_N_4_O_2_ [M+H]^+^ calculated: 361.1659; found: 361.1654

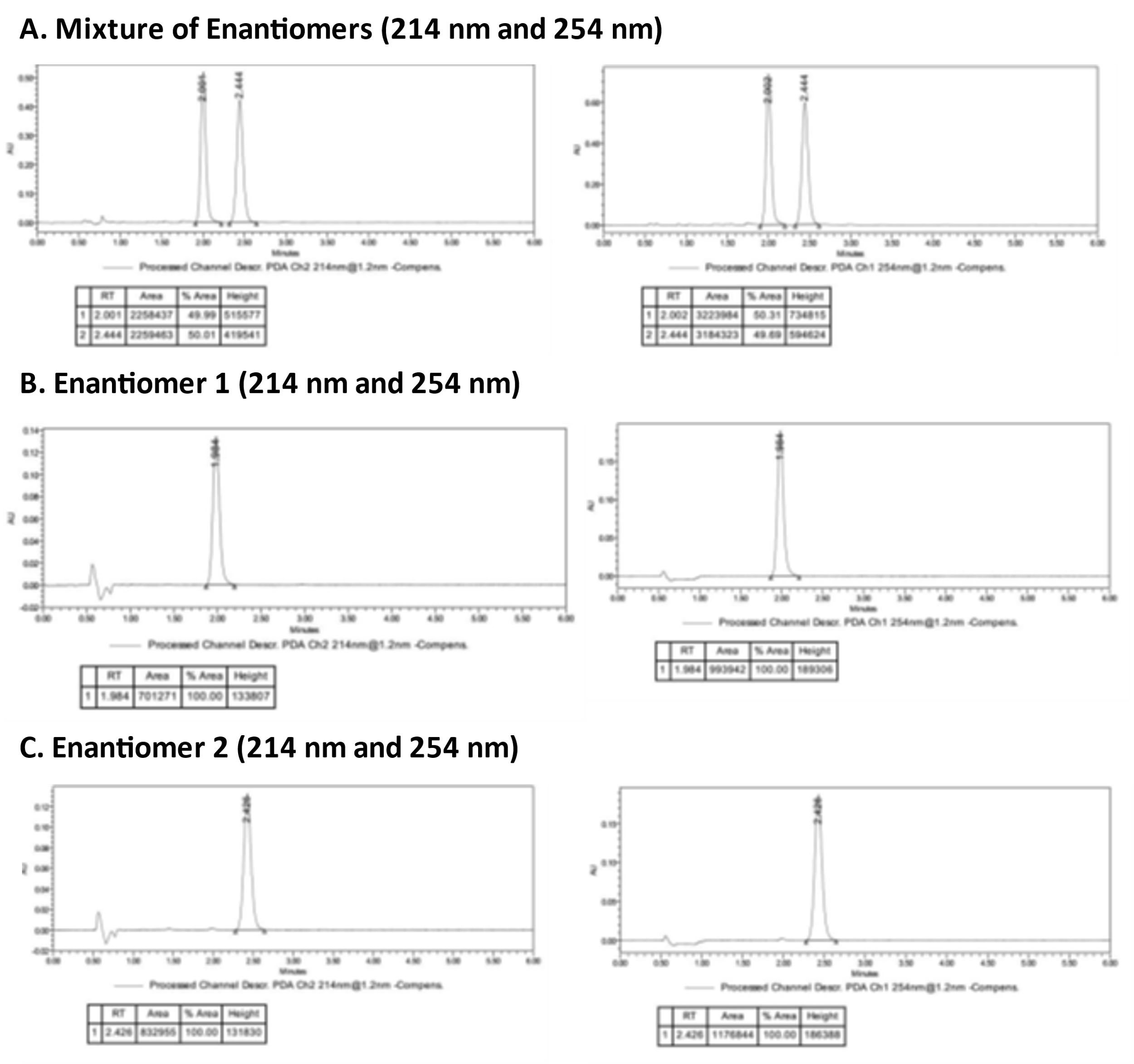

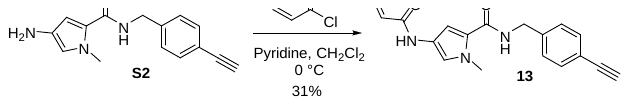

**4-acrylamido-*N*-(4-ethynylbenzyl)-1-methyl-1*H*-pyrrole-2-carboxamide (13):** To a solution of 4-amino-*N*-(4-ethynylbenzyl)-1-methyl-1*H*-pyrrole-2-carboxamide (50 mg, 0.20 mmol) in CH_2_Cl_2_ (10 mL) was added acryloyl chloride (26.8 mg, 0.296 mmol) and pyridine (31.2 mg, 0.395 mmol) at 0 °C and the mixture was stirred at 0°C for 1 hour. LCMS indicated starting material was consumed and a peak with the desired mass was formed. The mixture was washed with water (20 mL x 2) and brine (20 mL), dried over anhydrous Na_2_SO_4_, filtered and concentrated under reduced pressure. The crude product was purified by preparative TLC (1:1, petroleum ether/EtOAc) to give product that was further purified by HPLC (Column: Xbridge-C18 150 x 19 mm, 5 µm; Mobile Phase: 38-42% MeCN/H_2_O (0.1% formic acid); Flowrate: 20 mL/min; Wavelength: 214 nm) to afford **13** (19 mg, 31% yield) as an off-white solid. ^1^H NMR (400 MHz, DMSO-d_6_) δ = 10.07 (s, 1H), 8.63 (t, *J* = 6.1 Hz, 1H), 7.43 (d, *J* = 8.1 Hz, 2H), 7.29 (d, *J* = 8.1 Hz, 2H), 7.24 (s, 1H), 6.82 (s, 1H), 6.36 (dd, *J* = 10.0, 16.9 Hz, 1H), 6.17 (dd, *J* = 2.0, 16.9 Hz, 1H), 5.65 (dd, *J* = 2.0, 10.0 Hz, 1H), 4.39 (d, *J* = 5.9 Hz, 2H), 4.13 (s, 1H), 3.81 (s, 3H). MS: [M+H]^+^ 308.1

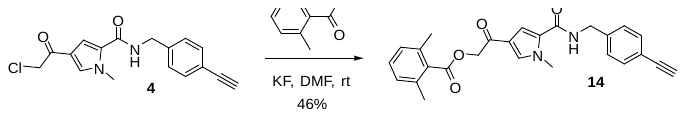

**2-(5-((4-ethynylbenzyl) carbamoyl)-1-methyl-1*H*-pyrrol-3-yl)-2-oxoethyl 2,6-dimethylbenzoate (14):** 4-(2-chloroacetyl)-*N*-(4-ethynylbenzyl)-1-methyl-1*H*-pyrrole-2-carboxamide (50 mg, 0.16 mmol) was treated with potassium fluoride (27.7 mg, 0.477 mmol) in DMF (3 mL), followed by 2,6-trifluorobenzoic acid (26.2 mg, 0.175 mmol). The reaction mixture was stirred at 25 °C for 3 hours. LCMS indicated starting material was consumed and a peak with the desired mass was formed. The mixture was poured over water (10 mL) and extracted with EtOAc (15 mL x 2). The combined organic fraction was washed with brine, dried over anhydrous Na_2_SO_4_, filtered, and concentrated under reduced pressure. The crude product was purified by HPLC (Column: Xbridge-C18 150 x 19 mm, 5 µm; Mobile Phase: MeCN/H_2_O (0.1% formic acid); Flowrate: 20 mL/min; Wavelength: 214 nm) to afford **14** (31 mg, 46% yield) as a white solid. ^1^H NMR (400 MHz, DMSO-d_6_) δ = 8.87 (t, *J* = 5.9 Hz, 1H), 7.92 (d, *J* = 1.6 Hz, 1H), 7.45 (d, *J* = 8.2 Hz, 2H), 7.38 (d, *J* = 2.0 Hz, 1H), 7.34 - 7.25 (m, 3H), 7.12 (d, *J* = 7.4 Hz, 2H), 5.38 (s, 2H), 4.42 (d, *J* = 5.9 Hz, 2H), 4.14 (s, 1H), 3.91 (s, 3H), 2.34 (s, 6H). MS: [M+H]^+^ 429.1

3. **Analysis for selection of cell lines**

The selection of cell lines were based on ranking the gene expression values obtained from human protein atlas  (“RNA HPA cell line gene data”, [https://www.proteinatlas.org/about/download](https://urldefense.com/v3/__https:/www.proteinatlas.org/about/download__;!!H9nueQsQ!px4hGpGISY9d8h2keYuk-MzJCRW9EVc-M5PxkV3C9Yi9GU2rWbm-S2sm4cGck6v1$)), and cross-referencing this list with Sigma top 100 cell lines for the convenience of ordering. The lowest 25% of cells lines for total DUB RNA expression were filtered out due to an assumption that these would not possess enough sensitivity for proteomics studies. The number of cell lines selected was based on “greedy” algorithm, which consists of iteratively adding each cell line that would induce the most new DUB protein identifications compared with the previous pool. We hypothesized that a combination of three cell lines, an osteosarcoma (U2OS), glioblastoma (U87-MG) and a breast cancer cell line (T47D) would allowed identification of >80% of the DUBome by proteomics.

4. **Live cell protein labelling profiling of probes using click chemistry, SDS-PAGE and in-gel fluorescence**
a. Live cell probe incubation

For each experiment, sterile treated 12-well plates (1 mL media working volume) were seeded with either U2OS, U87-MG or T47D cells in McCoy 5a, low glucose DMEM or RPMI-1680 media, respectively, containing 10% FBS, and incubated at 5% CO_2_ and 37◦C. After 24h, when cells had achieved 90–100% confluency, plates were treated with either DMSO vehicle (0.1% (v/v)) or 0.01, 0.1, 1 and 10 µM of probe, and incubated at 37 °C for 2 h. All compound treatments were performed with a 1000× DMSO stock of each probe which was added directly to well with gentle mixing.

The cells were relieved of media, washed twice with PBS (0.5mL), and then lysed with lysis buffer (50  µL (1% (v/v) Triton X-100, 0.1% (w/v) sodium dodecyl sulfate (SDS), EDTA-free complete protease inhibitor cocktail (1×, Roche) in PBS on ice for 10 min. The lysates were scraped and transferred to corresponding Lo-Bind Eppendorfs. Each lysate was sonicated for 1 minute, and the samples centrifuges at 5,000 x g at 4 ˚C for 5 minutes to pellet insoluble cellular debris. The supernatant was collected and protein concentration determined using the DC Protein Assay (Bio-Rad) as per manufacturer’s instructions, and normalised to lowest concentrated sample (e.g. 1 mg/mL, using lysis buffer).

b. Click chemistry and precipitation

The desired amount of lysed protein from each sample was made up to 0.5–2 mg mL-1 with PBS to a total volume of ≤300 µL. The following “click mixture” was prepared separately, preparing 6 µL for every 100 µL of lysate:

1 µL Azide TAMRA click reagent (AzT, 10 mM in DMSO, 1 vol; final concentration in reaction 0.1 mM),

2 µL CuSO_4_ (50 mM in H_2_O, 2 vol; final concentration in reaction 1 mM),

2 µL Tris(2-carboxyethyl)phosphine (TCEP, 50 mM in H_2_O, 2 vol; final concentration in reaction 1 mM),

1 µL Tris(benzyltriazolylmethyl)amine (TBTA, 10 mM in DMSO, 1 vol; final concentration in reaction 0.1 mM).

The click mixture was vortexed and incubated at rt for 2 min before 6 µL of the mixture was added to every 100 µL of lysate. The reaction mixtures were shaken at rt for 1 h before being quenched with EDTA (500 mM in H2O) to a final concentration of 10 mM.

A table-top centrifuge was pre-chilled to 4 °C. Proteins were precipitated by adding H_2_O (1 vol), MeOH (2 vol) and CHCl_3_ (0.5 vol), vortexing briefly then centrifuging at 17,000 × g for 5 min. The top H_2_O/MeOH layer was discarded and the middle layer of protein pellet and lower CHCl_3_ layer retained. The suspension was washed with MeOH (300 µL), and sonicated to break up the pellet. The proteins were pelleted by centrifugation at 10,000–17000 × g for 5–10 min or until a compact pellet was formed. The protein pellet was washed once more with MeOH (300 µL). The MeOH was decanted and the pellet air-dried for 1 min. The pellet was resuspended by sonicating and completely dissolving in 1% (w/v) SDS in PBS (to 5 mg/mL protein) before being made up to 1 mg/mL protein with PBS.

c. SDS-PAGE and In-gel fluorescence

10- or 15-well SDS-polyacrylamide gels with a 12% resolving gel and 4% stacking gel were used for all gel electrophoresis experiments. All gels were run using a Bio-Rad Mini-PROTEIN® Tetra Cell with a Bio-Rad PowerPac™ Basic power supply. In general, 10 µg of protein was run per well in a volume of 10 µL.

Samples were prepared by adding 5 µL of 4× loading buffer (1:4 β-mercaptoethanol:5× NuPAGE LDS sample buffer) to 10 µL of sample and boiling at 95 °C for 10 min. The samples were briefly centrifuged then 10 µL of each sample was added to a well of the gel, with at least one well also containing Precision Plus Protein™ All Blue Prestained Protein Standard (3.5 µL, Bio-Rad). The gels were run in running buffer (25 mM trizma base, 194 mM glycine, 1% (w/v) SDS) for 15 min at 90 V then up to 1 h 15 min at 150 V. The fluorescence on the gel was detected using a Typhoon™ FLA 9500 biomolecular imager (GE Healthcare Life Sciences) detecting TAMRA fluorescence, and the contrast normalized using Fiji (ImageJ) software. The protein loading was verified by staining with Coomassie Brilliant Blue. Coomassie stained gels were imaged using the digitization method (trans-illumination) on an ImageQuant LAS-4000 Imaging System (Fujifilm) and the contrast normalized in Fiji.

5. **Chemical Proteomic profiling of probes in live cells**
a. Live cell probe incubation

For each experiment, sterile treated 10 cm dishes (10 mL media working volume) were seeded with either U2OS, U87-MG or T47D cells in McCoy 5a, low glucose DMEM or RPMI-1680 media, respectively, containing 10% FBS, and incubated at 5% CO_2_ and 37◦C. After 24h, when cells had achieved 90–100% confluency, plates were treated with either DMSO vehicle (0.1% (v/v)) or 0.3 or 3 µM of probes **1-14**, and incubated at 37 °C for 2 h. All compound treatments were performed with a 1000× DMSO stock of each probe which was added directly to dish with gentle mixing.

The cells were relieved of media, washed twice with PBS (5mL), and then lysed with lysis buffer (500 µL (1% (v/v) Triton X-100, 0.1% (w/v) sodium dodecyl sulfate (SDS), EDTA-free complete protease inhibitor cocktail (1×, Roche) in PBS on ice for 10 min. The lysates were scraped and transferred to corresponding Lo-Bind Eppendorfs. Each lysate was sonicated for 1 minute, and the samples centrifuges at 5,000 x g at 4 ˚C for 5 minutes to pellet insoluble cellular debris. The supernatant was collected and protein concentration determined using the DC Protein Assay (Bio-Rad) as per manufacturer’s instructions, and normalised to lowest concentrated sample (e.g. 1 mg/mL, using lysis buffer).

b. Click chemistry and precipitation

Click chemistry and precipitation steps were carried out as described in section 4b, on at least 400ug of protein per sample. The pellet was resuspended by sonicating and completely dissolving in 1% (w/v) SDS in HEPES 50 mM pH 8.0, before being made up to 1 mg/mL protein with HEPES 50 mM pH 8.0 to achieve a final concentration of 0.2% SDS. Samples were centrifuged for 5 min at maximum speed at room temperature to make sure no pellet formed and protein was fully dissolved.

c. Neutravidin biotin enrichment and on-bead digestion

Pull down of biotinylated proteins was achieved by incubating samples with pre-equilibrated (3x 1 mL washes with 0.2% (w/v) SDS in HEPES 50 mM pH 8 buffer) Pierce™ NeutrAvidin™ Agarose beads for 2 hours at room temperature. NeutrAvidin™ Agarose beads were used in ratio of 1 µL bead resin per 10ug of protein sample. The beads were subsequently washed by moderate shaking for 10 seconds then briefly pelleting by table-top centrifuge then vacuum aspirating the supernatant with fine-end pipette tips (to not disturb the agarose). Proteins were reduced and alkylated with 5 mM TCEP and 10 mM chloroacetamide in 100 µL 50 mM HEPES with moderate shaking for 10 min at rt. Proteins were digested on-bead by treatment with 2.5 µL trypsin or trypsin/LysC mix(Promega, 20 ug dissolved in 100 µL 50 mM HEPES) with vigorous shaking at 37 °C overnight. The trypsin reaction was quenched by adding 1x EDTA-free protease inhibitor (50X stock,). The beads were pelleted and the supernatant (150 µL) transferred to a new epppendorf tube. An extra bead wash (50 µL, HEPES 50 mM pH 8.0) was combined with previous supernatant (200 µL total). At this stage, 10 µL of sample was analysed using a Pierce™ Quantitative Fluorometric Peptide Assay (Catalog number: 23290) as per the manufacturers instructions to determine accurate an peptide amount for TMT labelling.

d. Isobaric TMT labelling and high pH reverse phase fractionation

Upon checking that the pH of each sample was ~8. approximately 10 ug of peptide sample was labelled with 1/10^th^ of an 0.8 mg vial of the appropriate TMT10plex™ Isobaric Mass Tag Labelling Reagent (Thermo Scientific) dissolved in acetonitrile (40 µL) with moderate shaking for 2 h at rt TMT-labelling was quenched by the addition of 1 µL of 5% (w/v) hydroxylamine and the samples from each TMT set were combined to form a “multi-plex” solution. Trifluoroacetic acid was added to achieve a 1% v/v solution and these samples were evaporated to dryness. Samples were then fractionated using the Pierce High pH Reversed-Phase Peptide Fractionation Kit (Catalog number: 84868) as per the manufacturers instructions. All fractions of each sample were collected in low-bind epppendorfs tubes, evaporated to dryness and stored at –80 °C.

e. LC-MS/MS analysis

Samples were rehydrated in 0.5% (v/v) formic acid, 2% (v/v) UPLC grade MeCN in Optima™ LC/MS H_2_O (Fisher Scientific) and dissolved completely by vortexing and sonication. Samples were filtered through 3x Durapore® membrane filters (Millipore) plugged into a p20 pipette tip by centrifuging the samples through the filters (4000 × g, 5 min) into a mass spectrometry vial. Samples were stored at 4 °C until ready for analysis.

Peptides were separated on an EASY-Spray™ Acclaim PepMap C18 column (50 cm × 75 µm inner diameter, Thermo Fisher Scientific) using a 3-hour linear gradient separation of 0–100% solvent B (80% MeCN supplemented with 0.1% formic acid): solvent A (2% MeCN supplemented with 0.1% formic acid) at a flow rate of 250 nL/min. The liquid chromatography was coupled to a Q-Exactive mass spectrometer via an easy-spray source (Thermo Fisher Scientific) which operated in data-dependent mode with survey scans acquired at a resolution of 70,000 at m/z 200. Scans were acquired from 350 to 1800 *m/z*. Up to 10 of the most abundant isotope patterns with charge +2 or higher from the survey scan were selected with an isolation window of 1.6 m/z and fragmented by HCD with normalized collision energy of 25. The maximum ion injection times for the survey scan and the MS/MS scans (acquired with a resolution of 35,000 at *m/z* 200) were 20 and 120 ms, respectively. The ion target value for MS was set to 106 and for MS/MS to 105, and the intensity threshold was set to 8.3 × 10^2^.

f. Database searching and proteomics data analysis

Peptide searches were performed in MaxQuant (version 1.6.10.43). Under group-specific parameters and type, reporter ion MS2 was selected, and the appropriate TMT10plex™ isobaric labels selected for both lysines and N-termini. The isotope errors contained in each TMT batch code was also entered. For all experiments, oxidation (M) and acetyl (protein N-term) were set as variable modifications, carbamidomethyl (C) was set as a fixed modification, trypsin/P was set as the digestion mode. Where multiple TMT sets were analysed, re-quantify and match between runs were selected, and latest UniProt FASTA files for the human proteome and contaminants databases were used.

Data analysis was performed in Perseus version 1.6.6.0. Reporter intensity corrected values were loaded into the matrix. Data was filtered by removing rows based on “reverse”, and “potential contaminant” columns. Data were log2 transformed and filtered by row, retaining those that had 2 valid values in each triplicate condition. To account for variance in protein abundance across different sample, the median of each channel was subtracted from each protein. If appropriate, multiple TMT data sets were normalized by subtracting the mean of each row within each TMT “plex”. The log2 fold enrichment for each probe was determined by subtracting the DMSO control value from each of the different probe treated conditions.

6. **Biochemical in vitro inhibition profiling of probes**

Biochemical DUB inhibition assays were performed at Ubiquigent™. In brief, activity based probes **1**-**3, 5-13** were screened against a panel of 41 DUBs at a single concentration (20 µM) for 15 min, followed by treatment with a ubiquitin-rhodamine(110)-glycine substrate for 40 min, according to the protocol below:

Each reaction was performed in duplicate in black 384 well plates (small volume, Greiner 784076) in a final reaction volume of 21 µΙ. Recombinant DUBs were diluted in reaction buffer (40 mM Tris, pH 7.5, 0.005% Tween 20, 0.5 mg/ml BSA, 5 mM DTT) to the equivalent of 0, 0.005, 0.01 , 0.05, 0.1 and 0.5pl/well. Reactions were initiated by the addition of 50 nM of a ubiquitin-rhodamine(110)-glycine substrate as a fluorescence polarisation substrate. Reactions were incubated at room temperature and read every 2 min for 40 min. Readings were performed on a Pherastar Plus (BMG Labtech). λ Excitation 540 nm; λ Emission 590 nm.

7. **Cell death assays for CNPy probe 12 (IMP-2373) and CMK probe 4**
a. LIVE/DEAD ® Viability/Cytotoxicity assay

Cell viability experiments were performed by Pfizer Inc. using LIVE/DEAD ® Viability/Cytotoxicity Kit (Invitrogen Cat# L3224) for mammalian cells, according to the manufacturer’s instructions. In brief, this is a two-color fluorescence cell viability assay that is based on the simultaneous determination of live and dead cells with two probes that measure recognized parameters of cell viability—intracellular esterase activity and plasma membrane integrity. The polyanionic dye calcein is well retained within live cells, producing an intense uniform green fluorescence in live cells (ex/em ~495 nm/~515 nm). EthD-1 enters cells with damaged membranes and undergoes a 40-fold enhancement of fluorescence upon binding to nucleic acids, thereby producing a bright red fluorescence in dead cells (ex/em ~495 nm/~635 nm). A ratio of these two readouts provides the raw data to calculate relative cell viability/cytotoxicity. Briefly, cells were added to compound plates at 10K cells per well and incubated at 37ᵒ 5% CO_2_ for 8h or 24h. Controls added to the plate included DMSO only and cells treated with Lysis buffer (Promega G1821). Reagents were diluted as follows 2 µM Calcein-AM, 4 µM EtHD-1 and 0.5 µg/mL Hoechst 33342 in Phenol red free tissue culture media. Cells were washed with 1x PBS and diluted dye mixture was added to cell and incubated for 30 mins at 37ᵒC 5% CO_2_.  Plates were then imaged on the Opera Phenix spinning disk confocal microscope utilizing a 10x objective (NA 0.75). Images were obtained using 405 nm, 488 nm, and 561 nm lasers for relevant fluorophores. Images were processed and analyzed for relevant features and parameters using Harmony software (Perkin Elmer). Briefly, total live nuclei were identified using Hoechst 33342 for nuclei detection and thresholding based on 488 lasers was used to identify number of Calcein-AM positive cells or 647 laser to identify the number of EthD-1 positive cells. Data was normalized using the 100% dead values in lysis buffer treated cells and 100% live values in DMSO only treat cells. The EC_50_ values for probes **1-14** were evaluated by an 11-point concentration curve (0.0005 ® 50 µM), for 8 h and 24 h incubations in U2OS, U87-MG and T47D cell lines.

b. Sytox Green cell death imaging assay

U2OS, U87-MG or T47D cells were seeded 24 h before treatment in a sterile 96-well plate (Greiner) at a density of 20,000 cells per well to a final volume of 150 µL in McCoy 5a, low glucose DMEM or RPMI-1680 media, respectively, containing 10% FBS. PBS (100 µL) was added to the outer wells.

After 24h, the appropriate wells were treated with 18.6 µL of various concentrations of compound or control (either DMSO only or compound, at final compound concentrations in PBS, 0.1% DMSO, see table S1).

| **Conditions** | Conc (µM) | **No** |  | Conc (µM) | **No** |
| --- | --- | --- | --- | --- | --- |
| Puromycin |  | 1 |  |  |  |
| DMSO |  | 2 |  |  |  |
| CMK probe **4** | 50 | 3 | CNPy probe **12 (IMP-2373)** | 100 | 12 |
| CMK probe **4** | 25 | 4 | CNPy probe **12 (IMP-2373)** | 50 | 13 |
| CMK probe **4** | 12.5 | 5 | CNPy probe **12 (IMP-2373)** | 25 | 14 |
| CMK probe **4** | 6.25 | 6 | CNPy probe **12 (IMP-2373)** | 12.5 | 15 |
| CMK probe **4** | 3.125 | 7 | CNPy probe **12 (IMP-2373)** | 6.25 | 16 |
| CMK probe **4** | 1.5625 | 8 | CNPy probe **12 (IMP-2373)** | 3.125 | 17 |
| CMK probe **4** | 0.78125 | 9 | CNPy probe **12 (IMP-2373)** | 1.5625 | 18 |
| CMK probe **4** | 0.390625 | 10 | CNPy probe **12 (IMP-2373)** | 0.78125 | 19 |
| CMK probe **4** | 0.195313 | 11 | CNPy probe **12 (IMP-2373)** | 0.390625 | 20 |

Table S1.

The PBS compound stocks were pre-prepared in a compound stock plate as follows:

- A 100mM (CMK probe **4** = 50mM) DMSO stock was made up of each probe. 3 µL of each DMSO stock was then diluted with 297 µL of PBS (Final probe conc. = 1mM, 1% DMSO) in a 1.5 mL Eppendorf tube. 150 µL of each 500 µM stock was then transferred into the compound stock plate.
- A 1% DMSO only in PBS solution was also made up added to the appropriate wells for compound dilutions (100 µL DMSO in 9900 µL PBS).
- This 1mM stock was then diluted 2 -fold another 13 times (75 µL into 75 µL 1% DMSO in PBS, final conc in PBS, total volume 75µL)

The Cytotox Green dye reagent (5mM stock, DMSO) was diluted 2000-fold to 2.5 µM (2.5 µL stock into 5000 µL PBS, 15mL falcon). 18.6 µL of this was added to each well using a sterile reservoir to achieve a final concentration of 250 nM, 0.05% DMSO.

Any bubbles were removed from all wells by gently squeezing a wash bottle (containing 70-100% ethanol with the inner straw removed) to blow vapor over the surface of each well (remember to purge upside down first!). The cell plate was the placed into the IncuCyte™ Live-Cell Analysis system and the plate allowed the plate to warm to 37°C for 30 minutes prior to scanning. The cells were then incubated at 37◦C and monitored every 2 h for cell growth and death for 24 hours.

- Live imaging parameters:
  - a. Objective: 10x
  - Channel selection: Phase Contrast and Green Fluorescence (λ_ex_ 440–480 nm, λ_em_ = 504–544 nm)
  - Scan type: Standard (2 images per well)
  - Scan interval: Every 1 hour for 24 hours.

Image analysis

Cell images were analysed with Incucyte 2019B Rev 2 software. Three representative images for DMSO only,and high compound concentration across three time points were used to train the Incucyte software for complete analysis.

The following parameters were used for most experiments, but adjusted according to the cell specific parameters for each assay using the above image set as a guide.

For phase analysis, segmentation adjustment was set to 1 and cleanup parameters were unaltered from the default settings. A minimum area filter of 70 µm^2^ was applied while no filters were added for eccentricity.

For green analysis, Top-Hat segmentation was used with a radius of 100 µM and a threshold of 2 GCU, with edge split on and an edge sensitivity of 0. Cleanup parameters were unchanged and a minimum area filter of 30 µm^2^ and a minimum integrated intensity of 250 were used with other filters unaltered.

Data analysis

After complete analysis of all images, “green integrated intensity per image/phase area per image” ((GCU x µm²/Image) / (µm²/Image), denoted “G_P_”) values were extracted for each well at each time-point and exported to Microsoft Excel. Background subtraction was performed for each well using the value from the initial time point and the data exported to Origin Pro 2019.

Background corrected G_P_ values were grouped into their technical replicates and represented graphically ±SEM plotted against time for each concentration. To generate EC_50_ values over a concentration range, the linear growth phase of each of the replicate curves were taken and linear regression performed on each. The resultant rates were then plotted ±SEM against concentration and fitted using a four-parameter dose-response curve to generate an EC_50_ value for cytotoxicity.

8. **Activity-based DUB labelling in HEK293 cell lysates using TAMRA-Ub-VME and in-gel fluorescence**

HEK293 cell lysates (50 µL, 1mg/mL) resuspended in cold phosphate buffered saline buffer were treated with either DMSO (0.1%) or increasing concentrations of CNP probe **12** (1, 10, 30 and 100 µM, 0.1% DMSO) for 1 hour at ambient temperature. TAMRA-Ub-VME probe (suspended in 5% DMSO, 50mM sodium acetate buffer (pH 4.5) was then added (1 µM final concentration) and lysates incubated for 1 hour at ambient temperature. SDS-PAGE and in-gel fluorescence scanning were then performed as described previously in section 4c.

9. **Concentration dependent chemical proteomic profiling of CNPy probe 12 in live cells**

U2OS, T47D and U87-MG cells were seeded in 10cm dishes at 4 x 10^4^ cells/mL and incubated overnight at 37 °C. The following day, the increasing concentrations of CNPy probe **12** (3.125, 6.25, 12.5, 25. 50, 100 µM, each 10 µL, 0.1% final DMSO concentration) were added to the media and incubated with cells for 4 h at 37 °C (final concentration 0.1% (v/v) DMSO). The media was removed and cells were washed with PBS twice (5 mL), resuspended in 1 mL of PBS, scraped and transferred to an eppendorf tube. The cells were then snap frozen using liquid nitrogen and stored for later use.

Cells pellets were resuspended in 500 µL of cold HR buffer (50 mM Tris–HCl (pH 7.4), 5 mM MgCl2, 250 mM sucrose, 0.5% CHAPS (3-[(3-Cholamidopropyl)dimethylammonio]-1-propanesulfonate), 0.1% NP40, 1 mM DTT (added fresh from a 1 M stock solution before use), 1x EDTA-free protease inhibitor (Roche). Each lysate was sonicated for 1 minute, and the samples centrifuges at 5,000 x g at 4 ˚C for 5 minutes to pellet insoluble cellular debris. The supernatant was collected and protein concentration determined using the DC Protein Assay (Bio-Rad) as per manufacturer’s instructions, and normalised to lowest concentrated sample (e.g. 1 mg/mL, using lysis buffer). A total of 250ug protein per condition was aliquoted for Ub-ABP in-cell labelling (see section 7). The remaining sample (~300-600ug) was precipitated, clicked and precipitated again for the ABPP probe profiling experiment using the protocol described in section 4b. The pellet was resuspended as previously described in section 5b.

Neutravidin biotin enrichment and on-bead digestion, isobaric TMT labelling and high pH reverse phase fractionation, LC-MS/MS analysis, database searching and proteomics data analysis were all performed as previously described in section 5.

10. **Activity-based DUB labelling in live cells using HA-Ub-PA, HA enrichment and proteomics**

Ubiquitin activity based probe labelling experiments were performed with hemagglutinin (HA) tagged ubiquitin propargyl amide (HA-Ub-PA, obtained from UbiQ Bio, catalog number: UbiQ-078), according to the protocol published by Pinto-Fernández *et al*.^1^

a. Preparation of HA-Ub-PA stock

A 50 ug stock of HA-Ub-PA was dissolved in 10 µL of DMSO. To this solution of DMSO was then slowly added 190 µL of 50 mM sodium acetate buffer (pH 4.5) to achieve a final Ub ABP probe concentration of 0.25mg/mL (25µM), 5% DMSO.

b. HA-Ub-PA labelling

The 250ug protein samples (1mg/mL, 250µL), as prepared in section 9a, were then treated with HA-Ub-PA (5µL of (0.25mg/mL) 25µM stock in 50mM NaOAc pH 4.5) followed by the addition of 10µL of 50mM NaOH (to neutralise pH). Samples were then vortexed, spun briefly, and incubated at 37◦C for 45 min (265 μL total volume).

The reaction was quenched by the addition of SDS to 0.4% (30 µL of a 4% stock for a total volume of 300 µL) and NP-40 to 0.5% (33 µL of a 4% stock for a total volume of 330 µL), and samples were diluted to 1ml, 0.25 mg/ml, by the addition of 670 µL of NP-40 lysis buffer [pH 7.4, 50mM Tris, 0.5% (v/v) NP-40, 150mM NaCl, and 20mM MgCl_2_].

c. HA-enrichment and denaturing elution

To bind and pull down DUB– ABP complexes, 50 µL of anti-HA-Agarose slurry (Sigma, catalog number: A2095-1ML), previously washed four times with NP-40 lysis buffer, was added to the samples and incubated on a rotator overnight at 4◦C. After a first centrifugation step (2,000 g, 4◦C, 1min), beads were washed four times with 1000 µL of NP-40 lysis buffer. Protein complexes were eluted by boiling beads in 110 µL of 2× SDS Laemmli sample buffer (prepared from 2-fold dilution of 4x sample loading buffer with beta-mercaptoethanol) for 10 min at 95◦C.

d. Reduction/Alkylation and precipitation of eluted samples

DUB-probe immunoprecipitated sample eluates were diluted to 175 µL with proteomics grade water and reduced with 5 µL of DTT (200mM in 0.1M Tris, pH 7.8) for 30min at 37◦C. Samples were alkylated with 20 µL of iodoacetamide (100mM in 0.1M Tris, pH 7.8) for 15min at room temperature (protected from light), followed by protein precipitation using a double methanol/chloroform extraction method.^2^ Protein samples were treated with 600 µL of methanol, 150 µL of chloroform, and 450 µL of water, followed by vigorous vortexing. Samples were centrifuged at 17,000 g for 3 min, and the resultant upper aqueous phase was removed. Proteins were pelleted following the addition of 450 µL of methanol and centrifugation at 17,000 g for 6 min. The supernatant was removed, and the extraction process was repeated. Following the second extraction process, precipitated proteins were re- suspended in 25 µL of 6M urea and diluted to <1M urea with 125 µL of 50mM HEPES pH 8.0 buffer. Protein digestion was carried out by adding trypsin to a ratio 1:100 (1 ug of trypsin per sample), rocking at 1200 rpm and 37◦C overnight.

The subsequent isobaric TMT labelling and high pH reverse phase fractionation, LC-MS/MS analysis, database searching were all performed as previously described in section 5.

e. Proteomics data analysis

Data analysis was performed in Perseus version 1.6.6.0. Reporter intensity corrected values were loaded into the matrix. Data was filtered by removing rows based on “reverse”, and “potential contaminant” columns. Data were log2 transformed and filtered by row, retaining those that had 2 valid values in each triplicate condition. To account for variance in protein abundance across different sample, the median of each channel was subtracted from each protein, or each channels protein abundance was normalised to the abundance of ubiquitin protein in the Ub-ABP only treated sample. The log2 fold competition for each probe against Ub-ABP only was determined by subtracting the Ub-ABP only control value from each of the different probe concentration/competition conditions.

### 11. Concentration dependent chemical proteomic and Ub-activity-based profiling of CNPy probe 12 enantiomers 12a and 12b in T47D cells

T47D cells were seeded in 10cm dishes at 4 x 10^4^ cells/mL and incubated overnight at 37 °C. The following day, the either DMSO CNPy probe **12** (racemate), enantiomer **12a** or enantiomer **12b** (0.5, 5 and 50 µM, each 10 µL, 0.1% final DMSO concentration) were added to the media and incubated with cells for 4 h at 37 °C (final concentration 0.1% (v/v) DMSO). The media was removed and cells were washed with PBS twice (5 mL), resuspended in 1 mL of PBS, scraped and transferred to an eppendorf tube. The cells were then snap frozen using liquid nitrogen and stored for later use.

ABPP and Ub-activity-based profiling, sample processing and data analysis was carried out as described in sections 5 and 10, respectively.

### 12. DUB inhibitor profiling by click chemistry, biotin pull-down and western blotting

#### a.  Live cell inhibitor/probe incubation

For each experiment, sterile treated 6-well plate were seeded with either U87-MG or T47D cells in low glucose DMEM or RPMI-1680 media, respectively, containing 10% FBS, and incubated at 5% CO_2_ and 37◦C overnight. On the other day, plates were treated with either DMSO vehicle (0.1% (v/v)) or indicated concentration of UCHL1/USP30 inhibitor. After incubating the cells with inhibitors at 37 °C for 1 h, 10 μM CNPy probe **12** probe/DMSO was added and incubated for another 1 h at 37 °C.

The cells were relieved of media, washed twice with PBS (1mL), and then lysed with lysis buffer (100 µL (1% (v/v) Triton X-100, 0.1% (w/v) sodium dodecyl sulfate (SDS), EDTA-free complete protease inhibitor cocktail (1×, Roche) in PBS on ice for 10 min. The lysates were scraped and transferred to corresponding Lo-Bind Eppendorfs. Each lysate was sonicated for 1 minute, and the samples centrifuges at 5,000 x g at 4 ˚C for 5 minutes to pellet insoluble cellular debris. The supernatant was collected, and protein concentration determined using the DC Protein Assay (Bio-Rad) as per manufacturer’s instructions, and normalised to lowest concentrated sample (e.g. 1 mg/mL, using lysis buffer).

#### b. Click chemistry and precipitation

Before performing Click chemistry, 20 μL of each sample was aliquoted as TL (Total Lysate) samples and mixed with 4x Laemmli sample buffer (Bio-RAD, catalogue number: 1610747) containing 10% (v/v) β-mercaptoethanol (Sigma, catalogue number: 444203) with boiling at 95 °C for 5 min. The left samples were carried out click chemistry and precipitation as described in section 4b.

The pellet was resuspended by sonicating and completely dissolving in 1% (w/v) SDS in HEPES 50 mM pH 8.0, before being made up to 1 mg/mL protein with HEPES 50 mM pH 8.0 to achieve a final concentration of 0.2% SDS. Samples were centrifuged for 5 min at maximum speed at room temperature to make sure no pellet formed and protein was fully dissolved.

#### c. Neutravidin biotin enrichment and western blot analysis

Neutravidin biotin enrichment was performed as described in section 5c. After beads wash, the samples are eluted with 2x Laemmli sample buffer containing 10% (v/v) β-mercaptoethanol at 95 °C for 10 min. Proteins were separated by precast SDS-PAGE gel (12%, Bio-RAD, catalogue number:4561046) and transferred to nitrocellulose membranes (Amersham™ Protran® , GE Healthcare) by wet-tank transfer (Bio-RAD) in Tris-Glycine transfer buffer supplemented with 20% (v/v) MeOH for 1 h at 100 V. Membranes were blocked in 5% (w/v) slimmed milk in Tris-buffer (50 mM Tris pH 7.4, 150 mM NaCl) containing 0.01% (v/v) Tween-20 (TBST) for 1 h before incubation with the following primary antibody in the corresponding buffer overnight at 4°C:

UCHL1, ProteinTech, catalogue number: 14730-1-AP, 1:2000 in 5% (w/v) slimmed milk in TBST

USP30, Invitrogen, catalogue number: PA5-53523, 1:1000 in 5% (w/v) slimmed milk in TBST

β-actin, Abcam, catalogue number: AB8227, 1:5000 in 5% (w/v) slimmed milk in TBST

GAPDH, Abcam, catalogue number: AB9485, 1:2500 in 5% (w/v) BSA in TBST

The membrane was washed three times with 10 mL TBST for 5 min and incubated with the corresponding HRP-conjugated secondary antibody (α-mouse-HRP or α-rabbit-HRP) in 5% (w/v) slimmed milk in TBST for 1 h at room temperature. After three washes with 10 mL TBST (10 min each), the membrane was incubated with HRP substrate (Luminata Crescendo, Millipore) and the chemiluminescence signal captured with an ImageQuant™ LAS 4000 imager.

### 13.  ABPP and whole proteomic profiling of MYC deregulated cancer cell line P493-6 using CNPy probe 12

#### a. Cell Culture and probe incubation

P493-6 Cells were grown in DMEM high glucose with 10% FBS, with added non-essential amino acids (NEAA, 1: 100, v/v), sodium pyruvate (NaPyr, 1: 100, v/v), HEPES (1: 100, v/v) and 2 μL β-mercaptoethanol per 500 mL of media at 5% CO_2_ and 37ºC.

For each experiment, on day zero (0 h), 10 mL cells suspension at a concentration of 1,000,000 cells/mL were seeded in the respective media to induce the respective MYC, high MYC: culturing media; low MYC: culturing media with 0.1 μg/mL of doxycycline and 1 μM of β-estradiol. On day three (71h), both high and low MYC cells were treated with either DMSO vehicle (0.1% (v/v)) or of CNPy probe **12** (25 µM) and incubated at 37 °C for 1h, to create four treatment groups (n =3, Table 2.). All compound treatments were performed with a 1000× DMSO stock of each probe which was added directly to flask with gentle mixing.

| **MYC Level** | **Conditions** | **N** | **Cell** | **TMT channel** | **TMT set** |
| --- | --- | --- | --- | --- | --- |
| **High** | **DMSO** | 1 | P-493-6 | 126 | 1 |
|  | **CNPy 12** | 1 | P-493-6 | 127 |  |
| **Low** | **DMSO** | 1 | P-493-6 | 128 |  |
|  | **CNPy 12** | 1 | P-493-6 | 129 |  |
| **High** | **DMSO** | 2 | P-493-6 | 126 | 2 |
|  | **CNPy 12** | 2 | P-493-6 | 127 |  |
| **Low** | **DMSO** | 2 | P-493-6 | 128 |  |
|  | **CNPy 12** | 2 | P-493-6 | 129 |  |
| **High** | **DMSO** | 3 | P-493-6 | 126 | 3 |
|  | **CNPy 12** | 3 | P-493-6 | 127 |  |
| **Low** | **DMSO** | 3 | P-493-6 | 128 |  |
|  | **CNPy 12** | 3 | P-493-6 | 129 |  |

**Table 2.**

The cells were relieved of media, centrifuged, washed twice with PBS (5mL), and then lysed with lysis buffer (500 µL (1% (v/v) Triton X-100, 0.1% (w/v) sodium dodecyl sulfate (SDS), EDTA-free complete protease inhibitor cocktail (1×, Roche) in PBS on ice for 10 min. The lysates were scraped and transferred to corresponding Lo-Bind Eppendorfs. Each lysate was sonicated for 1 minute, and the samples centrifuges at 5,000 x g at 4 ˚C for 5 minutes to pellet insoluble cellular debris. The supernatant was collected, and protein concentration determined using the DC Protein Assay (Bio-Rad) as per manufacturer’s instructions, and normalised to lowest concentrated sample (e.g. 1 mg/mL, using lysis buffer).

#### b. Western blotting analysis for MYC level

20 µL samples were aliquoted and mixed with 4x Laemmli sample buffer containing 10% (v/v) β-mercaptoethanol with boiling at 95 °C for 5 min. Western blotting analysis was performed as previously described in section 11c. Primary antibody information is as following:

c-MYC, Abcam, catalogue number: ab32072, 1:1000 in 5% (w/v) skimmed milk in TBST

Vinculin, Abcam, catalogue number: ab129002, 1:1000 in 5% (w/v) skimmed milk in TBST

#### c. Chemical Proteomic profiling of probes in live cells

500 µL samples (1 µg/µL protein concentration) were aliquoted for chemical proteomic profiling as previously described in section 5b-5f.

#### d. Whole proteome analysis of probe in live cells

100 µg samples (100 µL, 1 µg/µL protein concentration) were aliquoted, and subsequently reduced and alkylated with 5 mM TCEP and 10 mM chloroacetamide with moderate shaking for 45 min at r.t.. The resulting proteins were precipitated by adding H_2_O (1 vol), MeOH (2 vol) and CHCl_3_ (0.5 vol), vortexing briefly then centrifuging at 17,000 × g for 5 min. The top H_2_O/MeOH layer was discarded and the middle layer of protein pellet and lower CHCl_3_ layer retained. The suspension was washed with MeOH (500 µL), and sonicated to break up the pellet. The proteins were pelleted by centrifugation at 10,000–17000 × g for 5–10 min or until a compact pellet was formed. The protein pellet was washed once more with MeOH (500 µL). The MeOH was decanted, and the pellet air-dried for 1 min. The precipitated proteins were re-suspended in 20 µL of 6M urea and diluted to <1M urea with 100 µL of 50mM HEPES pH 8.0 buffer. Protein digestion was carried out by adding trypsin to a ratio 1:100 (1 ug of trypsin per sample), rocking at 1200 rpm and 37◦C overnight.

#### e. Isobaric TMT labelling

Sample replicates were arranged for labelling in three TMT sets as described in Table 2. Upon checking that the pH of each sample was ~8. approximately 10 ug of peptide sample was labelled with 1/10^th^ of an 0.8 mg vial of the appropriate TMT10plex™ Isobaric Mass Tag Labelling Reagent (Thermo Scientific) dissolved in acetonitrile (40 µL) with moderate shaking for 2 h at rt TMT-labelling was quenched by the addition of 1 µL of 5% (w/v) hydroxylamine and the samples from each TMT set were combined to form a “multiplex” solution. Trifluoroacetic acid was added to achieve a 1% v/v solution and these samples were evaporated to dryness.

The subsequent and high pH reverse phase fractionation, LC-MS/MS analysis, and database searching were all performed as previously described in section 5.

#### f. Proteomics data analysis

Data analysis was performed in Perseus version 1.6.6.0. Reporter intensity corrected values were loaded into the matrix. Data was filtered by removing rows based on “reverse”, and “potential contaminant” columns. Data were log2 transformed and filtered by row, retaining those that had at least 2 valid values in each triplicate set of conditions. To account for variance in protein abundance across different samples, the median of each channel was subtracted from each protein. Multiple TMT data sets were normalized by subtracting the mean of each row within each TMT “plex”.

For the whole proteome data, a student’s t-test (FDR = 0.05, S0 = 0.1) was performed to determine statistically significance changes in DUB abundance between untreated medium and high MYC cell lines (S11A-B).

For the ABPP data, the mean of the replicates for each protein within control group (DMSO only) was calculated, and then this value was subtracted from each value for the treated replicates from the same cell line (high or medium MYC) to determine the relative CNPy probe **12** engagement for each protein within that particular cell line (as represented in volcano plots Fig S11C-D). Then, a students t-test (FDR = 0.05, S0 = 0.1) was performed on this normalised data to determine the statistically significance changes in CNPy probe **12** engagement across the two cell lines (medium and high MYC) (S11E-F). Changes in DUB abundance and activity are highlighted in blue on these graphs to show their relative Log_2_fold change and statistical significance compared with the background.
