## Supplementary information 3 - 1H NMR spectra of chemical probes for "Electrophile scanning by chemical proteomics reveals a potent pan-active DUB probe for investigation of deubiquitinase activity in live cells"

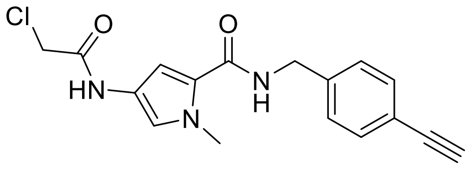


Compound **1**

$^{1}$H NMR (400 MHz, $CDCl_3$)

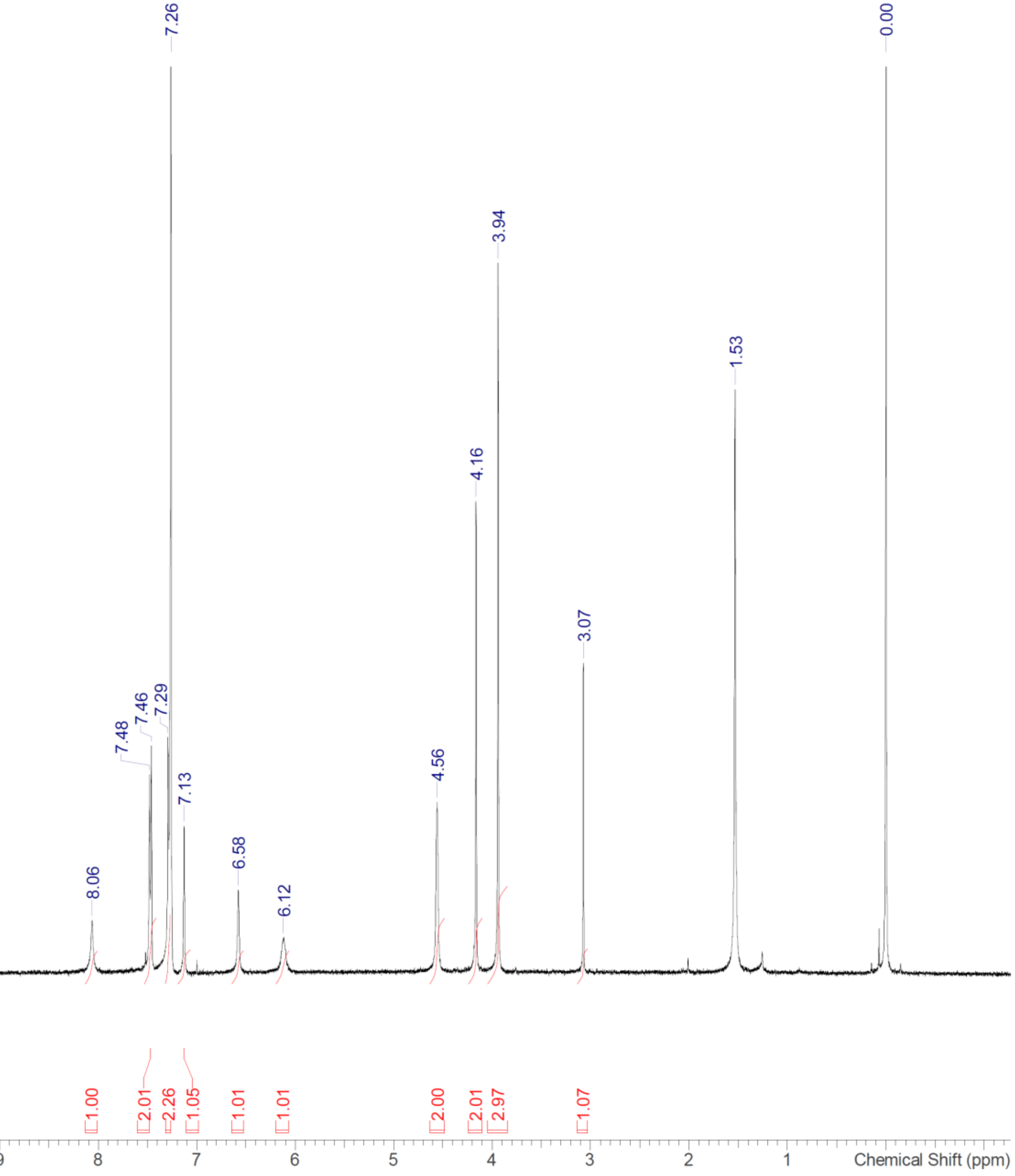

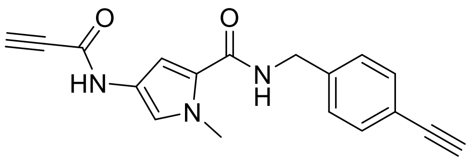


Compound **2**

$^{1}$H NMR (400 MHz, DMSO-$d_6$)

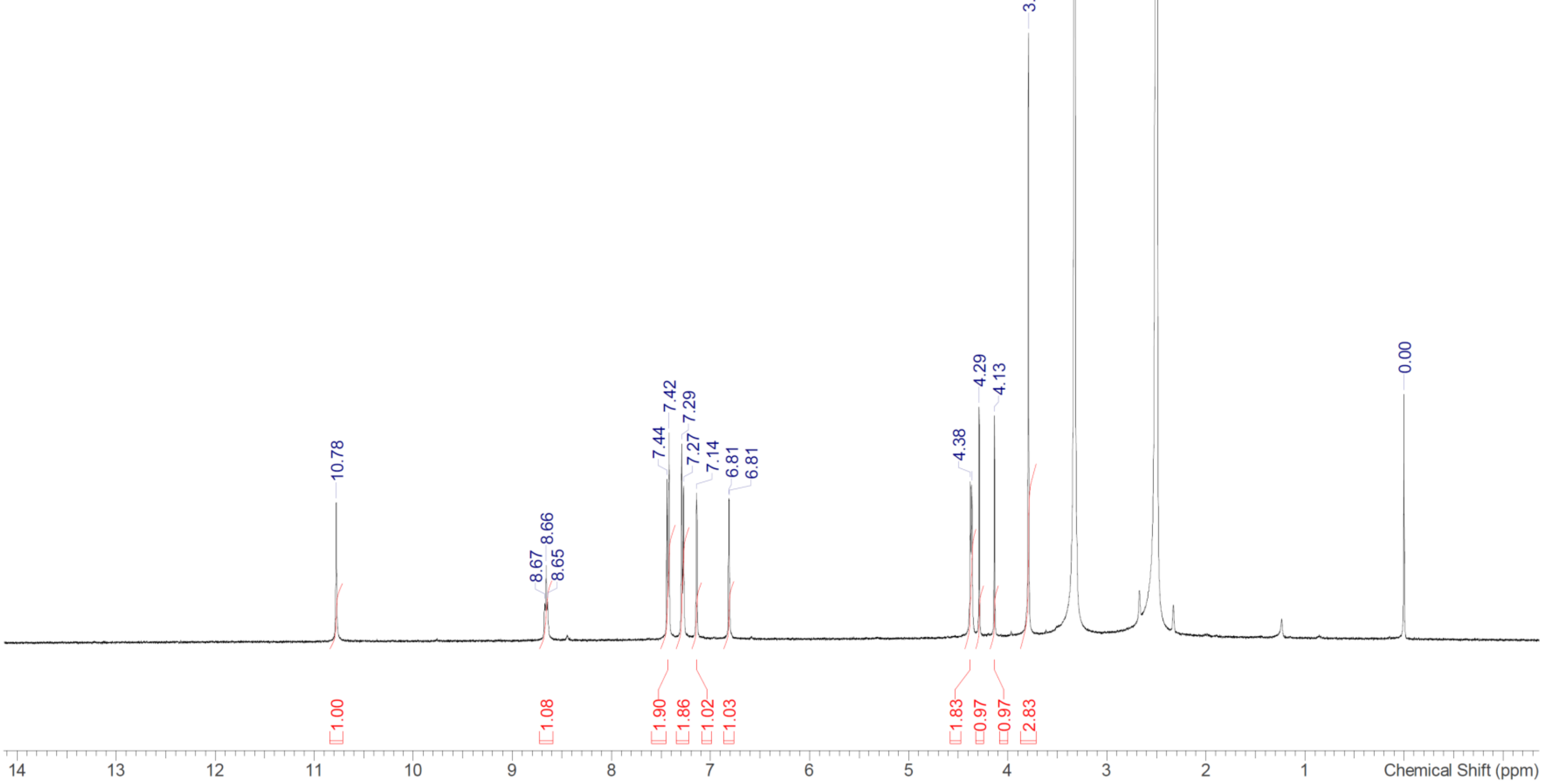

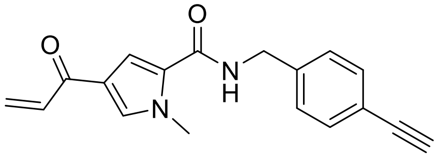

Compound **3**

$^1$H NMR (400 MHz, DMSO-$d_6$)

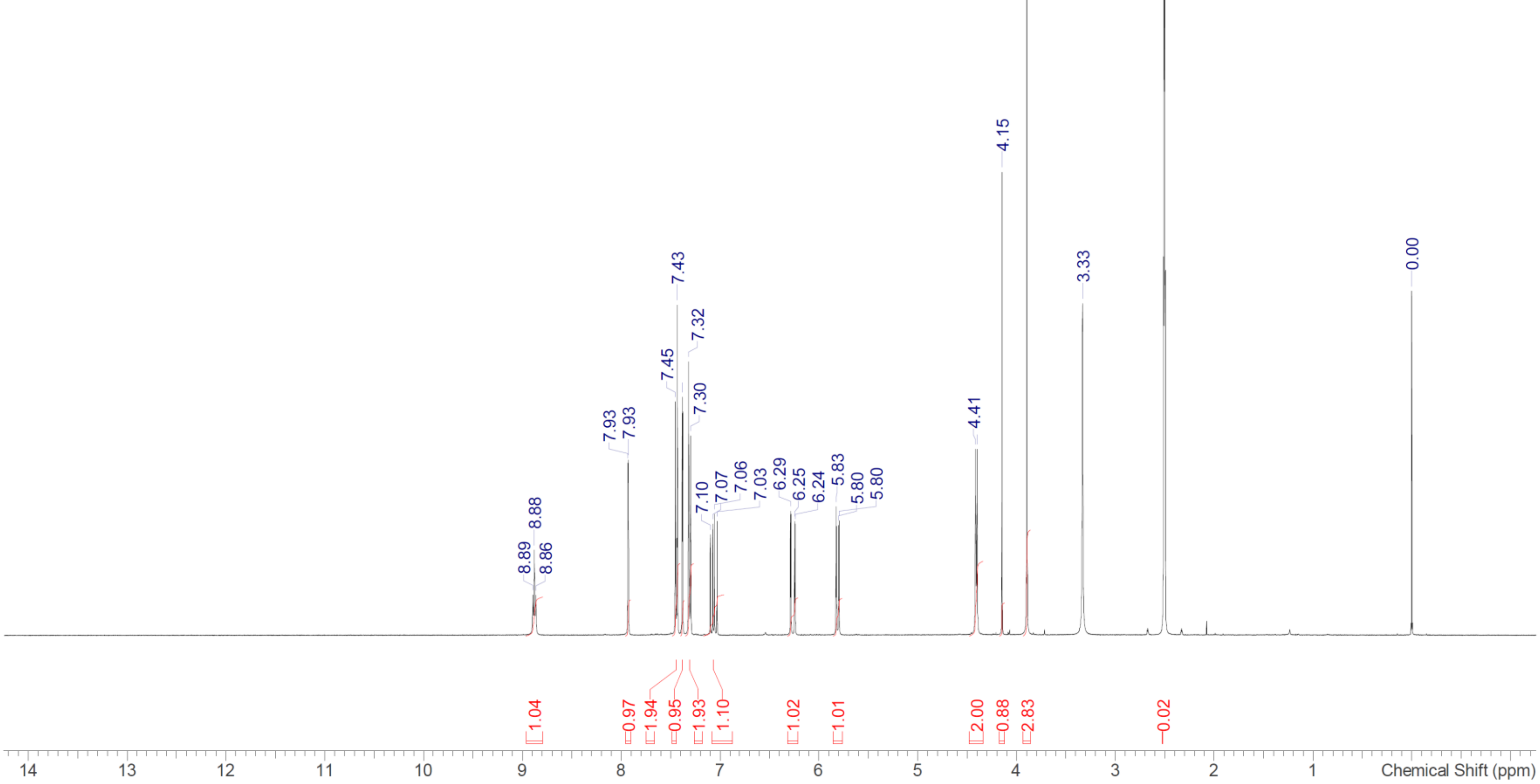

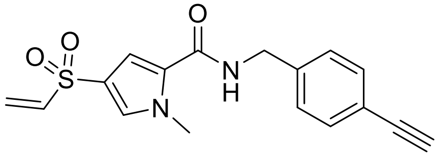

Compound **5**

$^1$H NMR (400 MHz, DMSO-$d_6$)

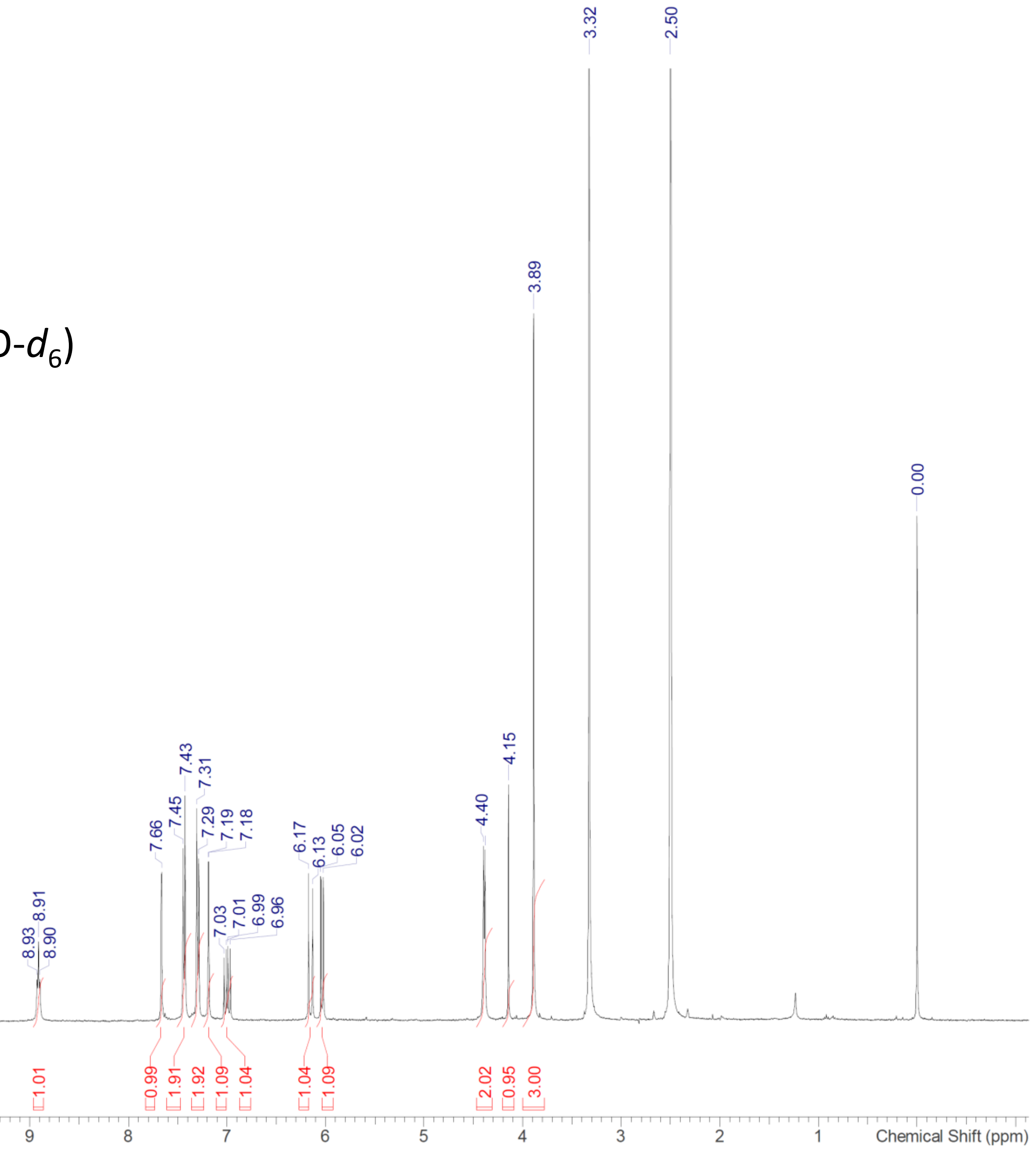

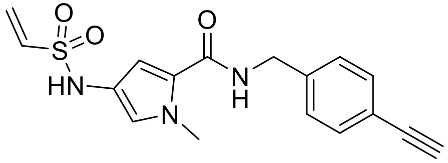

Compound **6**

$^1$H NMR (400 MHz, DMSO-$d_6$)

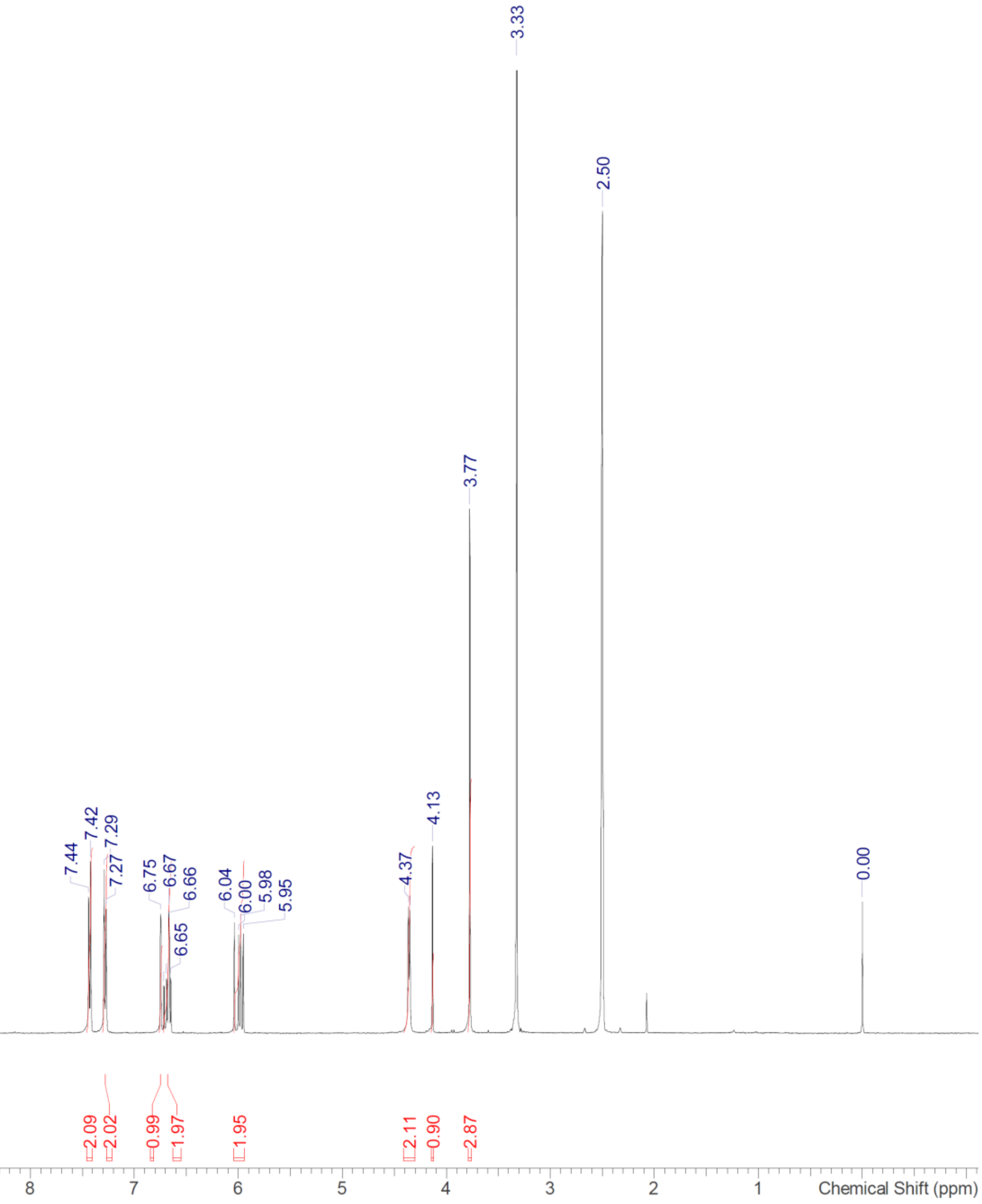

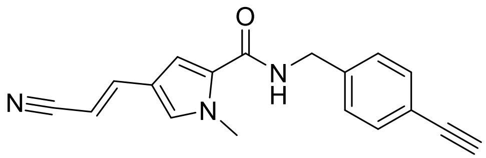

Compound **7**

$^{1}H$ NMR (400 MHz, DMSO-$d_6$)

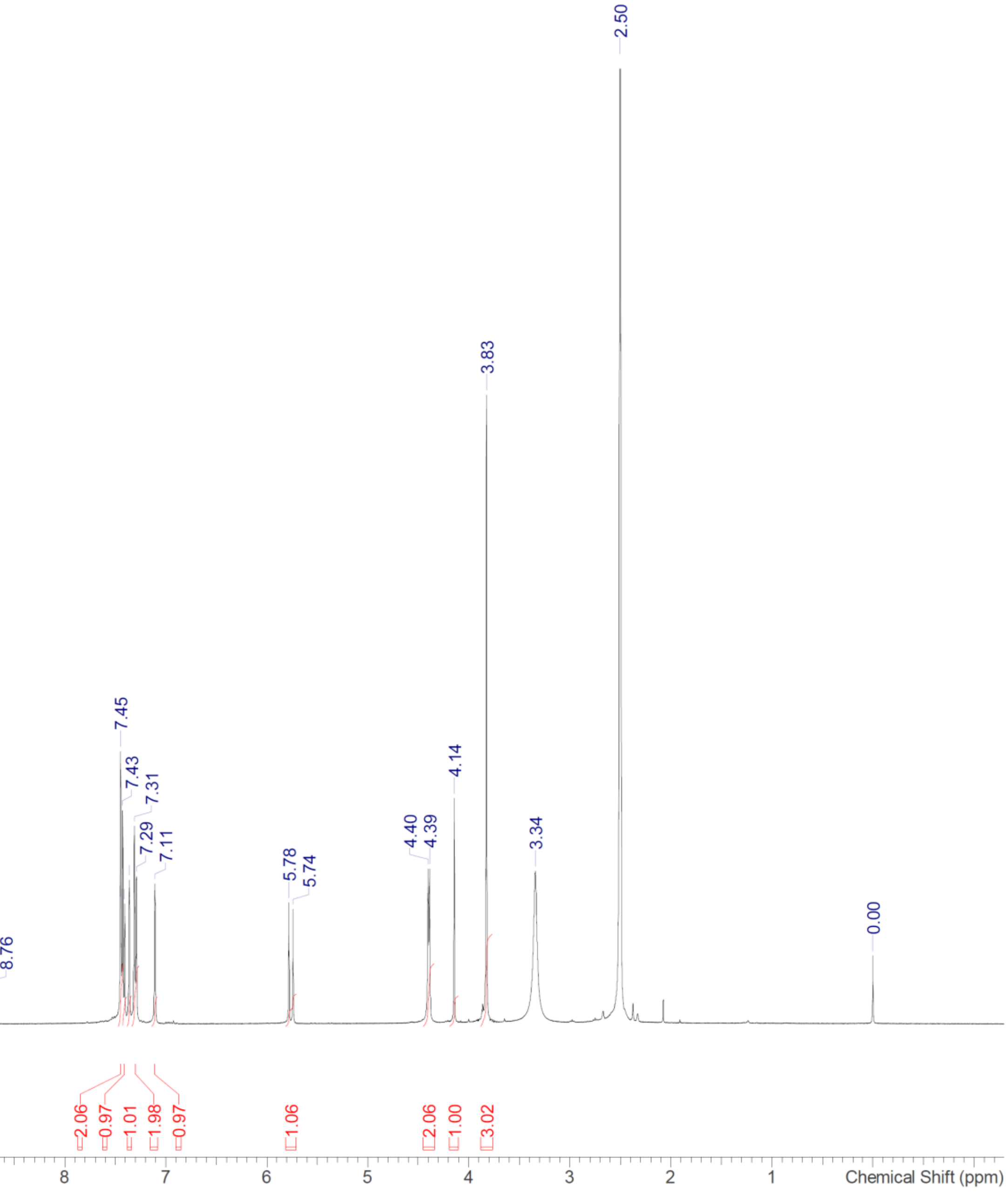

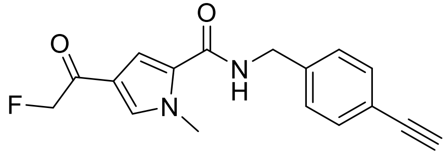

Compound **8**

$^{1}$H NMR (400 MHz, DMSO-$d_6$)

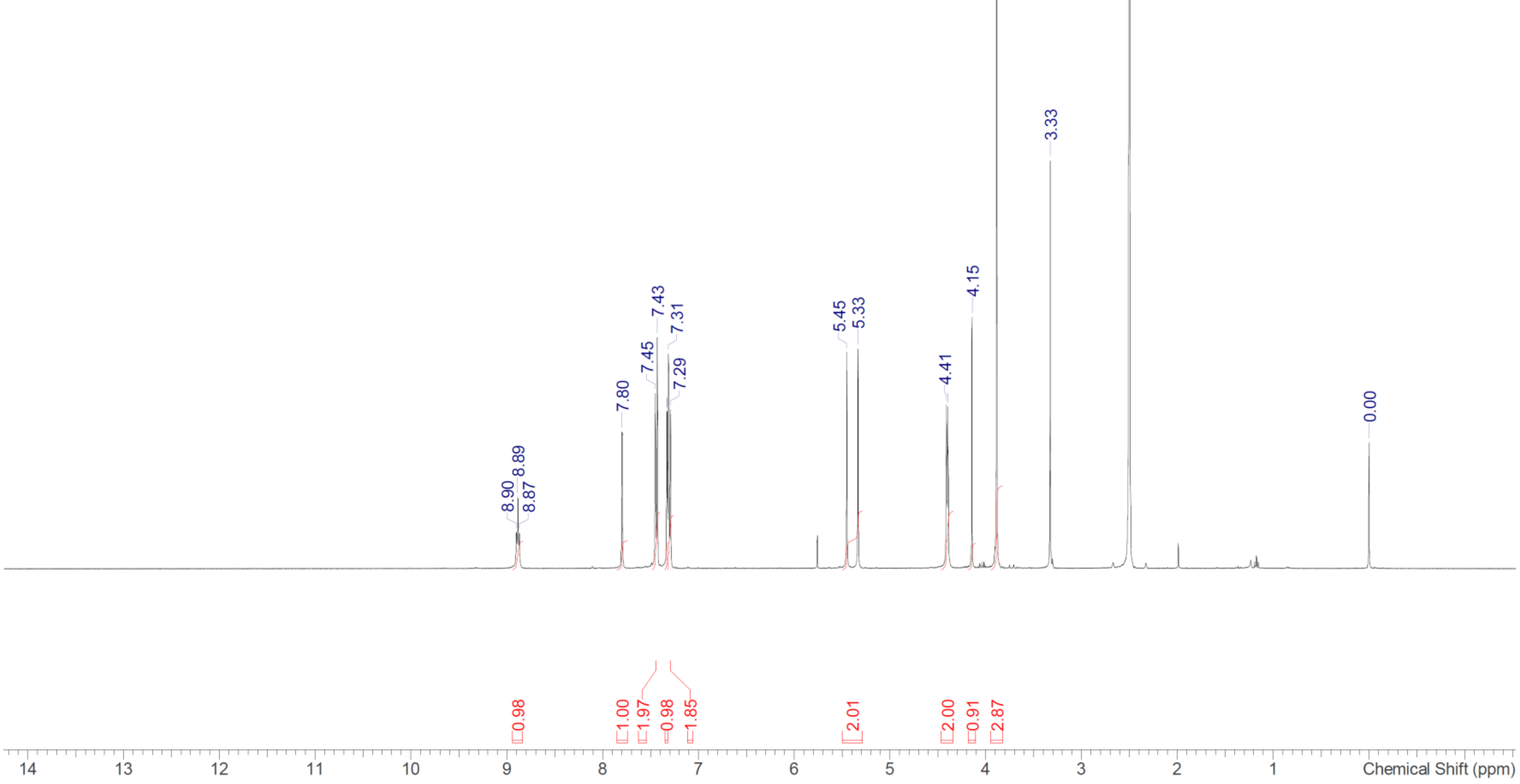

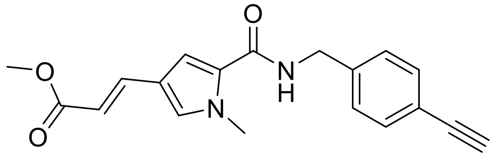

Compound **9**

$^1$H NMR (400 MHz, DMSO-$d_6$)

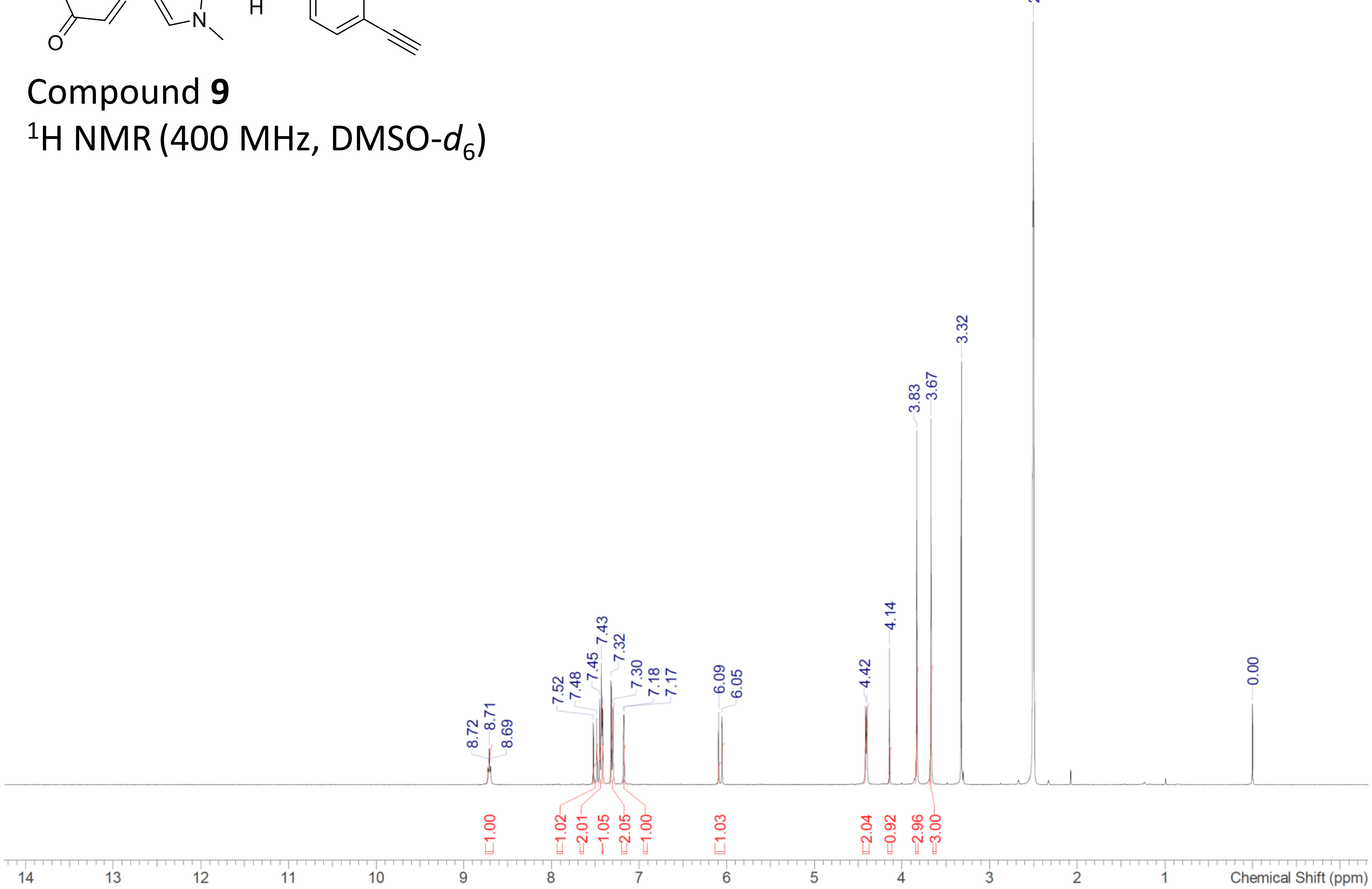

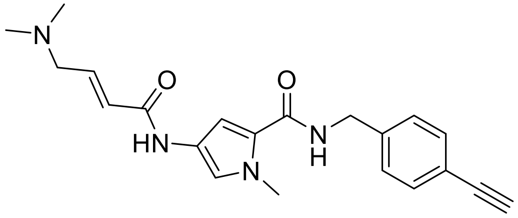

Compound **10**

$^{1}$H NMR (400 MHz, DMSO-$d_6$)

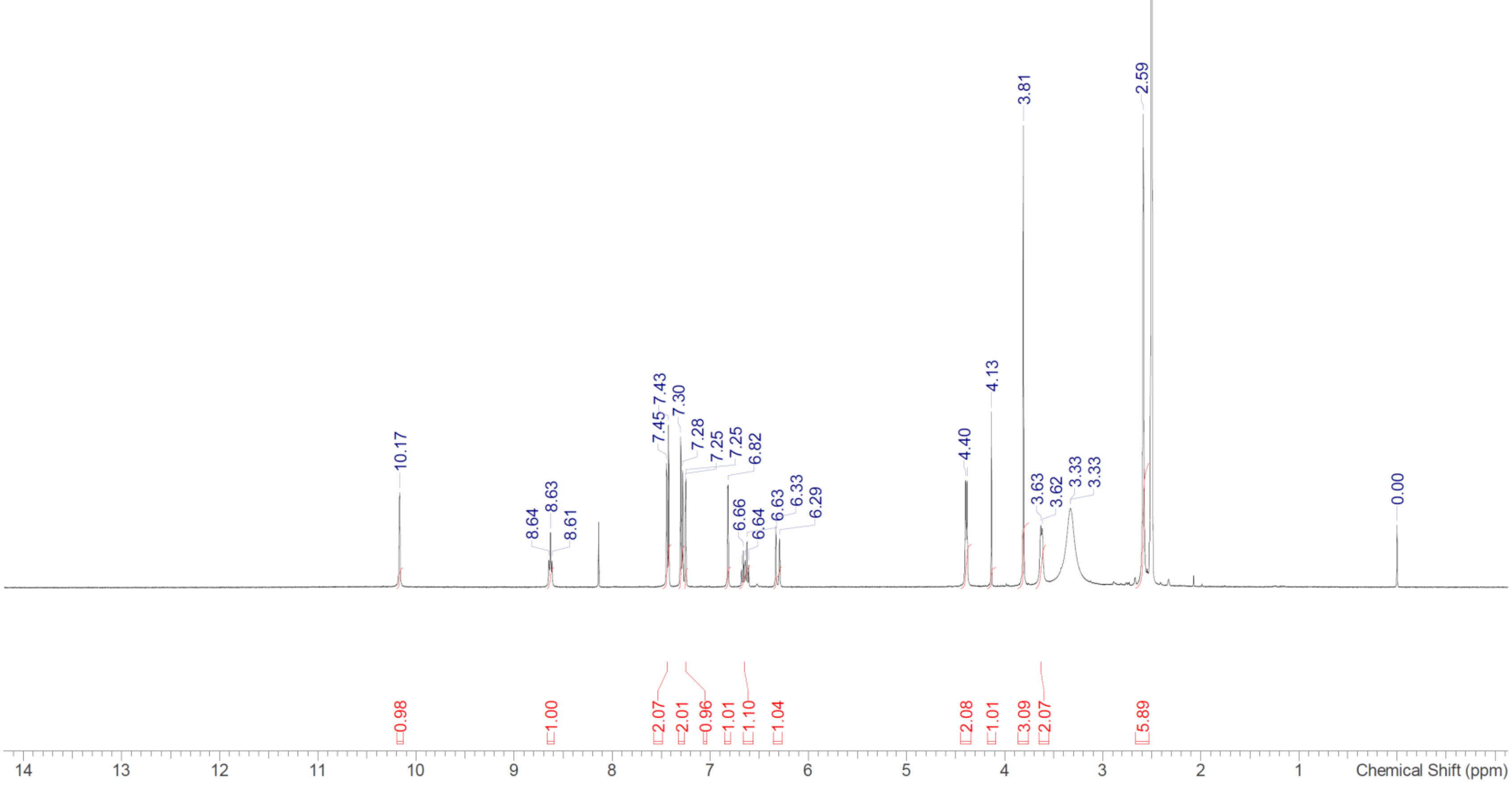

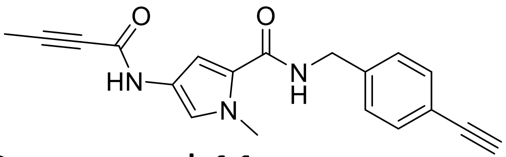

Compound **11**

$^{1}$H NMR (400 MHz, DMSO-$d_6$)

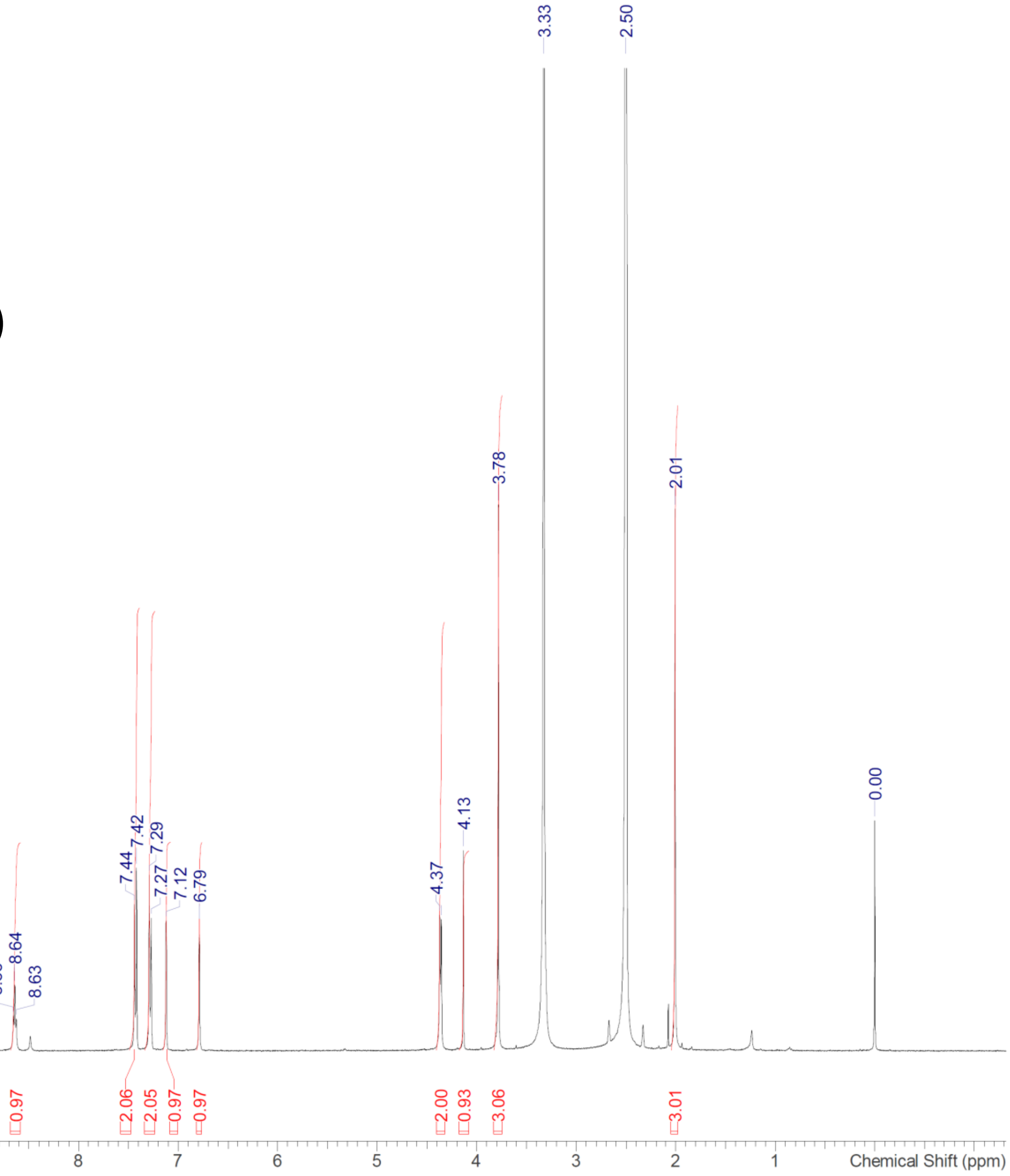

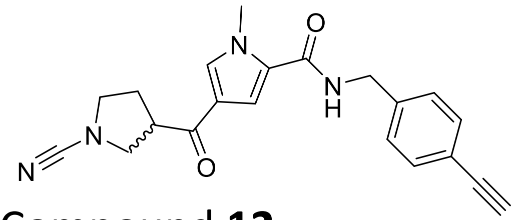


Compound **12**

$^1$H NMR (400 MHz, DMSO-$d_6$)

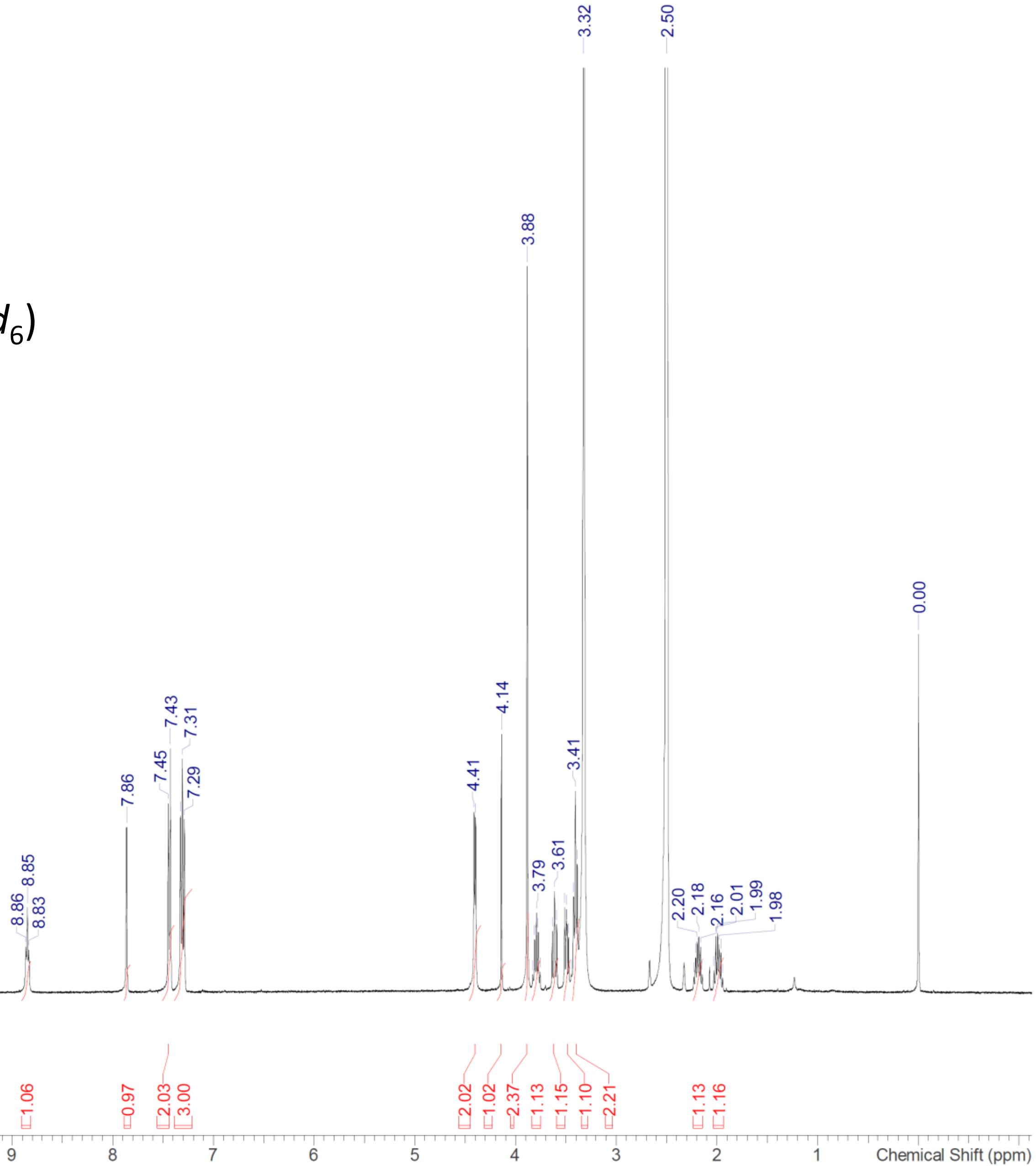

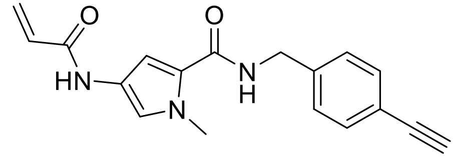


Compound **13**

$^{1}$H NMR (400 MHz, DMSO-$d_6$)

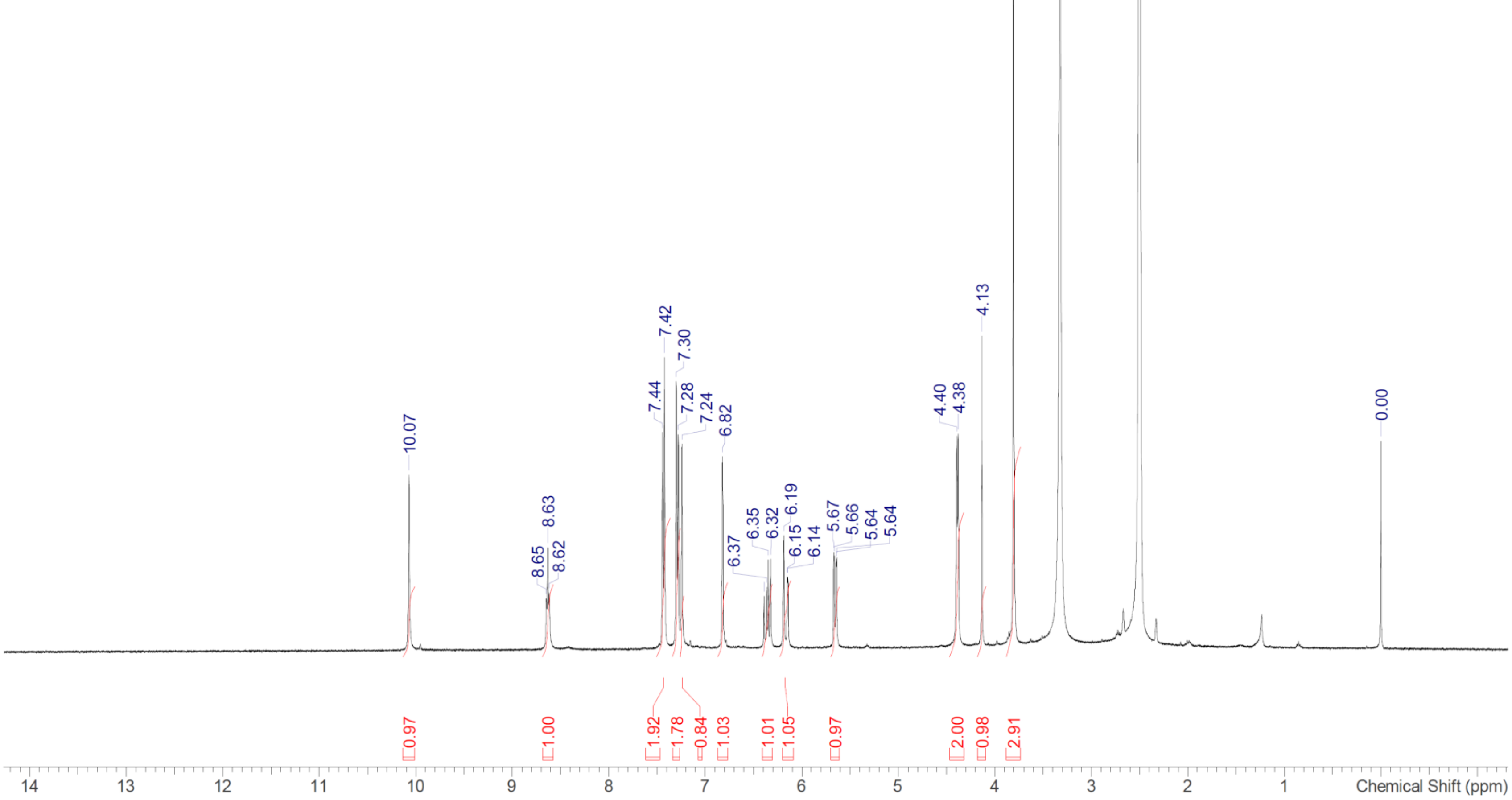

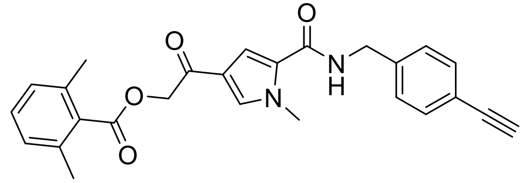

Compound **14**

$^1H$ NMR (400 MHz, DMSO-$d_6$)

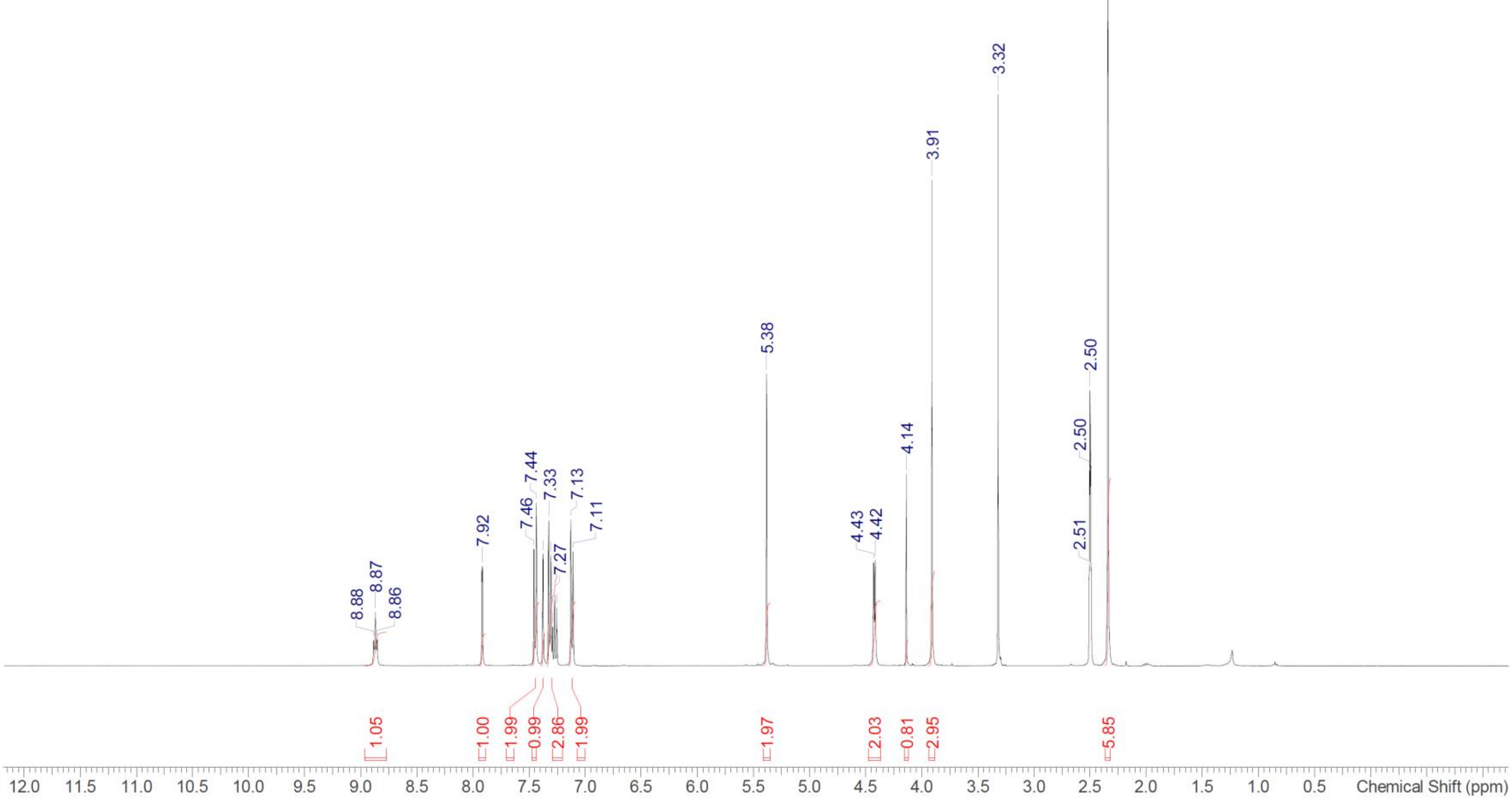
